# Anti-PD-1/PD-L1 Therapy Triggers Cognitive Deficits and Anxiety-Like Behaviors Through Tumor-Initiated Neuroinflammatory Niches in Male Mice

**DOI:** 10.1101/2025.09.03.673981

**Authors:** C. Nicola, M. Pedard, M. Dubois, L. Desrues, P. Neveu, G. Riou, I Johnston, K.P. Dembele, P. Lecras, D. Vaudry, S. Adriouch, F. Joly, P. Hilber, O. Wurtz, H. Castel

**Affiliations:** Univ Rouen Normandie, Inserm, Normandie Univ, CBG UMR 1245, Rouen, France; Institute for research and biomedical innovation (IRIB), 76000 Rouen, France; Services Unit PLATON, Cancer and Cognition Platform, University of Caen Normandy, 14000 Caen, France; Univ Rouen Normandie, Inserm, UMR1234, F-76000, Rouen, France; School of Psychology, The University of Sydney, Sydney, Australia; Univ Rouen Normandie, Inserm, US, HERACLES, 76000 Rouen, France ANTICIPE U1086 INSERM-UCN, Equipe Labellisée Ligue Contre le Cancer, Centre François Baclesse, Normandie Université UNICAEN, 14000 Caen, France; Clinical Research Department, Centre François Baclesse, 14000, Caen, France; Medical oncology department, CHU de Caen, 14000 Caen, France

**Author notes:** Corresponding author. Dr Hélène Castel, 1 Univ Rouen Normandie, Inserm, Normandie Univ, CBG UMR 1245, Genetics, Biology and Plasticity of Brain Tumors – NeuroGlio 25 Rue Tesnière, CURIB, 76821 Mont-Saint-Aignan, France.

## Abstract

Checkpoint inhibitors are promising immunotherapy to treat cancer patients, but their cognitive impact has not been evaluated despite several neurological adverse events. We studied the impact of immune desert or inflamed cancers when combined with immune checkpoint inhibitors (ICI) anti-PD-1/anti-PD-L1 on mouse behaviors and brain immune cells infiltration/homeostasis, and neuroinflammation in male mice. We showed that systemic inflammation, brain-barriers permeability accompanying meningeal infiltration of peripheral macrophages and neuroinflammation as well as deficits in cognition or emotional reactivity, depending on immuno-inflammatory or immune-desert cancer type. Combined with cancers, anti-PD-1 and PD-L1 treatments exacerbated the decline in executive functions and hippocampal vascular inflammation. PD-L1 specifically relayed the infiltration of the Tγδ lymphocytes subpopulation in choroid plexus and leptomeninges implicated, whose systemic neutralization counteracted anti-PDL1-induced cognitive deficits and anxiety in mice bearing immune-inflamed cancer. Our findings highlight new systemic biomarkers of cold or hot cancer, treated with anti-PD-1/anti-PDL-1, and associated with cognitive and emotional alterations in mice; guiding ways of intervention to secure the cancer curation and improve patient’s quality of life under ICI treatment.

**Competing Interest Statement:** The authors have declared no competing interest.

**One Sentence Summary:** Impact of cancer and checkpoint inhibitors on cognitive functions

## Introduction

Advances in clinical oncology over the last decade have significantly improved the long-term survival of some population of cancer patients but whose quality of life (QoL), return to work or autonomy can be severely compromised by various combined or consecutive treatments leading to acute and late toxicities including cognitive impairment^1–5^. Cancer-Related Cognitive Impairment or CRCI^6,7^ primarily related to chemotherapy, includes cancer patients’ complaints about their perceived cognitive ability and objective measured of deficits in short-term and working memory, attention, executive functions, and/or information processing speed^8–13^. These symptoms may persist in nearly 35% patients years after the end of the chemotherapy period^8–13^, and can be worsened by a lower cognitive reserve and ageing- (>65 years) related cognitive decline^14,15^. A number of studies have used structural and functional MRI modalities or PET to examine the underlying brain basis of CRCI thus showing decreased volume and microstructures in gray and white matter in the prefrontal cortex and parahippocampal gyrus, functional connectivity and brain activities often suggesting damages in frontal cortex, as well as metabolic changes in the posterior cingulate gyrus associated with declined memory in breast cancer patients after chemotherapy^13,16–25^.

The neurobiological mechanisms that underpin CRCI are not completely explained that these neuroimaging studies, but some associations were described between the plasma cytokines IL-1β, IL-6, TNF-α, TNF-RII and cognitive complaints or fatigue or emotion^26–32^. Interestingly, more DNA damage and lower telomerase activity were each associated with higher levels of sTNF-RII and poor attention/decline in motor (movement) speed in breast cancer patient survivors 3 to 6 years after chemotherapy and/or radiation^33^. These data highlight that peripheral inflammation and biological ageing are contributing process in the development of CRCI in cancer patients^34^. However, it remains difficult to bridge the gap between these chemotherapy-resulting systemic maker consequences, the role of cancer itself and cognition, fatigue and/or mood disorders in the onset of CRCI.

More recently, studies have focused the inflammatory and immune-cancer related context in CRCI. Approximately 20-30% of patients with non-central nervous system (CNS) cancer previously reported cognitive deficits after diagnosis and before cancer treatments^13,35–37^. Neuropsychological studies have shown cognitive deficits ranging from 16% to 33% breast cancer patients and verbal episodic memory deficits associated with grey matter atrophy prior to surgery and chemotherapy^37,38^, and/or with increased systemic pro-inflammatory IL-6, IL-8 and TNF-α cytokines^32,39,40^. Cognitive impairment and/or cortical area or thickness^41^ has also been demonstrated in cancer patients with colorectal, lung, testicular of hematological cancers^10,42–49^, with less impact of prostate or ovarian cancer^9,48,50–52^ likely depending on the tumor microenvironment (TME) and/or the stage of its development, while systemic contributors and neurobiological mechanisms are still mostly unknown.

Interestingly, preclinical studies investigated the role of breast cancer on cognitive functions in rodent models and commonly described significant spatial learning and memory impairment associated with reduced volume of frontal lobes and hippocampus, neurogenesis and increased inflammatory markers such as IL-6 in blood of syngeneic models of breast cancer-bearing mice exhibiting^53,54^. Currently, how the local inflammatory status and immune responses within TME can induce alterations in systemic immunity^55^ is currently under debate, which explains why their specific impact on cerebral functions is not known. This is a prerequisite to better explore the distinct impacts of new immunotherapy such as CART-cells and immune check-point inhibitors (ICIs) on cognitive functions^56^.

Solid tumors are now defined as “immunoinflammatory or hot” with a high inflammatory status and immunogenicity (tumoral specific antigens) with TME composed of B-lymphocytes (BL), CD8+/CD4+ TL as well as Treg and myeloid-derived suppressor cells (MDSC) infiltrations, “immune-excluded with high mutation level and CMH presentation, but with a high immunosuppressive stroma and content in MDSC, and “immuno-desert or cold” with absence of inflammation, poor CD8+/CD4+ TL infiltration, and high Treg and MDSC cells^57–59^. With the advent of new immune-based therapies, the constant TME remodeling is emerging as a fundamental concept underlying the impact of cancers and immunotherapy on cognitive and emotional behaviors^60^. Neurological toxicities are frequently observed following immunotherapy, especially CAR-T cells therapies, such as ICANS, but controversial results have been obtained concerning the cognitive impact of immune checkpoint inhibitors (ICI) and CAR-T cell several months following immunotherapy^60^. The severe immune related-adverse events (irAEs) associated with ICI is mainly due to T-cell overactivation, mainly as result of autoimmune reactions of varying organs, including pneumonitis, colitis, hypophysitis and encephalitis^61,62^.

Some patients treated with ICI develop neurological irAEs within 6 to 13 weeks from start of ICI treatment with increased risk in patients with pre-existing auto-immune disease, with melanoma with less advanced oncological disease, or treated with combination of anti-CTLA4 and anti PD-1^63^. Thus, under ICI, a majority of patients with immune related encephalitic diffuse meningo-encephalitic syndrome exhibit altered level of consciousness and/or neuropsychological changes^64^. Most of the irAEs are reversible with immunosuppressive treatments. In the absence of early management of these neurological symptoms by immunosuppressive treatments, long-term cognitive impairment resides, and the mechanisms are rarely investigated^56^. In agreement with the cognitive impact of ICIs, a pre-clinical study previously showed that mice bearing cancer and treated with an anti-CTLA-4 exhibit anxiety behaviors and cognitive deficits; in association with microglial activation within hippocampal brain structures^65^, thus suggesting long-term effects of ICIs on cognitive functions and neuroinflammation. Understanding the mechanistic interplay between systemic immunity, neuroinflammation, and neuronal signaling is therefore critical to anticipate, prevent, and manage cognitive complications in cancer patients undergoing ICI therapy.

The objective of the present study was to investigate activity, emotional reactivity and cognitive functions of immunocompetent mice heterotopically xenografted with murine cancer cell lines (B16F10, B16F10-Ova and MC38) exhibiting various immunoinflammatory status, to relay cancer-related immune-score to animal behaviors and to decipher potential inflammatory and immune mechanisms sustaining the neurobiological defects. Naïve or cancer-bearing mice were also treated by anti-PD-1 and/or anti-PD-L1 while behavioral assessment was performed during the period of tumor growth. Plasma cytokines measurements and FACS-analyzed immune cell proportion and diversity were performed to establish key blood signatures of the effects of ICI combined with the different immunogenic cancers on behaviors and vascular permeability, brain neurotoxicities and neuroinflammation. To highlight a PD-L1-evoked specific γδ-TL subpopulation with an innate and adaptive functions on neurocognitive dysfunctions and brain inflammation, cancer bearing mice treated by anti-PD-L1 were challenged with an antibody directed against γδ-TL. We here unraveled for the first time the impact of immunogenic cancers on emotional and cognitive functions as well as BBB integrity and neuroinflammation and demonstrate the priming effect of cancer of anti-PD1 and more importantly anti-PD-L1 on γδ-TL-mediating anxiety and brain dysfunctions. These data should contribute to the development of strategies of diagnosis, prevention of mitigation of ICI-induced CRCI, thereby improving QoL of cancer patients and survivors.

## Material and methods

### Animals

C57BL/6J male mice aged of 8 weeks were purchased at Janvier Laboratories (Janvier, Le Genest Saint Isle, France) and were housed (5 animals per cage) under controlled conditions (22 ±1°C; 12h/12h light/dark cycle; ad libitum access to food and water). Upon receipt, mice were randomly assigned to treatment groups. Group and sample size for each experiment are indicated in each figure legend. The number and the suffering of animals were minimized in accordance with the guidelines of the European Parliament and Council Directive (2010/63/EU): This project was approved by the “Comité d’Ethique NOrmandie en Matière d’EXperimentation Animale” and the French Research Minister (APAFIS#25535-2020032512491133 v2).

### Cell culture

B16F10, B16F10-Ova and MC38 cell lines were kindly provided by Pr Sahil Adriouch of U1234 PANTHER Laboratory, Rouen, France. MC38 cells were cultured in Dulbecco’s Modified Eagle Medium (Gibco, Life Technologies, USA) containing 4.5 g/l D-Glucose and L-glutamine 580 mg/l, added with 110 mg/l sodium pyruvate (Thermo Fisher Scientific, Illkirch, France). The medium was supplemented with 10% fetal bovine serum (Gibco, Life Technologies, USA), 1% Penicillin-Streptomycin Solution (100 IU/ml penicillin, 100 µg/ml streptomycin) (A5955, Sigma-Aldrich, Darmstadt, Germany), 1× MEM Nonessential Amino Acids (Thermo Fisher Scientific) and 1 mM HEPES (Gibco, Life Technologies). B16F10 and B16F10-Ova melanoma cells were cultured in RPMI 1640 medium (Gibco, Life Technologies) supplemented with 10% fetal bovine serum (Gibco, Life Technologies), 100 μg/ml of streptomycin and 100 units/ml of penicillin. Cells were maintained in a humidified incubation chamber at 37°C and 5% CO2. Cells were passaged every 2–3 days or at 50%–70% confluency by lifting with 0.05% Trypsin-EDTA and subculturing at 1:10–1:15 ratios.

### Cancer model and treatment

Low passage cells were resuspended at 5x10^4^ cells/100 μl in PBS for MC38 and at 5x10^4^ cells/150 μl of matrigel (356231, Sigma-Aldrich) for B16F10 and B16F10-Ova cells and injected subcutaneously on the shaved right flank of C57BL/6 mice. Tumor volume growth was monitored twice per week and calculated using the formula (mm3) = 0.5 x (length) × (width)^2^. In our study, endpoints were defined as tumor volume superior to (1,5 cm^3^) and occurrence of any signs of suffering. To prepare tumor cell lysates, B16F10, B16F10-Ova and MC38 cells were divided among 1.5 ml tubes (10^6^ per 50 µl phosphate-buffered saline (PBS) per tube), and the tubes were subjected to 3 freeze-thaw cycles and stored at −80°C until use. The viability of cells was assessed using trypan blue staining (15250061, ThermoFisher Scientific). B16F10, B16F10-Ova or MC38 lysates were inoculated intradermally into the right flank of mice. Each immunization was performed twice a week for three consecutive weeks. 5 days (B16F10, B16F10-Ova) or 10 days (MC38) after tumor inoculation, mice were injected intraperitoneally with 5 mg/kg of anti-PD-1 (clone10F.9G2, BioXCell), anti-PD-L1 (clone BH7-1, BioXCell) or the respective isotype control IgG2a (Rat, clone 2A3) twice a week. Naïve C57Bl/6J mice were injected intraperitoneally with 100 mg/kg of anti-PD-1 (clone 10F.9G2, BioXCell), anti-PD-L1 (clone BH7-1, BioXCell) or the respective isotype control IgG2a (Rat, clone 2A3) for 3 weeks, twice a week.

### Behavioral Tests

Depending on tumor growth (B16F10/B16F10-Ova: D4-8, MC38: D13-16, lysates: D16-19, cancer-naïve mice: D22), short-term memory (NORT), spontaneous activity (OFT), anxiety (EPM and LDB) and resignation (TST and FST) were studied using a battery of behavioral tests. For all behavioral analyses, observers were blinded to group (tumor vs. treatment). At the end of experimental sessions, brain, tumors and blood samples were collected from each group to lead immunohistochemical experiments and quantification of cytokines and peripheral immune cells, respectively.

#### Open field Test-OFT

The spontaneous activity was measured for 10 min by placing mice in an OFT apparatus (45 × 45 × 31 cm, L x l x H). The animal was placed into the center of the apparatus and several behavioral parameters (distance covered, speed, vertical activity and immobility) were automatically analyzed by the video tracking system Anymaze (Stoelting^®^, Dublin, Ireland).

#### Novel Object Recognition Test-NORT

Mice were familiarized to the OF arena (30 × 30 × 31 cm, L x l x H) for 10 min as described above. The following day, mice were exposed to the arena (5 min) containing two identical objects. Two hours after, one of the identical objects (“familiar”) was replaced with a novel object of similar dimensions, but different colors and textures, and mice were again allowed to explore the arena for 5 minutes. Clear visuospatial orientation to the object, as well as physical interaction with the object was coded as exploratory behavior, and the percent time spent exploring the novel versus the familiar object was calculated. The preference index was calculated as the ratio of the time spent exploring the new object to the total exploration time: Preference Index = (T_new_)/(T_new_ + T_familiar_). The discrimination index was calculated as the ratio of the difference between the time spent exploring the new object and the familiar object to the total exploration time Discrimination Index = (T_new_ − T_familiar_)/(T_new_ + T_familiar_).

#### Light and dark box-LDB

The light/dark exploration test was conducted to assess anxiety-like behavior. The light-dark box consisted of a dark compartment 40 × 19 × 35 cm (L x l x H) with a 7 × 7 cm aperture at floor level that opened onto a large plexiglas square arena (light compartment: 40 × 19 × 35 cm, L x l x H). In this test, mice were placed in a dark compartment of the light-dark box. The light intensity in the light chamber was kept around 120 lux. The number of entries into the light compartment (defined as all 4 paws out of the shelter) and time spent inside the light compartment over a 10 min session was calculated. Mice that did not enter the light compartment were assigned a 600 sec latency to enter.

#### Elevated Plus Maze-EPM

The anxiety-like behaviors of mice were assessed by means of the EPM. The device consists of four arms: two opposite secure arms closed by 19.5 cm vertical walls and two aversive open arms perpendicular to the previous ones with a 0.5 cm high rim (each arm is 25 cm long and 5 cm wide) and a central area of 5 × 5 cm, elevated at 41.5 cm above the ground. At the beginning of the test, the mouse was placed in the central area of the maze with its head pointing toward an open arm. The time spent in the open and closed arms, the number of entries as well as the number of head dips and stretch-attended postures (SAP) were measured. The animal’s behavior was videotaped and analyzed using the Anymaze software (Stoelting, Dublin, Ireland) for a single 6 min session.

#### Tail Suspension Test-TST and Forced swim test-FST

Depressive-like behaviors were evaluated in the TST. Mice were suspended at a height of 20 cm above the floor, by the use of adhesive tape applied to the tail, in a three-walled rectangular box (16 x 15 x 31 cm, L x l x H). The total duration of immobility (passive hanging) between periods of wriggling to avoid aversive situation as well as the latency to the first immobility were measured for a period of 6 min as an indication of the behavioral despair of mice. In FST, mice were placed for 6 min into a cylinder (17 cm in diameter) filled with 25°C tap water (at a height of 13 cm) with no way out. The immobility duration (excluding movement necessary to keep the head above water or to float) was measured as an indication of the behavioral despair of mice.

### Immunofluorescence

Mice were anesthetized using a ketamine/xylazine cocktail (100 mg/kg and 12,5 mg/kg) and killed by transcardiac perfusion with 15 ml ice cold PBS 1X (MFCD00131855, Sigma Aldrich, Germany) followed by 25 ml of 4% paraformaldehyde (PFA, 158127, Sigma Aldrich, Germany). Brains were post-fixed in 4% PFA overnight at 4°C and cryoprotected in 30% sucrose (15503022, Sigma Aldrich, Germany) for 24 hours at 4°C and then stored in PBS 1X until used for immunohistochemistry. Immunofluorescence was performed as follows. Free-floating sections were cut at 50 μm by using a vibratome (Leica Biosystems, VT 1000S, Germany). Sections were permeabilized with 0.5% Triton X-100 (10717503, Invitrogen, Illkirch, France) and then incubated for 1 hour at room temperature in blocking reagent composed by 5% normal donkey serum (D9663-10ML, Sigma Aldrich, Germany) and BSA 10% (A7030, Sigma Aldrich, Germany) in PBS 1X. Sections were then incubated in primary antibody (listed below) in blocking reagent overnight at 4°C, followed by incubation in secondary antibody (also listed below) for 2 hours at room temperature. Between each stage, sections were washed thoroughly with PBS 1X. Sections were incubated (5 min) with DAPI solution (28718-90-3, Sigma Aldrich, Germany) for nuclear staining and mounted with Mowiol solution (Merck Millipore, 475904) for microscope observations. Images were obtained by SP8 confocal microscopy (PRIMACEN, Upright confocal microscope, Leica Microsystems, Nanterre, France) and THUNDER imaging system (PRIMACEN, Upright confocal microscope, Leica Microsystems, Nanterre, France).

**Table 1.**
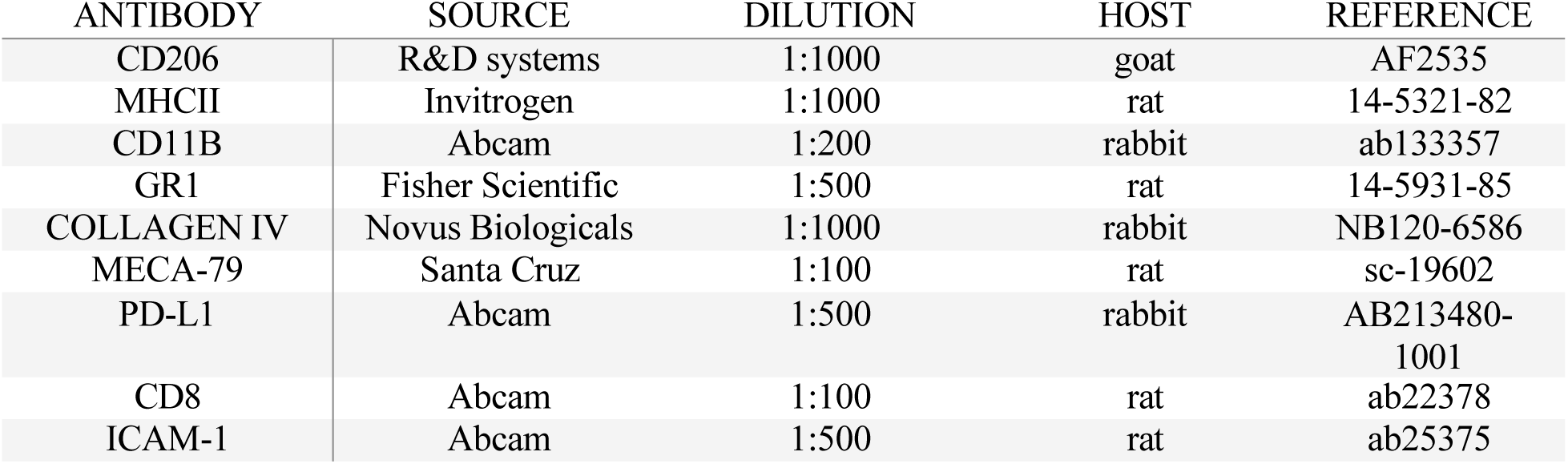
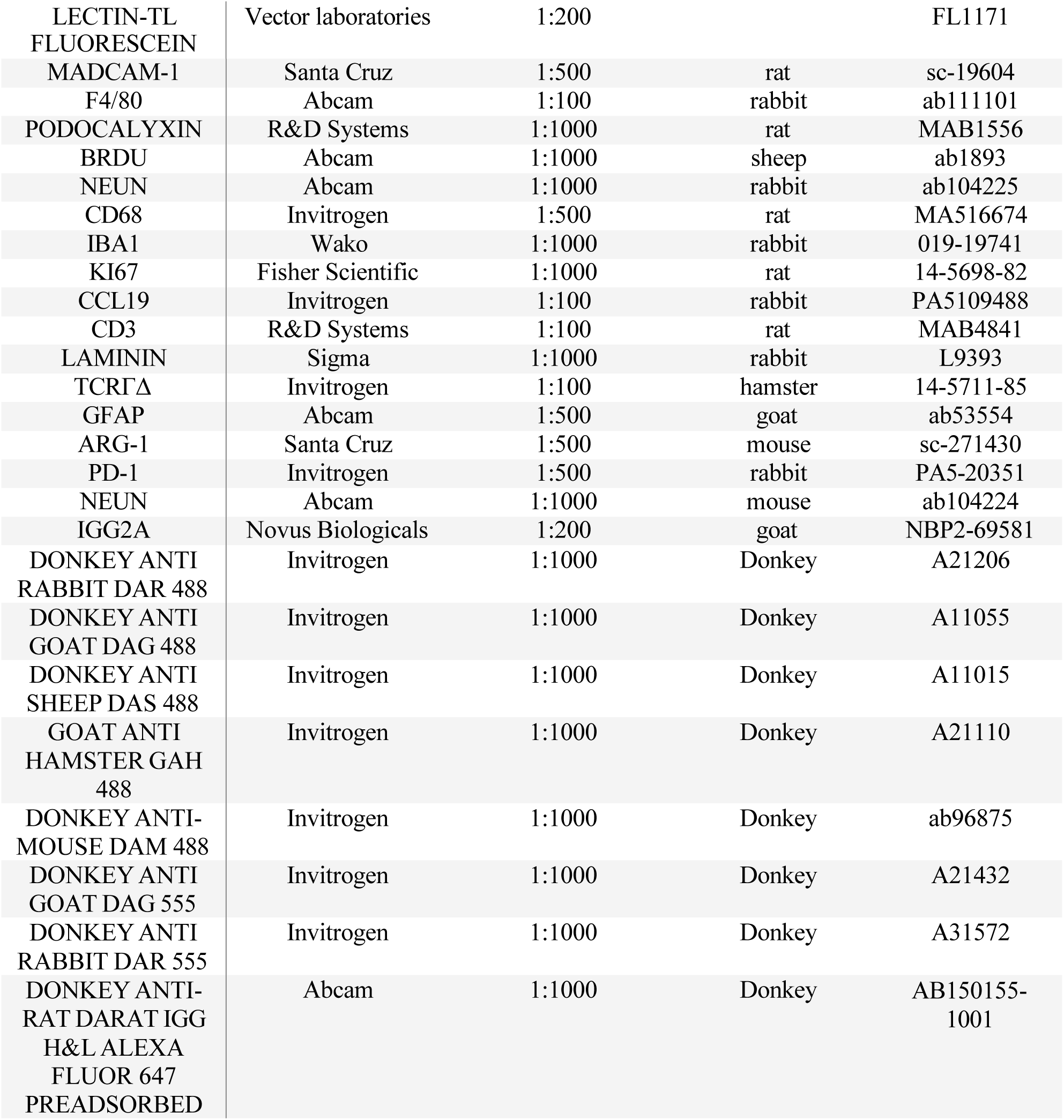
List of the antibodies used.

### Morphological analysis of Microglia

Image processing was performed following the protocol described by Young et al^66^. Color images of tissue sections labelled with the Iba1 antibody (Rabbit, Wako, 019-19741, dilution 1:1000, USA), were obtained using a fluorescence microscope (THUNDER, Leica) with Z-stack acquisition capability (30 μm) using a 63X objective. The brain area analyzed was the hippocampus (1.08 mm from the Bregma -1.46 mm). Cells were selected and cropped according to the following criteria: (i) random selection at the level of the dentate gyrus of the hippocampus (ii) no overlap with neighboring cells and (iii) complete nucleus and branches (at least in appearance). Selection was blinded. The analysis considered 4 animals per group, 4 images per animal and two cells per image, for a total of 8 cells per hippocampus selected from each animal. The density, fluorescence intensity and total number of cells were quantified using Image J software (version 1.8.0, NIH, USA). The fractal dimension^67^, lacunarity^68^, the size of the soma and length and number of branches per cell were quantified^69^.

## Meningeal whole mount preparation and immunostaining

Dissection, isolation and immunostaining of whole meninges were performed as described by Louveau et al.^70^. Briefly, after anesthesia by isoflurane and decapitation, the animal skullcap was harvested in ice-cold PBS containing 5 U/ml heparin and then placed in 4% PFA (158127, Sigma Aldrich) for 15 min at 4°C. After rinsing in PBS (MFCD00131855, Sigma Aldrich), whole meninges were dissected under a microscope by gently scraping the meninges from the skull using fine forceps. For immunostaining, whole mount meninges were incubated on a shaker for 1h at RT in 300 μl of PBS with 2% normal donkey serum (D9663-10ML, Sigma Aldrich) and BSA 1% (A7030, Sigma Aldrich, Germany) then overnight at 4°C in primary antibodies diluted in PBS with 1% BSA and 0.5% Triton-X-100. After three rinses in PBS, the meninges were incubated for 2 hours at RT in 300 μl of secondary antibodies diluted in PBS with 1% BSA and 0.05% Triton-X-100. Finally, meninges were flattened on a glass slide, dried and mounted in Mowiol (Merck Millipore, 475904) before examination with THUNDER imaging system (PRIMACEN, Upright confocal microscope, Leica Microsystems, Nanterre, France).

## Quantitative Real-Time PCR

Prior to tissue extraction, mice were anesthetized using a ketamine/xylazine cocktail (100 mg/kg and 12,5 mg/kg) and killed by transcardiac perfusion with 15 ml ice cold PBS1X (MFCD00131855, Sigma Aldrich). Hippocampus, prefrontal cortex, somatomotor cortex, cerebellum and dura mater were dissected, snap frozen, and stored in −80°C until analysis. Dura mater was dissected as described above. Total RNAs were extracted with Trizol reagent (Invitrogen, Waltham, Massachusetts, USA) and further purified with the NucleoSpin® RNA Plus (Macherey-Nagel, ThermoFisher scientific, Waltham, Massachusetts, USA). The RNA concentration was determined by measuring absorbance on a Nanodrop (ThermoFisher scientific) at 260 nm and nucleic acid purity was verified with 260/280 nm and 260/230 nm ratio. cDNA was synthesized from 1 µg of RNA using the Im-Prom II Reverse Transcriptase (Promega, Madison, Wisconsin, USA). RNA was extracted using a RNeasy mini kit (Qiagen, Hilden, Germany) according to the manufacturer’s instructions. cDNA was transcribed using TaqMan reverse transcription reagents and random hexamers according to the manufacturer’s instructions. The determination of the expression level of genes was done by real time PCR in 384-well plates with a 5 µl reaction volume in the presence of 1X Fast SYBR Green PCR Mastermix (4385612, Thermofisher,) containing pre-set concentrations of dNTPs, MgCl2 and the SYBR Green reporter dye along with a set of specific primers. The distribution of cDNA samples and reaction mixes was performed by a Bravo Automated Liquid Handling system (Agilent, CA, USA). The PCR reaction was conducted with a QuantStudio 12 k Flex thermal cycler (ThermoFisher scientific). Relative expression was calculated using the ΔΔCt method and ACTB was used as reference gene. Statistical analysis was performed on the normally distributed ΔCt values. The IPA system (version 42012434, Ingenuity Systems, Qiagen China Co., Ltd.) was used for subsequent bioinformatics analysis, which included canonical pathway analysis, disease and function, regulator effects, upstream regulators and molecular networks. IPA uses a network generation algorithm to segment the network map between molecules into multiple networks and assign scores for each network^71,72^. The score is generated based on hypergeometric distribution, where the negative logarithm of the significance level is obtained by Fisher’s exact test at the right tail. For canonical pathway analysis, disease and function, the −log (P-value) >2 was taken as threshold, the Z-score >2 was defined as the threshold of significant activation, whilst Z-score <−2 was defined as the threshold of significant inhibition. For regulator effects and molecular networks, consistency scores were calculated, where a high consistency score indicates accurate results for the regulatory effects analysis. For upstream regulators, the P-value of overlap < 0.05 was set as the threshold.

**Table 2.**
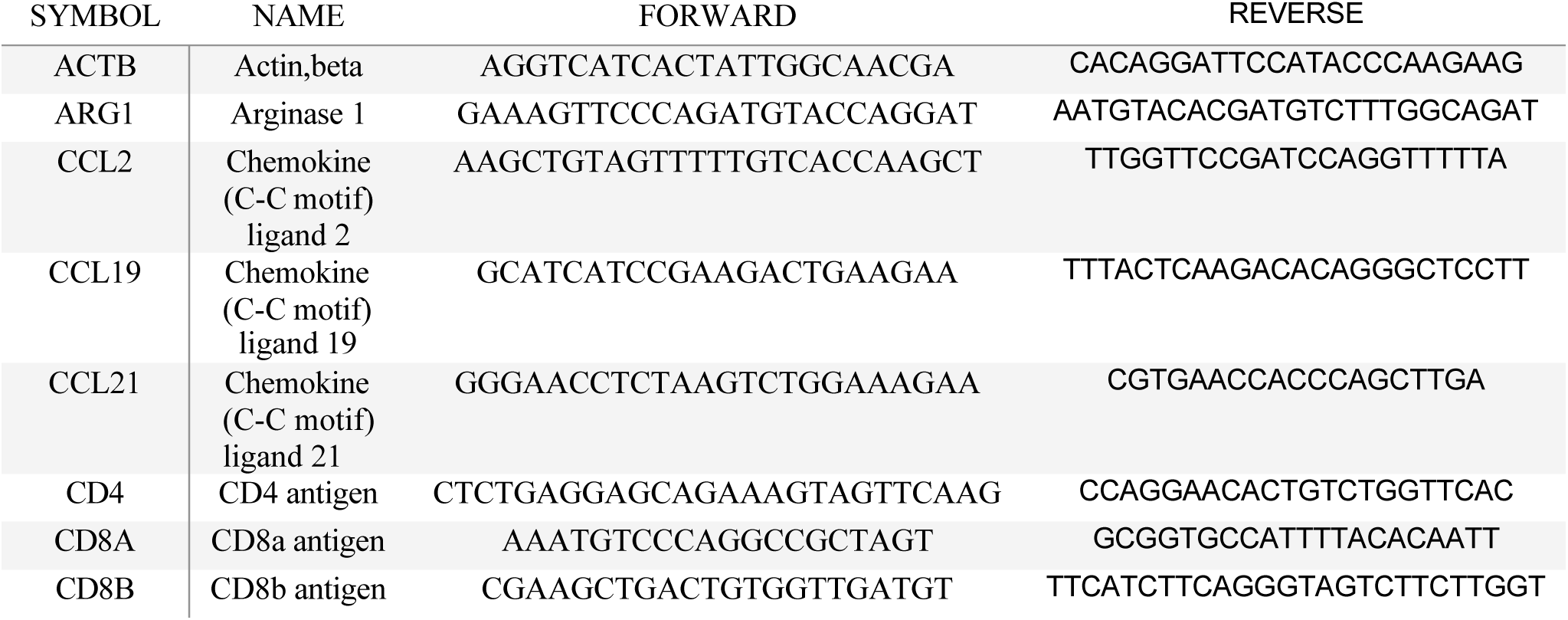
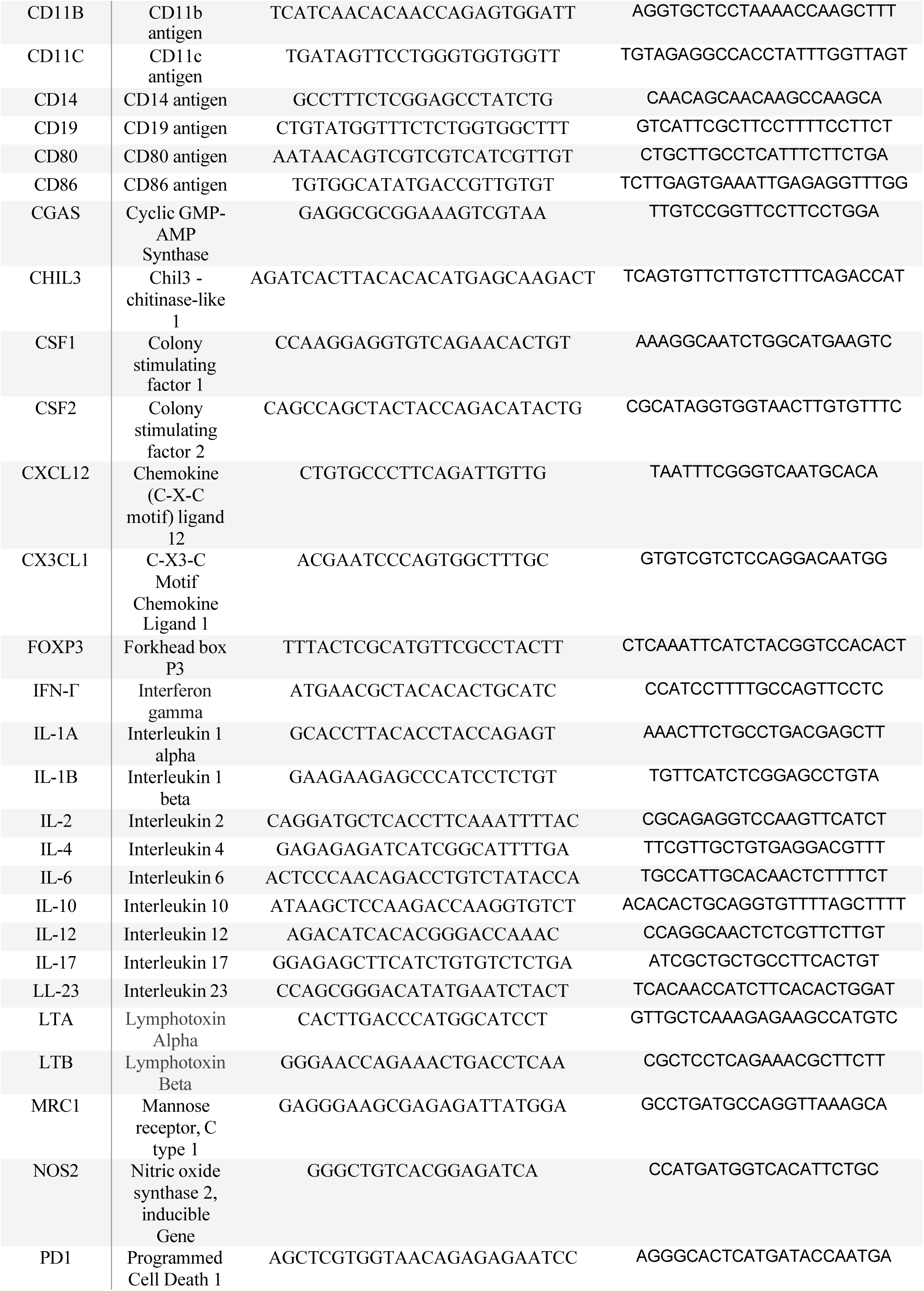
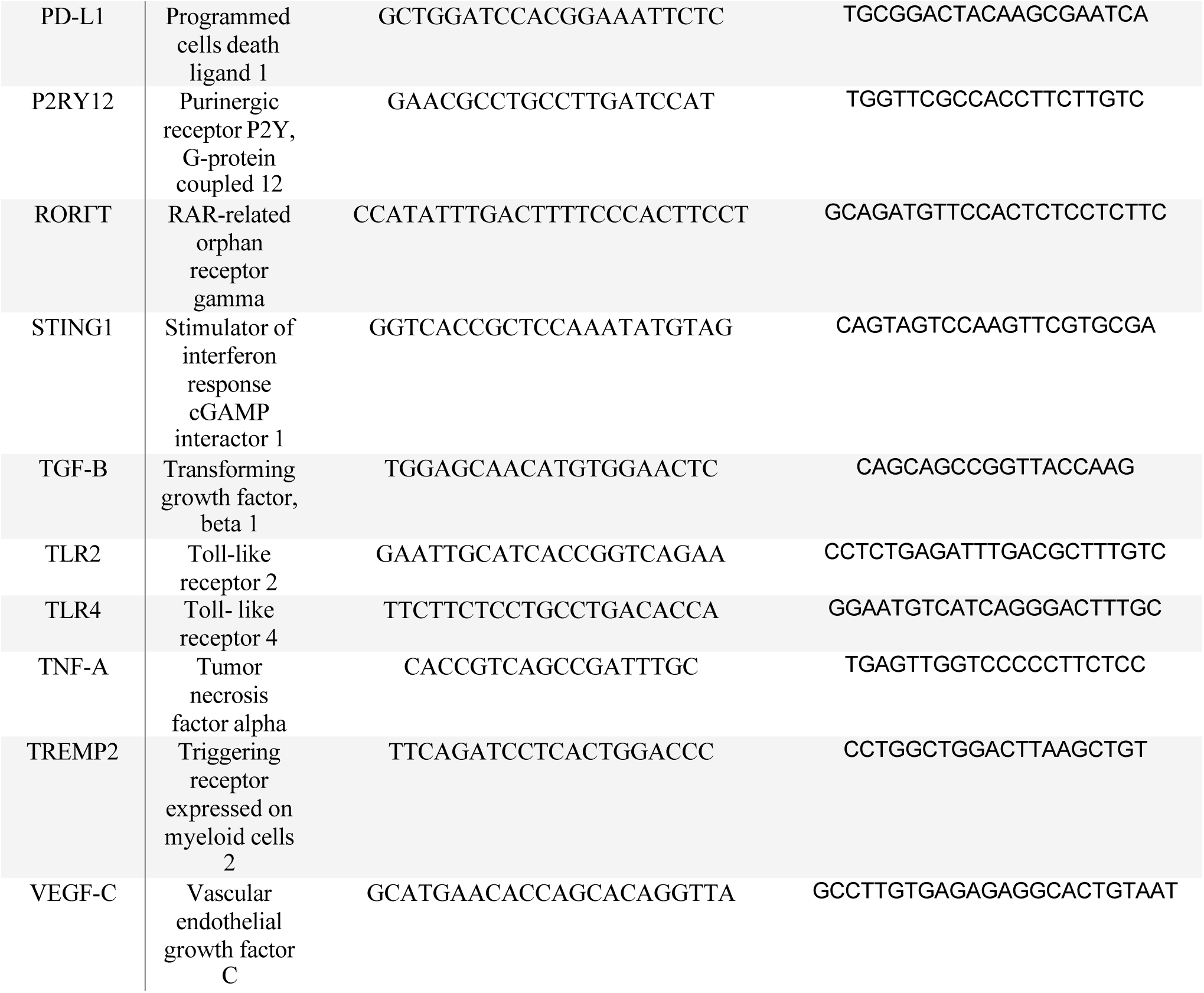
List of couples of primers used.

## Flow cytometry

Prior to perfusion, mice were anesthetized using a ketamine/xylazine cocktail (100 mg/kg and 12,5 mg/kg) and blood was collected by cardiac puncture using a 25-gauge needle, then placed in an EDTA coated tube. Red blood cells were then lysed with 1X RBC lysis buffer (00-4333-57, Invitrogen). The resulting cell suspension was washed with FACS Buffer (BSA 1%, EDTA 0.5 mM) then cells were incubated in 100 μl of FACS buffer containing antibodies solution for 30 min at 4°C. Cell suspension was then washed with FACS buffer and stored in 500 µl of FACS buffer at 4°C until use.

## Gating Strategy

Cells were gated on SSC singlet, and FSC singlet. Immune cells were defined as CD45^+^ cells. From myeloid cells, dendric cells (CD11b^+^CD11c^+^), monocytes (CD11b^+^CD11c^-^), macrophages (CD11b^+^Ly6C^-^Ly6G^-^), anti-tumoral macrophages (CD11b^+^Ly6C^-^Ly6G^-^MHCII^+^), pro-tumoral macrophages (CD11b+Ly6C^-^Ly6G^-^CD206^+^), monocytic derived-myeloid suppressor cells (CD11B^+^Ly6C^+^Ly6G^-^), neutrophils (Cd11b^+^Ly6C^-^Ly6G^+^MHCII^-^) and polymorphonuclear-myeloid suppressor cells (Cd11b^+^Ly6C^low^Ly6G^+^MHCII^+^) were identified. From lymphocytes, CD3^+^ cells were identified as T lymphocytes, and further phenotype as CD4^+^ T helper or CD8^+^ cytotoxic T lymphocytes, Th17 lymphocytes (CD4^+^RORγT), regulatory (CD4^+^FoxP3^+^) or TCRγδ (CD3^+^TCRγδ^+^) T lymphocytes. CD3^-^ cells were divided into NK1.1^+^ natural killer cells or CD19^+^ B lymphocytes. Flow cytometry analysis was performed on a BD Sony analytic flow cytometer (Sony biotechnology, San Jose, California).

## Antibodies and FACS analyses

All antibodies were used at 1:1000 dilution. The following anti-mouse antibodies were used, with clone, fluorophore, and reference indicated in parenthesis: CD45 (REA737, VioGreen, 130-110-803, Miltenyi Biotec), CD11b (M1/70, FITC, 101206, Biolegend), CD11c (N418, PerCP, 117328, Biolegend), MHCII (M5/114.15.2, BV605, 107639, Biolegend), CD206 (APC, FAB2535A, R&D systems), Ly6G (1A8, BV421, 1238135, Sony Biotechnology), Ly6C (REA796, Vioblue, 130-111-921, Miltenyi Biotec), CD3 (17A2, BV421, 100228, Biolegend), CD4 (REA1211, PE, 130-123-206, Miltenyi Biotec), CD8 (53-6.7, FITC, 11-0081-82, Invitrogen), NK1.1 (PK136, PerCP, 1143630, Sony Biotechnology), CD19 (6D5, AF700, 115527, Biolegend), FoxP3 (MF-14, AF647, 1232040, Sony Biotechnology), RORγt (Q31-378, BV786, BD Biosciences) and TCRγδ (GL3, BV605, 118129, Biolegend).

## Cytokines Quantification

Blood (400 µl) was sampled by cardiac puncture and stored into tubes coated with EDTA to prevent coagulation. Samples were kept on ice during the whole procedure. Blood tubes were centrifuged for 10 min at 4°C at 1300 G. Supernatant plasma was isolated and stored frozen (-80°C) until use. The plasma concentration of cytokines was analyzed by the enzyme-linked immunosorbent assay (ELISA) according to the manufacturer’s recommendations (Q-Plex Mouse Cytokine-Inflammation, Quansys Biosciences, Logan, UT, USA). The cytokines measured were: IL-1β, IL-1α, IL-6, IL-13, IL-10, IL-17, Granulocyte Macrophage Colony-Stimulating Factor (GM-CSF), Monocyte Chemotactic Protein-1 (MCP-1). Individual ELISA Kit was used to measure IL-4 (Invitrogen, BMS613), TNF-α (Invitrogen, BMS607-3) and ICAM-1 (Invitrogen, EMICAM1ALPHA) plasmatic concentration. The chemiluminescence from the 96-well plates was imaged with ChemiDoc XRS+ (Bio-Rad, Marnes–La–Coquette, France) and an 8-point calibration curve (7 points plus 1 blank). Then, the images of the Q-plex plates were analyzed in Q-ViewTM software (Quansys Biosciences) to obtain the standard curves and sample values in pg/ml. All data are expressed in pg/ml. Undetectable values are expressed as half of the minimal quantity detected by the kit within this experimental sequence.

### *In vivo* BBB permeability assay

*In Vivo* Blood-brain Barrier Permeability Assay was performed as described previously^73^. Briefly, PBS-mice and B16F10- and MC38-bearing mice, receiving or not anti-PD-1, anti-PD-L1 or IgG2a were intraperitoneally injected with FITC-labelled 3 kDa dextran (100 µl of 2 mM stock in PBS, Sigma Aldrich, 46944-100 MG-F, Darmstadt, Germany), followed by anesthesia (ketamine/xylazine cocktail) 5 min later. After a circulation time of 10 min for the tracer, the animals were prepared for transcardiac perfusion with PBS 1X, 400 µl of blood were collected by cardiac puncture, and brains were cut into two hemibrains immediately frozen at −30°C in Isopentane (PHR1661, Sigma Aldrich) and stored at −80°C. For fluorescence measurement, first series of hemibrains were thawed on ice, weighed and homogenized in PBS, followed by centrifugation (15000 G) for 20 min at 4°C. Supernatants (100 µl) as well as equal volumes of serum (1/5 dilution in PBS 1X) were loaded into a 96-well black plate and fluorescence was measured at the corresponding excitation (490 nm)/emission (520 nm) in a plate reader (Tecan, Männedorf, Switzerland). Sham animals (without tracer injection) were used to subtract autofluorescence values. The permeability index (ml/g) was calculated as the ratio of tissue RFUs/g tissue weight to serum RFUs/ml serum. For fluorescence tracer extravasation imaging, the remaining hemibrains were cut into 30 µm slices with cryomicrotome (CM 3050S, Leica, Heidelberg, Germany, PRIMACEN platform) and stained as previously described.

### *In vivo* depletion assays

To deplete peripheral macrophages B16F10- or PBS-mice were intraperitoneally injected with clodronate liposomes (CLD-8909, Encapsula Nanoscience, Brentwood, USA) (110 mg/kg, 100 µl) at day1, day 4 and day 8 while MC38-bearing mice were injected with clodronate at day 7, day 10 and day 13 after tumor inoculation (Day 0). The control animals were injected with an equal volume of empty liposomes. 3 days after the last intraperitoneal injection of clodronate liposomes or empty liposomes, the mice were transcardially perfused, 400 µl of blood were collected by cardiac puncture, and brain were collected. To deplete TCRγδ Lymphocytes, anti-TCR γδ (UC7-13D5, BioXCell) or isotype control (polyclonal Armenian hamster IgG, BioXCell), were injected intraperitoneally (500 μg in 250 μl of NaCl) on D4 and D7 after tumor inoculation for B16F10-bearing mice and on D10 and D13 after tumor inoculation for MC38-bearing mice. These mice received concomitant administration of anti-PD-L1 (5mg/kg, clone BH7-1, BioXCell) or isotype control α-IgG2a (Rat, clone 2A3), injected as detailed above.

## Statistical Analysis

Prior to analysis, each experimental condition group normality was checked by Shapiro-Wilk normality test and variances were analyzed. Outliers were identified by using the Grubs test (alpha: 0.05). Multiple comparisons were performed by using one-way ANOVA or the Kruskal– Wallis test (for non-normally distributed data) followed by Bonferroni or Dunn’s correction, respectively. T-tests or Mann‒Whitney tests (for non-normally distributed data) were used for comparisons of two groups. To test the different evolution between two groups over time, a two-way ANOVA followed by Sidak’s multiple comparison test. Correlations were calculated by linear regression (Spearman’s r_s_). Differences were considered at significant at p < 0.05. Quantification was performed using ImageJ software. Statistical analysis was performed using GraphPad Prism-7 software (la Jolla, CA, USA). Data are presented as the mean ± SEM.

## Results

### Cognitive deficits and emotional reactivity in mice bearing immuno-desert, immuno-excluded and immuno-inflamed cancers

To evaluate the impact of cancer-associated immune status on cognitive functions and emotional reactivity, we studied the melanoma cell line B16F10 (B16F10), known to be poorly immunogenic^74,75^, the more immunogenic melanoma B16F10-Ova (B16F10-Ova)^76,77^ and the highly immunogenic colon cancer MC38 cell line^78^, which were subcutaneously injected in the flank of C57B/l6 immunocompetent mice **(Fig. 1a)**. A battery of behavioral tests was performed on B16F10, B16F10-Ova and MC38-bearing mice to investigate spontaneous activity in the open field test (OFT), anxiety in the elevated plus maze test (EPM), resignation in the tail suspension test (TST) and short-term memory/executive functions in the novel object recognition test (NORT) (**Fig. 1a**). To decipher the contributing impact of immune challenge and of the immunogenic TME animal behavioral, tumoral lysates obtained by freezing and thaw cycles derived from B16, B16-Ova and MC38 cell lines were intradermally injected once/week for two weeks before behavioral assessment.

**Figure 1.**
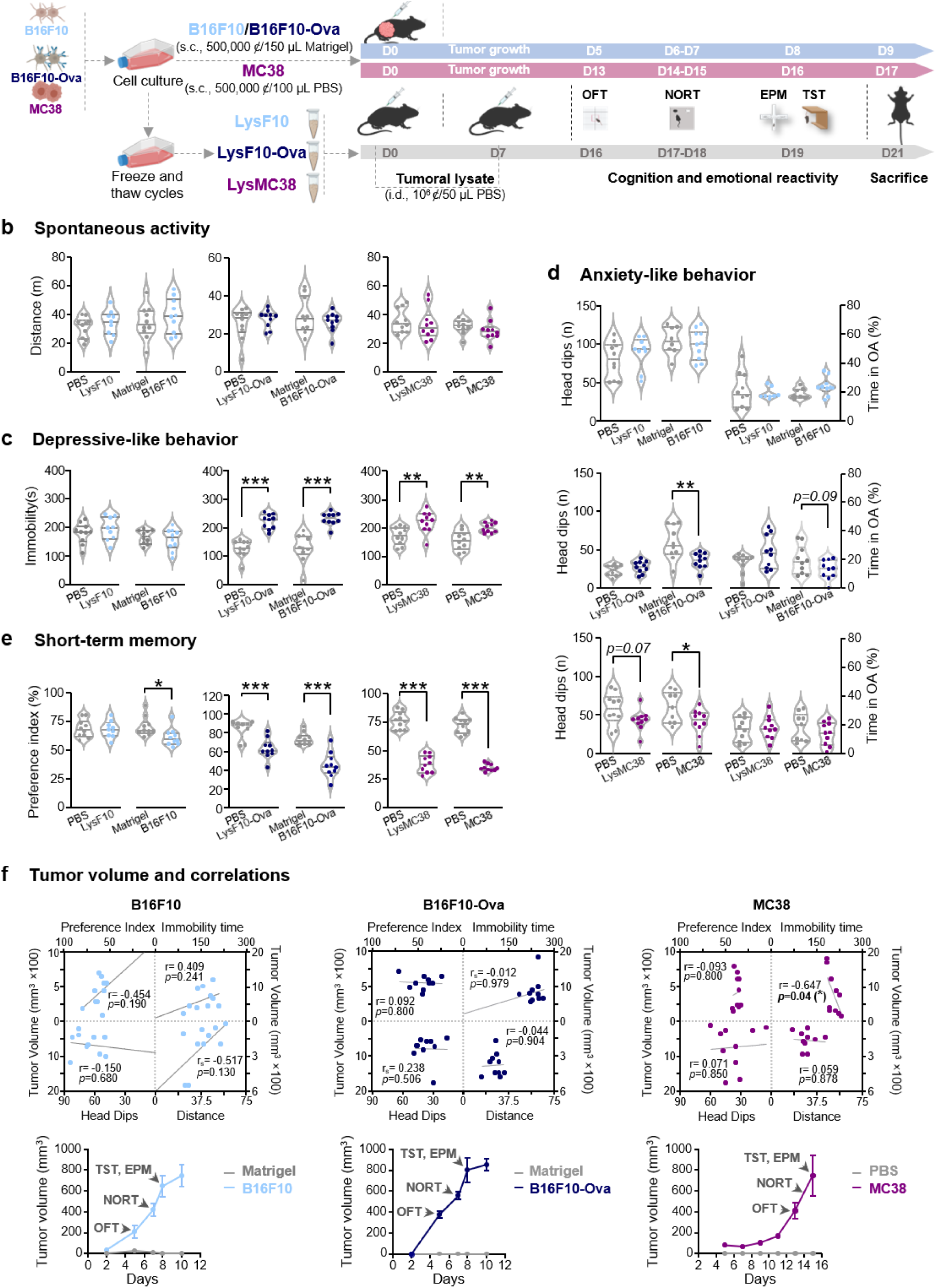
Impact of cancer-associated immune challenge on activity, emotional reactivity and cognition in mice. **a.** Experimental timeline of tumoral cell lines B16F10-(blue), B16F10-Ova (blue) and MC38 (purple) or their respective tumoral lysates inoculation and behavioral assessment in mice. B16F10/B16F10-Ova tumoral cells (500000ȼ/150μl matrigel) or MC38 (500000ȼ/100 μl PBS) were subcutaneously (s.c.) injected in the right flank of C57Bl/6 mice (n=10). Tumoral lysates were obtained by freeze and thaw cycles (3x) and they were injected intradermally (i.d.,10^6^ȼ/50 μl PBS) at D0 and D7 on the right flank of mice. Depending on the kinetic of tumor growth (B16F10/B16F10-Ova: D5-8, MC38: D13-16, lysates: D16-19), spontaneous activity in the open-field test (OFT), cognitive functions in the novel object recognition test (NORT), anxiety in the elevated plus maze (EPM) and resignation in the tail suspension test (TST) were evaluated. Mice were sacrificed the day after the end of the behavioral session and blood was collected. D: day, i.d.: intradermally. **b-e.** Impact of B16F10, B16F10-Ova, MC38 tumors or of their respective lysates on spontaneous activity (left panel), anxiety (middle panel) and resignation (right panel) (**b),** depressive-like behavior (**c),** anxiety-like behavior (**d**) and short-term memory (**e**) (n=10). Statistical analyses using Unpaired t-test or Mann-Whitney test; Data are represented as violin plots with symbols for individual data points and median in dotted line. Data are expressed by mean ± SEM, * p<0.05, ** p <0.01, *** p <0.001. **f. *Top*,** Regression curves showing correlations between preference index (cognition), immobility (resignation), number of head dips (anxiety), distance (activity) and the tumor volume in in B16F10-, B16F10-Ova- and MC38-bearing mice (n=10). Statistical analyses were performed by using Spearman correlation. ***Bottom*,** Tumor growth kinetic curves as a measure of volume of B16F10- (blue), B16F-Ova- (dark blue) and MC38-(purple)-bearing mice compared with respective controls matrigel (light grey) or PBS (dark grey) mice (n=10). The timepoints of behavioral tests were indicated by black arrows.

Indeed, whole-tumor lysate constitutes a valid alternative source of cancer antigens and is expected to be effective to stimulate CD4^+^ helper T and CD8^+^ TLs, in the absence of TME **(Fig. 1a).** The locomotor and spontaneous activity were impaired only in MC38-bearing mice, no significant differences being observed between B16 or B16-Ova-bearing mice or lysates-injected mice compared with respective noncancer mouse controls **(Fig. 1b).** Resignation, considered as being a marker of depressive-like behavior, was also impaired in B16-Ova- and MC38-bearing mice and corresponding lysates-injected mice compared with their respective controls **(Fig. 1c).** The anxiety behavior revealed by a diminished number of head dips in the EPM was increased in B16-Ova-mice and in both MC38-lysate-injected and MC38-bearing mice **(Fig. 1d).** Then, cognitive performances were evaluated in the NORT consisting in evaluation of the time spent on the novel object compared to the time spent on the familiar object (preference index) and the duration of exploration for the new object compared to the familiar one as a proportion of the animal’s total exploration time. Here a significant decrease of preference and discrimination index was measured in all cancer bearing mice and in B16-Ova- and MC38-lysates-injected mice compared with control groups **(Fig. 1e).** These data indicate that independent of the immuno-status, cancers’ development results in impairment in attention processes and/or short-term memory, and that more the cancer is immune-inflamed, more spontaneous activity and emotional reactivity were found gradually altered. The absence of correlation between behavioral results and tumor mass volume allows us to exclude influence of tumoral mass volume on behavioral results obtained for B16F10 and B16F10-Ova, except for the immobility time of mice bearing MC38, showing an increased depressive-like behavior with the size of the tumor (**Fig. 1f).** Globally, cognitive functions and emotional reactivity were altered relative to the cancer immune challenge, likely associated with TME.

The immuno-inflamed or immune-desert TME of B16F10, B16F10-Ova and MC38 was characterized through immunohistochemical investigation of immune challenge markers. In 2D tumor sections, the number of MHC-II^+^ anti-tumoral tumor-associated macrophages (TAM) and CD11b^+^ monocytes were quantified in tumoral bulk showing higher levels of MHC-II+ and CD11b^+^ cells in B16-Ova and MC38 compared with B16 tumors (**Fig. 2a)**. In addition, a lower density of pro-tumoral CD206^+^ TAM and CD11b^+-^GR1^+^ MDSCs was detected in MC38 compared with B16 tumors (**Fig. 2a)**, thus suggesting that MC38 is indeed immune-inflamed, infiltration of CD4^+^ and CD8^+^ LTs as well as an increased presence of Treg and MDSC^58^ whereas inflammatory tumors are defined by important leukocyte infiltration, including CD4^+^/CD8^+^ LTs, Treg, TAMs or MDSCs^79^. The density of LTs but also LBs frequently correlates with the presence of tumor-associated high endothelial venules (HEVs)^80–82^, blood vessels specialized in recruiting lymphocytes to lymph nodes and other lymphoid organs^83^. HEVs express high levels of sulphated sialomucins (Meca-79) recognized by the lymphocyte receptor L-selectin (CD62L)^83^. In Figure 2b, thus, tumoral vascular network and HEV coverage were investigated by means of antibodies directed against collagen IV (green) and Meca76 (magenta). In MC38 tumors compared with B16F10 and B16F10-Ova, larger Meca-79^+^collagenIV^+^ HEV in tumor burden (magenta, left panel) and higher density of CD8^+^ TL (cyan right panel) (**Fig. 2b**). PD-L1 can be expressed by many cell types, including T cells, epithelial cells, endothelial cells and mainly tumor cells while when engaged by PD-1 induces T cell exhaustion, *i.e.* the progressive loss of effector function due to chronic low-affinity antigenic stimulation^84^. The overexpression of PD-L1 in cancer cell lines likely limits the cytotoxic antitumor response of CD8^+^ LT^85^. Here HEV established in MC38 tumors were found associated a lower expression of PD-L1 compared with the PD-L1 intensity labeling measured in B16 and B16-Ova (**Fig. 2b)**. In order to attribute an inflammatory and immunogenic profile to each of the three cancer cell lines, we established an immune scoring system, termed “immunoscore” and “immunosuppressive score” adapted to anti-tumoral/inflamed and pro-tumoral cancers, respectively. By correlating the different intensity levels of immunolabel markers (**Fig. S1a**), we used the positive correlations between CD8^+^ TL, Meca-79^+^ HEV density, CD11b^+^ monocytes and MHCII^+^ TAMs, by attributing for each labeling a score from 0 to 3 corresponding to the [0-25%] to [75-100%] percentiles proving an immunoscore from the sum of each item (**Fig. S1a and Fig. 2c**). In the same way, a negative correlation was evidenced between CD8^+^ TL, Meca-79^+^ HEV and PD-L1 expression, also including the quantification of CD206^+^ TAMs and Gr1^+^CD11b^+^ MDSCs, in the calculation of immunosuppressive score (**Fig. S1a**). Based on these results, density values of MHCII^+^ TAMs, CD11b^+^ monocytes, CD8^+^ lymphocytes and of Meca-79^+^ HEVs were converted into percentiles values, their mean was calculated and converted into an immunoscore (**Fig. 2c, Fig. S1b)**. In the same way, density values of CD206^+^ TAMs, CD11b^+^Gr1^+^ MDSCs and expression of PD-L1 were converted into percentiles values converted into score values from 0 to 3 thus giving an immunosuppressive score (**Fig. 2c, Fig. S1c)**. Consequently, B16 tumors were characterized by the lowest immunoscore (1.400±0.5099) and the highest immunosuppressive score (6.400±0.5099) compared with B16-Ova and MC38. B16-Ova were characterized by an intermediate immunoscore compared with B16 and MC38 exhibiting a similar immunosuppressive score than B16. In contrast, MC38 showed an elevated immunoscore (9.400 ± 0.9798) and a poor immunosuppressive score (2.800±0.4899) compared with B16 (**Fig. 2c**). These data suggest that we can consider B16-derived tumors xenografted in C57B/6 mice is comprised of an immuno-desert TME, B16-Ova of an immuno-excluded TME and MC38 of an immuno-inflamed TME, which enable us to study the role of the different immunogenic status of cancers on animal behavior and brain functions. We then proposed correlation maps of the different behavioral parameters, e.g. preference index for cognitive functions, immobility for resignation, time in open arms and head dips for anxiety, distance for spontaneous activity, and immunescore or immunosuppressive score. The absence of correlation between behavioral results and tumor mass allowed us to exclude the influence of mass volume of animal behaviors (**Fig. 2d**). Globally, except for head dips behaviors, no significant correlations were evidenced between behavioral items and immunosuppressive or immunoscore suggesting that TME inflammatory status do not represent a relevant predictive biomarker for CRCI symptoms (**Fig. 2d**).

**Figure 2.**
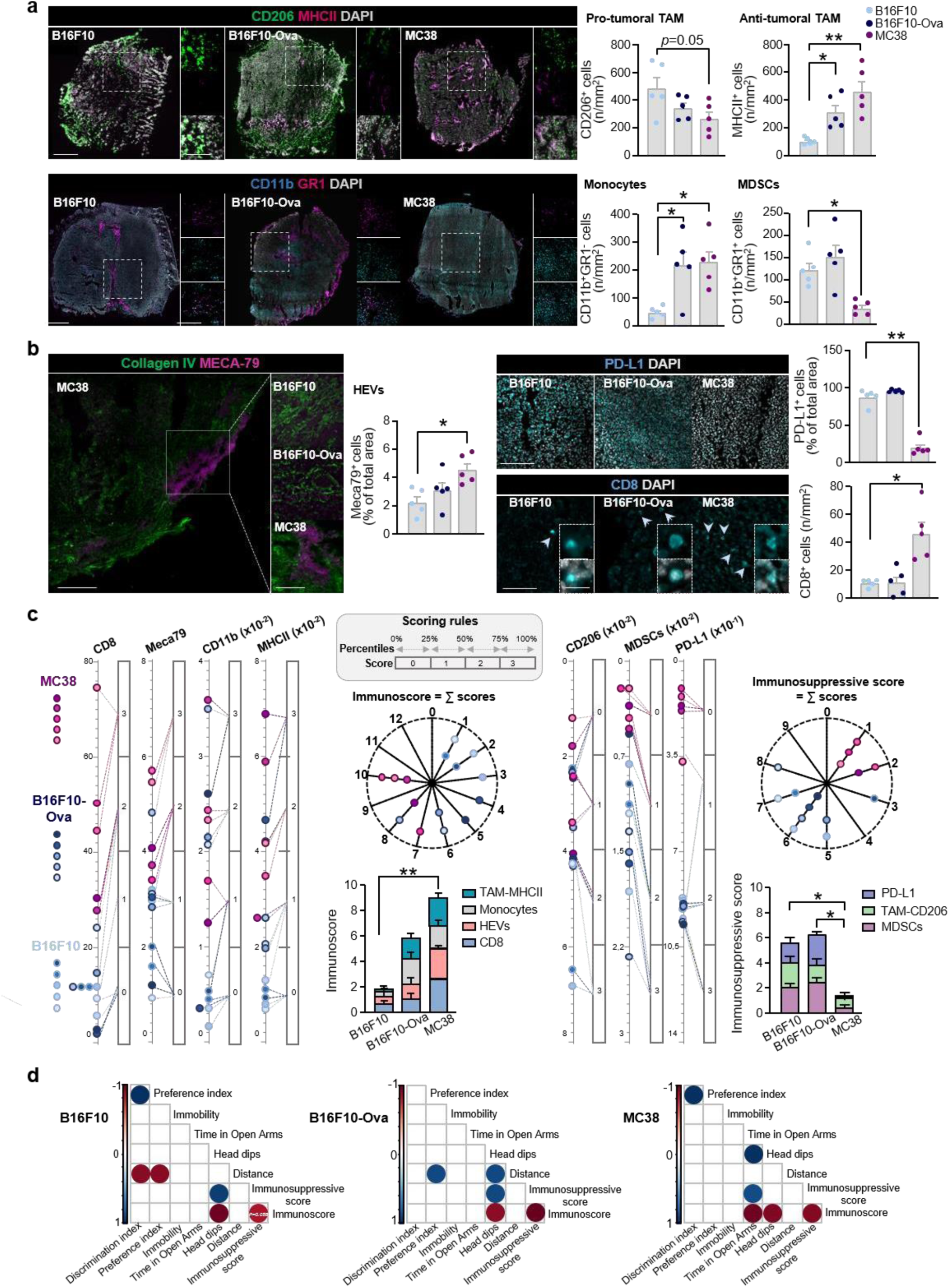
Characterization of immune-desert, immune-excluded and immune-inflamed microenvironment of B16F10, B16F10-Ova and MC38 tumors in C57Bl/6 mice. **a.** In the upper panel, CD206 (green), MHCII (magenta) and DAPI (grey) immunoreactivity in B16, B16-Ova and MC38 tumor slices (30 μm). The boxed areas represent magnification of clusters of MHCII^+^ and CD206^+^ cells. Scale bar: 500 μm and 100 μm. In the lower panel, GR1+ (magenta), CD11b+ (blue) and DAPI (grey) immunoreactivity in B16F10, B16F10-Ova and MC38 tumor slices. The boxed areas represent magnification of clusters of GR1^+^, CD11b^+^ and GR1^+^CD11b^+^ cells. Scale bar: 500 μm and 100 μm. On the right, bar of quantifications of CD206^+^ or MHCII^+^ TAMS, monocytes (CD11b^+^GR1^-^) and MDSCs (CD11b^+^ GR1^+^) density in B16F10, B16F10-Ova and MC38 tumor slices. Scale bars: 1 mm, zoom 100µm. **b.** On the left, representative immunolabeling of collagen IV^+^ (green) and MECA-79^+^ (magenta) vessels in B16F10, B16F10-Ova and MC38 tumor slices. Boxed areas show magnification of HEVs (MECA79^+^CollagenIV^+^). On the left, bars of quantification of MECA-79^+^ coverage, expressed as the percentage of total area, in B16F10, B16F10-Ova and MC38 tumor slices. Scale bars: 200 µm, zoom 100µm. On the right, representative microphotographs of DAPI (grey) and PD-L1 (cyan) immunolabeling in B16F10, B16F10-Ova and MC38 tumor slices with bars of quantification of PD-L1^+^ cells, expressed as percentage of total cells. Below, DAPI (grey), and CD8+ (cyan) immunoreactivity in B16F10, B16F10-Ova and MC38 tumor slices with bars of quantifications of CD8^+^ cytotoxic TL density. The boxed areas represent magnification of DAPI (grey) nuclear staining with CD8+ (cyan) TL overlapping. Scale bars: 50 µm, zoom 10 µm. Statistical analysis was performed by using one-way ANOVA with Bonferroni correction for multiple comparisons (n=5). Data are represented as bars with symbols for individual data points and they are expressed by mean ± SEM, * p<0.05, ** p <0.01, *** p <0.001. **c.** Chart illustrating the immunoscore (left) and the immunosuppressive score (right) calculation methods in B16, B16-Ova and MC38 tumor slices. Densities of CD8^+^ TL, Meca-79^+^ HEVs, CD11b^+^ monocytes and MHCII^+^ anti-tumoral TAMs in tumor slices were converted into percentile values. Percentiles values were then converted into score values from 0 attributed to [0-25%] percentile, to 3 for [75-100%] percentile. The mean of each score was then calculated to generate immunoscore. The staining intensity values of CD206^+^ TAMs, CD11b^+^GR1^+^ MDSCs and percentage of PD-L1^+^ cells in tumor slices were converted into percentile values. Percentiles values were then converted into score values with score 0 for [0-25%] to 3 [75-100%] percentiles. The mean of each score was then calculated to generate immune-suppressive score. **d.** Correlation map between immunoscore, immune-suppressive score and behavioral results. From the heatmap of Kendall correlation coefficients, only significant correlations for each B16F10, B16F10-Ova and MC38 mice are displayed (adjusted p-value < 0.05). Blue circle indicates a positive correlation and red circle, a negative correlation. Color intensity is proportional to the amplitude of the correlation coefficient in a range of 0 to 1/-1.

### Systemic immune microenvironment associated with cognitive alterations and emotional reactivity in mice bearing peripheral cancers

Given the lack of a link between inflammatory status of peripheral solid tumors and mice behaviors, we decided to explore the systemic immune composition in cancer-bearing and lysate-injected mice and the level of 11 inflammatory mediators (IL-17, IL-10, IL-4, INF-γ, TNF-α, IL-6, MCP-1, IL-1α, IL-1β and ICAM-1) was quantified **(Fig. 3a-c)**. Increased levels of MCP-1, IL-6 and ICAM-1 were measured in B16-Ova and MC38-bearing mice, increased levels of IL-17 and decreased levels of IL-1α were measured exclusively in MC38-mice, while levels of TNF-α were found decreased in B16F10-mice and increased in MC38-mice when compared to PBS cancer-free control mice **(Fig. 3b and d)**. Interestingly, these alterations were not observed in lysate injected mice but increased levels of IL-1α were measured in B16F10- and B16F10-OVA-lysate mice while increased levels of IL-1β were measured in mice receiving MC38 lysate compared to PBS-injected mice **(Fig. 3c)**. Because systemic cytokine levels and ICAM-1 were found associated with cognitive complaints, fatigue and depression in cancer patients^7–13^, potential correlations between cytokines levels and behavioral parameters were tested. Among all cancers, we found a negative correlation between IL-6 and ICAM-1 and spontaneous activity (distance crossed), between IL-6 and MCP-1 and anxiety-like behaviors (head dips number), and between IL-6, ICAM-1, MCP-1 and short-term memory (preference index) (**Fig 3e, f, h**).

**Figure 3.**
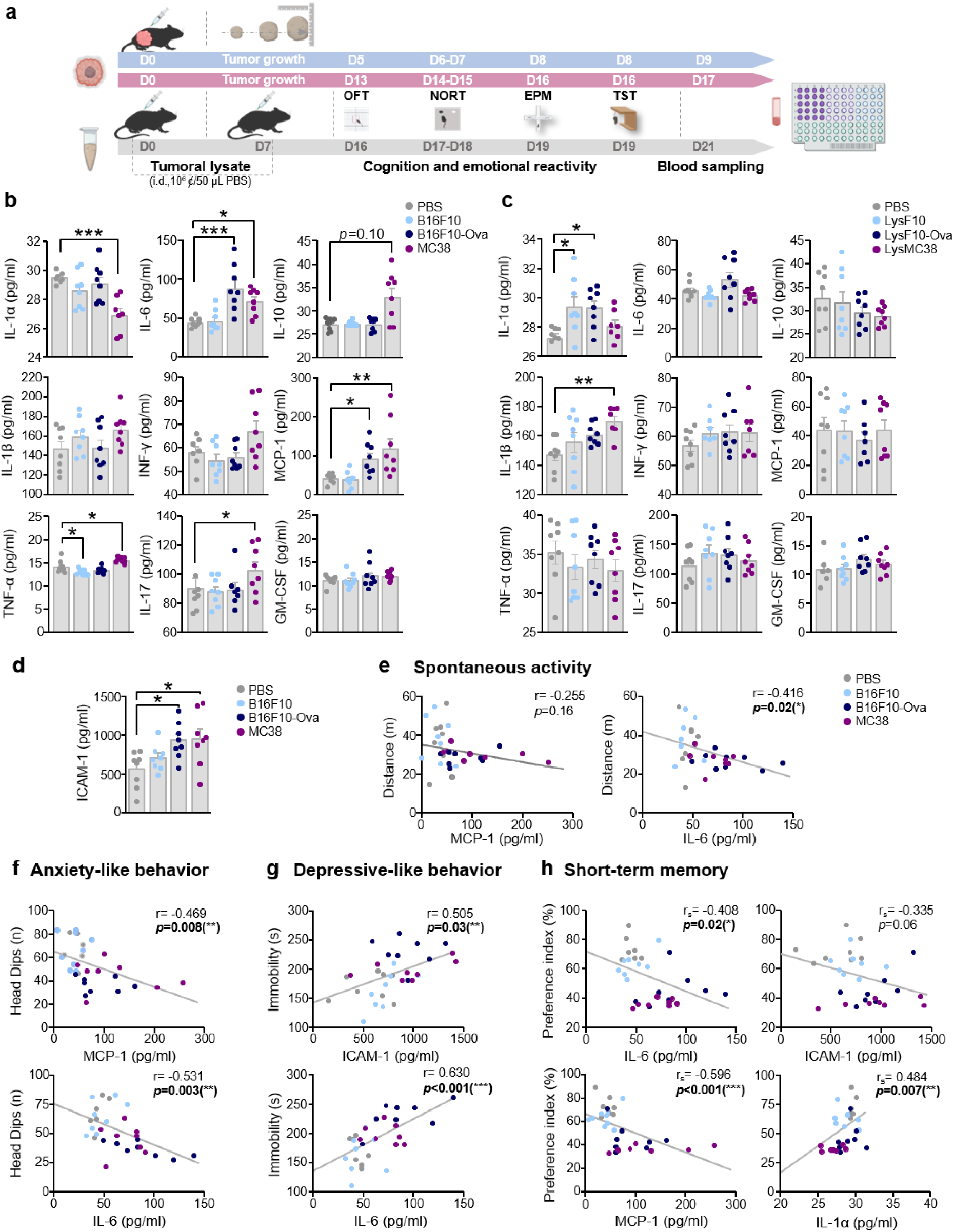
Systemic plasma inflammatory markers in B16F10, B16F10-Ova and MC38 tumor-bearing mice and correlation with behaviors. **a.** Experimental timeline of tumoral cell lines B16- (blue), B16-Ova (blue) and MC38 (purple) or their respective tumoral lysates inoculation and behavioral assessment in mice. Mice were sacrificed the day after the end of the behavioral session and blood was collected. **b.** Histograms levels of various plasma cytokines from the CYTOX score from PBS-mice compared with B16F10-, B16F10-Ova and MC38-bearing mice (n=7). Statistical analyses were represented by using one-way ANOVA with Bonferroni correction for multiple comparisons. Data are represented as bard with symbols for individual data points and they are expressed by mean ± SEM, * p<0.05, *** p<0.001. **e-h**, Regression curves showing correlation between head dips (anxiety), immobility (resignation), and preference index (cognition), and some cytokines or ICAM-1 in PBS mice and B16F10, B16F10-Ova and MC38-bearing mice (n=7). Statistical analyses were performed by using Spearman correlation. D: day, i.d.: intradermally, EPM: elevated plus maze, GM-CSF: Granulocyte Macrophage Colony-Stimulating Factor, ICAM-1: Intercellular Adhesion Molecule 1, IL: interleukin, INF: interferon, Lys: lysate, MCP-1: Monocyte Chemotactic Protein-1, NORT: novel object recognition test, OFT: open field test, TST: tail suspension test, TNF: Tumor Necrosis Factor.

Inversely, positive correlations were found between plasma levels of IL-1α and short-term memory and of IL-6 and ICAM-1 and resignation-like behaviors (immobility) (**Fig. 3g and h**). All together, these results suggest that systemic immune mediators act as potential relays of the impact of immune-desert, immune-excluded, immune-inflamed TME and respective tumoral lysates on cognitive functions and emotional reactivity.

### Peripheral cancers induce blood-CSF barrier inflammation favorizing myeloid brain-infiltration which supports ventriculomegaly and neuroinflammation

To further study how systemic inflammatory mediators can impact brain functions, we first focused on CNS-barriers such as the blood-cerebrospinal fluid barrier (BCSFB). Indeed, systemic cytokines such as IL-1β or TNF-α have already been demonstrated to enhance the expression of adhesion molecules such as ICAM-1 and MAdCAM-1 in choroid plexus (CP)^14,15,86^, thus favorizing leukocyte adhesion and trafficking across the BCSFB^87^. In B16F10- and MC38-bearing mice, increased ICAM-1 and MadCam-1 coverage of Lectin^+^ CP areas as well as higher lateral ventricle volumes (B16: 0.3679±0.02786; MC38: 1.151±0.2422) were observed compared with PBS-mice, highlighting an original role of either immune-desert or immune-inflamed cancer on BCSFB inflammation and CSF distribution (**Fig. 4a, S2a)**. To further verify if upregulation of adhesion molecules in the CP was associated with homing and migration of leukocytes, we analyzed the number of F4/80^+^ macrophages and CD3^+^ TLs in CP and CSF-filled spaces such as perivascular spaces and leptomeninges by immunohistochemistry. Increased number of F4/80^+^ macrophages was observed in CP and in leptomeninges of MC38-bearing mice compared with PBS-mice (**Fig. S2b**). No alteration in the number of CD3^+^ TLs was found in CP (**Fig. S2c**) and perivascular spaces of Lectin^+^ hippocampal vessels of cancer-bearing mice, but increased number of CD3^+^ TLs was found in leptomeninges of MC38-mice (**Fig. S2d-f**), mice (**Fig. 4b**). Depletion of macrophages was confirmed by analyzing Cd11b^+^F4/80^+^ macrophages in blood using flow cytometry (**Fig. 4b**).

**Figure 4.**
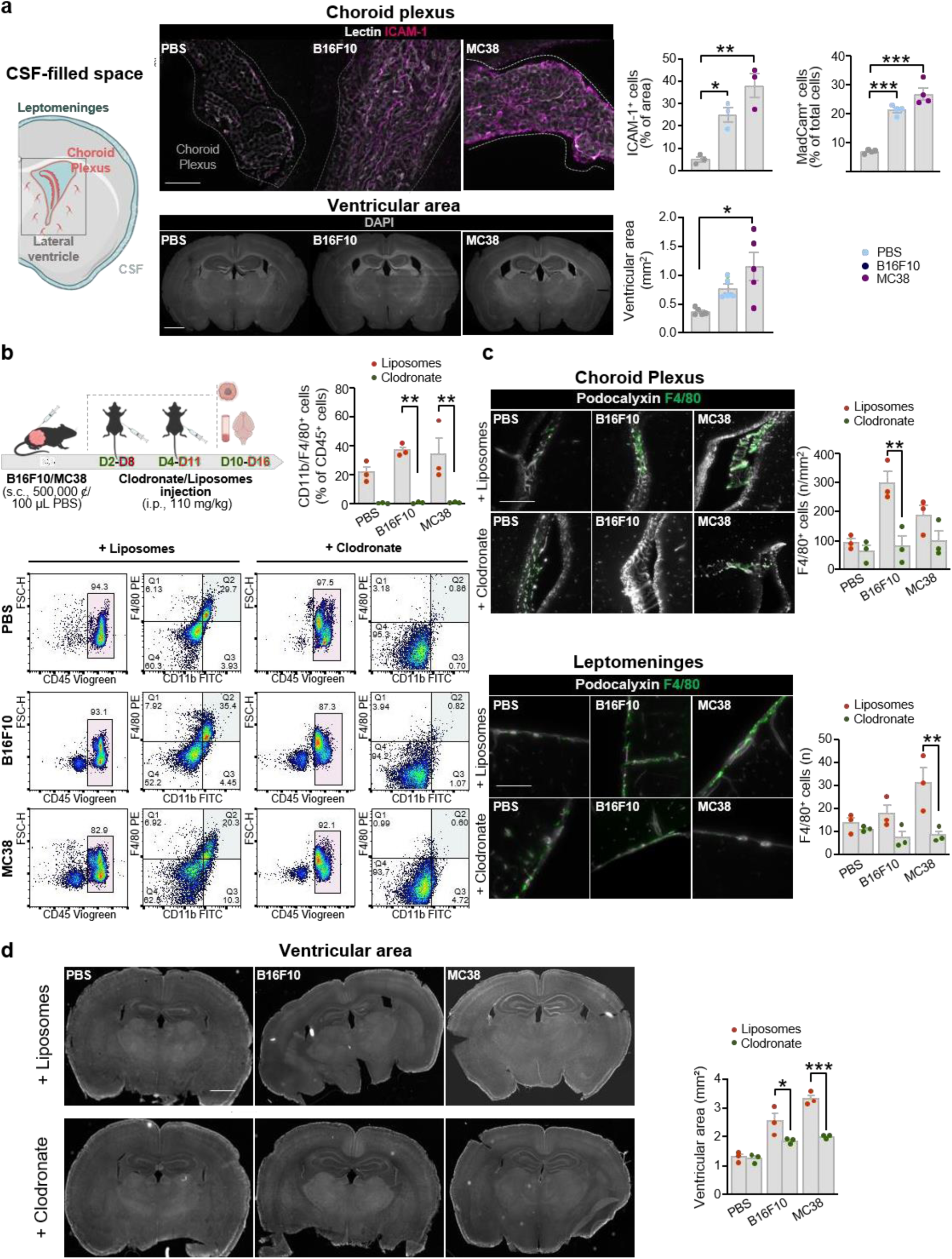
Vascular inflammation, ventriculomegaly and macrophages infiltration in choroid plexus and leptomeninges in cancer-bearing mice. **a.** On the left, schematic representation of CSF circulation from CP to leptomeningeal compartment. In the middle, representative images of ICAM-1+ (magenta) and Lectin+ (grey) immunoreactivity in CP of PBS-control or B16F10- and MC38-bearing mice. Scale bar: 50 µm. Below, representative images of MadCam-1+ (green) and Lectin+ (grey) immunoreactivity in CP of PBS-control or B16F10- and MC38-bearing mice. Scale bar: 100 µm. On the right, representative images of nuclei staining by DAPI (grey) immunostaining on cerebral slices of PBS or B16F10- and MC38-bearing mice. Scale bar: 1 mm. Below, bars of quantification of ICAM-1, expressed as percentage of total area, of MadCam-1^+^ cells, expressed as percentage of total cells and of the ventricular volumes (3rd and lateral ventricles) of B16- and MC38-bearing mice compared with PBS-control mice. Data are represented as bars with symbols for individual data points and expressed by mean ± SEM, n=3-4, * p<0.05, ** p <0.01, *** p < 0.001. **b.** Schematic timeline for *in vivo* macrophage depletion assay in B16- and MC38-bearing mice. After tumors inoculation (D0), mice received two intraperitoneal injections (i.p.) of 100 µl of Clodronate/Liposomes (110 mg/Kg) at day 2 and day 4 for B16F10-bearing mice and at day 8 and day 11 for MC38-bearing mice. At day 10 and at day 16, B16F10- and MC38-bearing mice respectively were sacrificed, and blood and brain were collected. On the right, bars of FACS analysis quantification of CD11b^+^F4/80^+^ cells, expressed in percentage of CD45^+^ leukocytes, in PBS-, B16F10- and MC38-mice receiving liposomes (orange) and clodronate (green) (n=3). Statistical analyses using Mann-Whitney test; Data are represented as bars with symbols for individual data points. Data are expressed by mean ± SEM, ** p <0.01. Below, representative FACS profiles of PBMC from PBS-, B16F10- and MC38-bearing mice treated with liposomes (left) and clodronate (right). Peripheral blood cells were stained with CD45-Viogreen; F4/80-PE and CD11b-FITC antibodies. **c**. Representative images of macrophages (F4/80^+^, green) in CP (Podocalyxin+, grey) and leptomeninges (Podocalyxin+, grey) in PBS-, B16F10-, MC38-bearing mice treated with liposomes (upper panel) and clodronate (lower panel). On the right, Quantification bars of 4/80^+^ staining density in CP and meninges (n=3). Statistical analyses using Mann-Whitney test; Data are represented as bars with symbols for individual data points. Data are expressed by mean ± SEM, ** p<0.01. Scale bar: 50 µm. **d**. Representative images of DAPI immunoreactivity on whole brain coronal section of PBS-, B16F10- and MC38-bearing mice treated with liposomes (left) or clodronate (right). On the right, quantification of ventricular volumes (3^rd^ ventricles and lateral ventricles) of PBS-, B16- and MC38-bearing mice treated with liposomes (pink) or clodronate (blue) (n=3). Statistical analyses using Mann-Whitney test; Data are expressed by mean ± SEM, * p< 0.05, *** p<0.001. Scale bar: 1mm.

Suggesting that myeloid and lymphocytic CSF-infiltration occurs exclusively in presence of immune-inflamed cancer. To verify the origin of CSF-homing F4/80^+^ macrophages, we depleted systemic macrophages by means of clodronate-liposomes administration once a week during two weeks by i.p. injection to PBS, B16 or MC38 In cancer-bearing mice receiving clodronate, within the CP and leptomeningeal Lectin^+^ areas, the number of F4/80^+^ macrophages was normalized to the number observed in PBS-cancer-free controls, thus suggesting that the increased number of F4/80^+^ cells likely result from infiltration of peripheral macrophages (**Fig. 4c**). As an increment in the amount of CSF, either due to excessive production or reduced reabsorption, is frequently associated with altered expression of the AQP4 present at the glia limitans of parenchymal vessels and in the ependymal cells lining the ventricular system, we first verified if loss of AQP4 in these areas was at the origin of ventriculomegaly observed in B16- and MC38-bearing mice^16^ However, a significant decrease in AQP4^+^ staining intensity was observed in endfeet of perivascular GFAP^+^ astrocytes exclusively in B16-mice (**Fig. S3**), suggesting that diminution of ACQ4 expression only partially contributes to CSF accumulation observed in cancer-bearing mice. However, as a key role of CP-macrophages has been demonstrated in pathophysiology of hydrocephaly^17^, we decided to study if macrophagic infiltration occurring in CP of cancer-bearing mice significantly contributes to ventriculomegaly. Interestingly, systemic depletion of CD11b^+^F4/80^+^ macrophages by clodronate administration prevented ventricular volume enlargement in cancer-bearing mice (**Fig. 4d**), suggesting that BCSFB perturbation in presence of immune-inflamed cancer can lead to peripheral macrophages infiltration and subsequent CSF accumulation in ventricles, potentially underlying exacerbated behavioral alterations observed in MC38-bearing mice.

During peripheral and central inflammation, leukocytes infiltration may occur in other CSF-filled compartments such as the *velum interpositum* (VI), in proximity of the hippocampal area^18^ (**Fig. 5a**). In our study, the presence of F4/80^+^ macrophages was investigated within the VI ERTR7^+^ and perivascular space of DG Lectin^+^ vessels (**Fig. 5a**). Interestingly, while no F4/80^+^ macrophages were found in the DG of all cancer and cancer-free control mice, a significantly higher density of F4/80^+^ macrophages was measured in the VI of MC38-bearing mice compared with B16F10- and PBS-control mice (**Fig. 5a**). Moreover, immunohistochemical analysis revealed a higher density of CD206^+^ (M2-like) pro-tumoral phenotypic macrophages in VI of MC38-bearing mice compared with B16F10 or PBS mice, with no significant different in density of anti-tumoral MHCII^+^ macrophages (**Fig. 5b**). To study whether this myeloid infiltration proximal to the hippocampus could impact hippocampal neurogenesis, thus contributing to the alteration of cognitive processes, cancer and control mice were injected with BrdU two days before sacrifice, with the aim to detect the number of dividing neural precursor cells (NPC) (**Fig. 5c**). Within the DG, a decrease in number of BrdU^+^ cells (green) lying the granular layer NeuN^+^ (grey) was observed in B16- and MC38-mice compared with PBS-mice, suggesting a direct impact of cancer on neurogenesis. As microglia are the caretakers of the neurogenic niche, constantly scanning the microenvironment and when activated, regulating progenitors proliferation^19^, we also tested their inflammatory status by means of the CD68 marker, and showed that density of CD68^+^Iba1^+^ reactive microglia increased, and of Iba1^+^Arg1^+^ anti-inflammatory microglia decreased in the DG of B16F10- and MC38-bearing mice compared with PBS-control mice (**Fig. 5d and Fig. S4a**).

**Figure 5.**
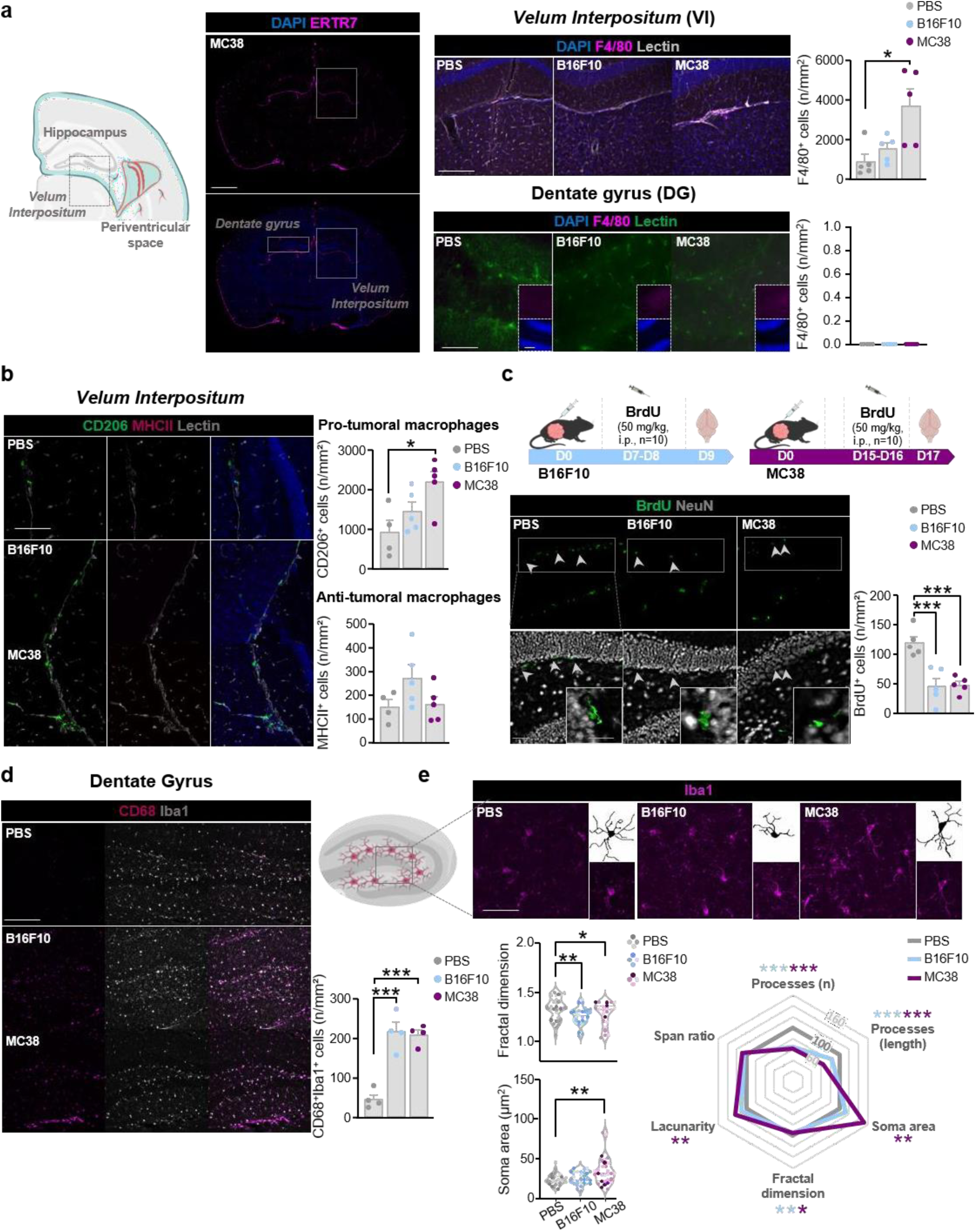
Hippocampal inflammation and impaired neurogenesis in cancer-bearing mice. **a.** On the left, schematic representation of *velum interpositum* localized below the hippocampus and near lateral ventricles in mouse brain. In the middle, representative images of nuclei staining by DAPI (grey) and leptomeninges (ERTR7^+^, magenta) immunostaining showing positions of dentate gyrus and ERTR7^+^ *velum interpositum* on whole cerebral slices of MC38-bearing mice. Scale bar: 1 mm. On the right, representative micrographs and statistical quantification of macrophages (F4/80^+^, magenta) density (n/mm^2^) and nuclei (DAPI, blue) immunolabeling in *velum interpositum* (Lectin^+^, grey) and dentate gyrus vessels (Lectin^+^, green) of PBS and B16F10- and MC38-bearing mice. Scale bars: 100 µm. **b.** Representative images and quantification of pro-tumoral macrophages (CD206^+^, green) and anti-tumoral (MHCII^+^, magenta) immunolabeling in *velum interpositum* (Lectin^+^, grey) of PBS and B16F10- and MC38-bearing mice. Scale bar: 100 µm. **c.** Schematic diagram showing the timeline for bromodeoxyuridine (BrdU) administration in B16F10- (light blue) and MC38- (pink) bearing mice. After tumors inoculation (D0), B16F10 and MC38 mice received BrdU (50 mg/kg) by intraperitoneal injection (i.p.) once a day for 2 days before sacrifice. At D9 (B16F10) or D17 (MC38) mice were sacrificed, and brain were collected for immunofluorescence assays. Below, representative micrographs of neuronal precursors (BrdU^+^, green) and mature neurons (NeuN^+^, grey) in the dentate gyrus of the hippocampus of PBS and B16F10-, MC38-bearing mice. The white squares below the images correspond to magnification showing absence of co-localization of neuronal precursors (BrdU^+^, green) with mature neurons (NeuN^+^, grey) in the dentate gyrus (DG). On the right, histogram showing the variation in the number of BrdU^+^ neuronal precursors (density, n/mm^2^) in the dentate gyrus of PBS mice compared with B16F10- and MC38-bearing mice. Scale bar: 100 µm, zoom: 10 µm. **d.** Representative images of reactive (CD68^+^, magenta) microglia (Iba1^+^, grey) immunolabeling in dentate gyrus of PBS-mice and B16F10- and MC38-bearing mice. On the right, histogram showing the variation in the number of CD68^+^Iba1^+^ reactive microglial cells (density, n/mm^2^) in dentate gyrus of B16F10- and MC38-bearing mice compared to PBS-mice. Scale bar: 100 µm. Statistical analyses were performed by using one-way ANOVA or Kruskal-Wallis test with Bonferroni or Dunn’s correction for multiple comparisons. Data are represented as bars with symbols for individual data points and they are expressed by mean ± SEM, n=4-5, * p<0.05, *** p <0.001. **f**. Representative example of photomicrographs used for the morphological analysis of microglial cells (Iba1^+^, magenta). The boxed areas represent magnification of isolated Iba1^+^ microglial cells (bottom) and corresponding binary image (top) from the dentate gyrus of PBS, B16F10- and MC38-bearing mice. Analysis of the morphology of the microglial cells was performed on 8 cells of dentate gyrus of PBS-, B16F10- and MC38-bearing mice (n=4). Scale bar: 50 μm, zoom 10 μm. Below, quantification of fractal dimension coefficient and area of the soma of microglial cells of PBS-, B16F10- and MC38-bearing mice. Data are represented as violin-plots with individual points representing individual microglial cells (n of cells=32, n of mice = 4 for experimental group). On the right, comparative Radar Plot Chart of morphological parameters (number of processes, length of processes, area of the soma, fractal dimension, lacunarity and span ratio) analyzed on microglia cells in dentate gyrus of PBS-, B16F10- and MC38-bearing mice. Statistical analyses were performed by using one-way ANOVA or Kruskal-Wallis test with Bonferroni or Dunn’s correction for multiple comparisons, * p < 0.05, ** p <0.01, *** p <0.001. BrdU: Bromodeoxyuridine, NeuN: neuronal nuclei antigen, CD206: cluster of differentiation 206, CD68: cluster of differentiation, DAPI: 4’,6-diamidino-2-phenylindol, Iba-1: ionized calcium-binding adapter molecule 1, MHCII: major histocompatibility complex 2.

In a good agreement, by means of microglial morphologic analysis^20^, we measured increased inflammatory morphological indicators such as diminished fractal dimension, length and number cells processes in microglial cells of B16- and MC38-mice DG compared to cancer-free PBS-mice, and increased soma area exclusively in MC38-mice, supporting an exacerbated inflammatory profile of microglia in DG of MC38-mice (**Fig. 5e, S4b**). However, within the DG in particular, the quantification of the GFAP intensity signals as a marker of astrogliosis failed to show any changes between cancer-bearing mice and PBS control mice (**Fig. S4c**). Altogether, these results evidence that BCSFB perturbations can favorize peripheral-to-brain myeloid infiltration in proximity of cognitive-relevant brain areas such as hippocampus leading to hippocampal inflammation and altered neurogenesis potentially underlying behavioral alterations in cancer-bearing mice.

### Myeloid cells infiltration in perivascular spaces is supported by circumventricular organs, permeability of the brain blood barrier and specific meningeal dura mater inflammation in cancer-bearing mice

Similarly to CP, circumventricular organs (CVOs) such as median eminence, lack endothelial BBB and have been recently identified has the main cerebral entry site of peripheral myeloid cells in context of peripheral cancer^21^. In MC38-mice, a significant accumulation of both anti-tumoral MHCII^+^ and pro-tumoral CD206^+^ myeloid cells was found within Laminin^+^ vessels when compared to PBS control mice (**Fig. S5a**). In parallel, significant accumulation of perivascular pro-tumoral CD206^+^ myeloid cells was found in proximity of Laminin^+^ vessels in the Willis circle and parenchyma of MC38-bearing mice compared to PBS cancer-free controls (**Fig. 6a**), indicating that CVOs, beyond CPs, could represent a relevant entry site for myeloid cells colonizing CSF-filled perivascular spaces in presence of a peripheral immune-inflamed cancer. As perivascular macrophages accumulation has been demonstrated to directly impact BBB permeability^22^, we verified BBB integrity in our context by injecting a dextran-FITC tracer (3kD, i.p.) 15 min before sacrifice to cancer-bearing and cancer-free mice (**Fig. 6b**). After blood and brain harvesting following intra-cardiac perfusion, a ratio of fluorescence in plasma and hemi-brain lysates was used to calculate extra-vascular dextran diffusion in PBS-, B16F10 and MC38 mice. A significant increase of the permeability index, indicative of BBB leakage, was observed in both B16F10- and MC38-bearing mice compared with PBS-mice (**Fig. 6b**). From cerebral slices, extravascular Dextran_488_ staining (green) was found absent in podocalyxin (endothelial marker)^+^ hippocampal vessels of PBS-, B16F10 or MC38 mice, detectable in proximity of vessels of cerebellum, frontal and somato-motor cortex of B16F10 and MC38 mice and not of PBS-mice (**Fig. 6c**), and distributed diffusible in the extra-vascular compartment of the whole-mount meningeal dura matter, exclusively in MC38 mice **(Fig. 6c**), suggesting that endothelium-macrophage crosstalk is at least in part, but not totally, responsible of BBB integrity perturbation.

**Figure 6.**
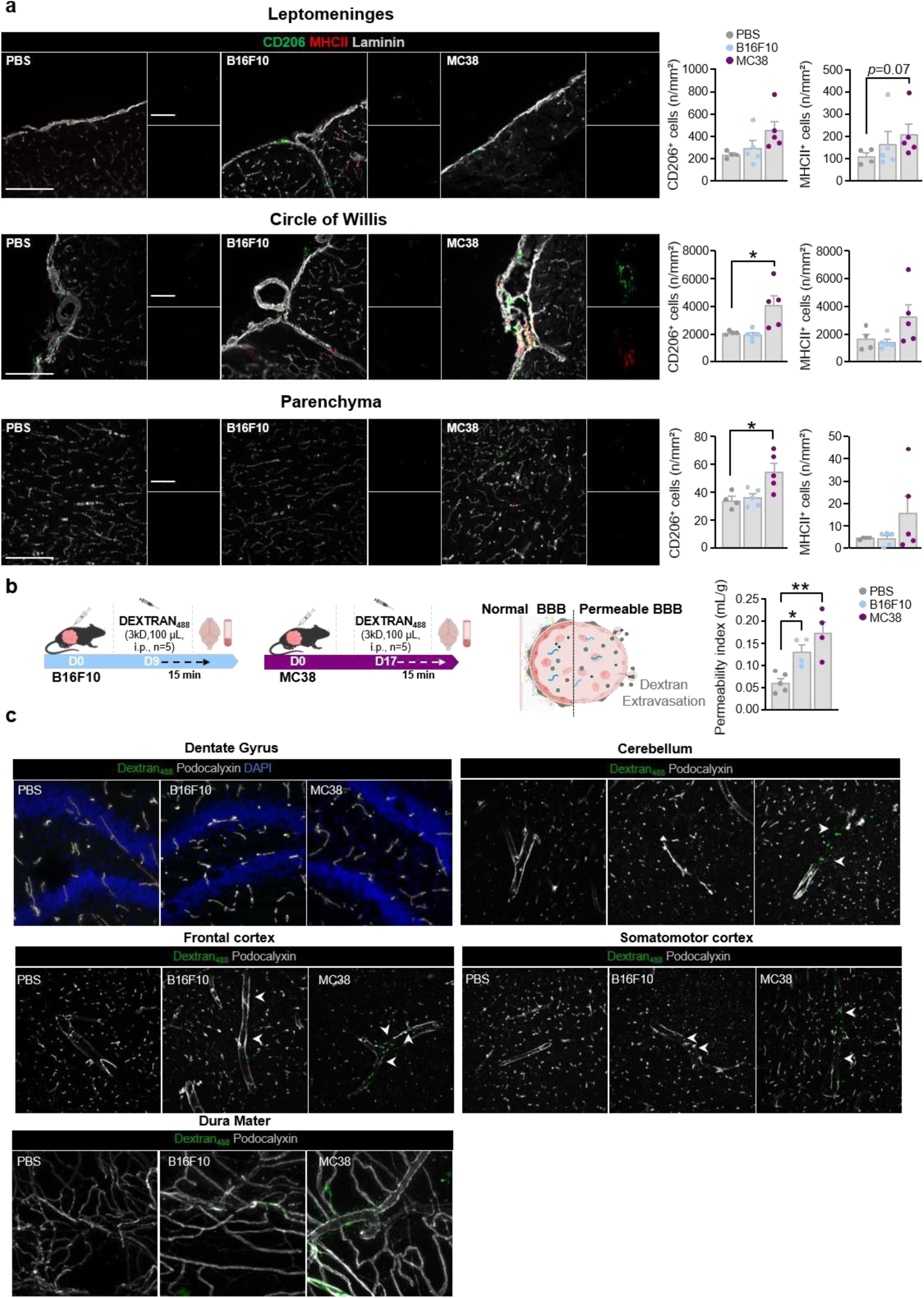
Peripheral cancers are associated to macrophages invasion of cerebral areas and blood-brain barrier permeability. Representative images and statistical quantification of density (n/mm^2^) of pro-tumoral (CD206^+^) and anti-tumoral (MHCII^+^) myeloid cells staining in Laminin^+^ (grey) leptomeninges (upper panel), Willis polygon (middle panel) and perivascular space of parenchyma vessels (lower panel) of B16F10- and MC38-bearing mice compared to PBS-controls. Scale bar: 100 µm. Statistical analyses were performed by using one-way ANOVA or Kruskal-Wallis test with Bonferroni or Dunn’s correction for multiple comparisons. Data are represented as bars with symbols for individual data points and they are expressed by mean ± SEM, n=4-5, * p<0.05. **b.** Schematic diagram showing the timeline of the *in vivo* assay of blood brain barrier permeability in B16- (blue) and MC38- (pink) bearing mice. After tumor cells inoculation (D0), mice received 100 µl of Dextran_488_ (3kD, i.p) at D9 for B16-bearing mice and at D17 for MC38-bearing mice. Post-injection (15 min), after mouse sacrifice by intracardiac perfusion, brains and blood were collected. The permeability index represents the fluorescence signal of cerebral tissue normalized to the weight of the tissue and to the plasma fluorescence signal. On the left, bars represent statistical comparison of permeability index of B16- and MC38-mice compared with PBS mice. Statistical analyses were performed by using one-way ANOVA test with Bonferroni correction for multiple comparisons. Data are represented as bars with symbols for individual data points and they are expressed by mean ± SEM, n=4-5, * p<0.05, ** p <0.01. **c.** Representative micrographs showing Dextran_488_ (green) extravasation from Podocalyxin^+^ (grey) vessels in the DC (upper panel, left), cerebellum (upper panel right), frontal cortex (lower panel, left) and somatomotor cortex (lower panel, right), and Meningeal dura mater (lower left panel) of PBS and B16- and MC38-bearing mice. Scale bar: 50 μm. CD206: cluster of differentiation 206, D: day, i.p.: intraperitoneal injection, MHCII: major histocompatibility complex 2.

Among CSF-filled compartments, dura mater meningeal inflammation has been demonstrated to play a key role in physiological control of short-term memory processes and anxiety-like behaviors^23^. For this reason, we next analyzed the impact of MC38 or B16 cancer on the expression of 48 genes targeting inflammation in the dura mater meningeal layer of PBS, B16- and MC38-mice (**Fig. 7a**). We thus highlighted increased mRNA levels in dura mater layer of chitinase-3-like-1 (Chi3l) and CD19 from B16 mice compared with PBS mice on the one hand, and of the triggering receptor expressed on myeloid cells 2 (TREM2), CD80, CD4 and the Toll-like receptor 2 (TLR2) in MC38 mice compared with PBS-mice, on the other hand. Moreover, the mRNA level of the Integrin alpha M (CD11b) as well as of the Stimulator of Interferon Genes (STING) was enhanced in both B16 and MC38-bearing mice compared with PBS mice (**Fig. 7b**). An intent to integrate these differentially expressed genes was done through the Ingenuity Pathway Analysis (IPA) software showing specific interaction maps for B16 and MC38 (**Fig. 7c**). Thus, dura layer mRNA levels upregulated in B16 mice may be involved in five canonical pathways such as the ‘Systemic lupus Erythematosus in B Cell Signaling Pathway’ found to be the highest-ranking signaling pathway, and in MC38 they may be involved in five canonical pathways associated with the “Toll-like receptor cascade” given with the highest rank (**Fig. 7c, Fig. S6**). All together these data suggest that behavioral alterations observed in presence of peripheral cancers could be sustained by CNS barriers permeability and inflammation leading to dural and hippocampal inflammation, susceptible to be exacerbated by additional anti-cancer treatments.

**Figure 7.**
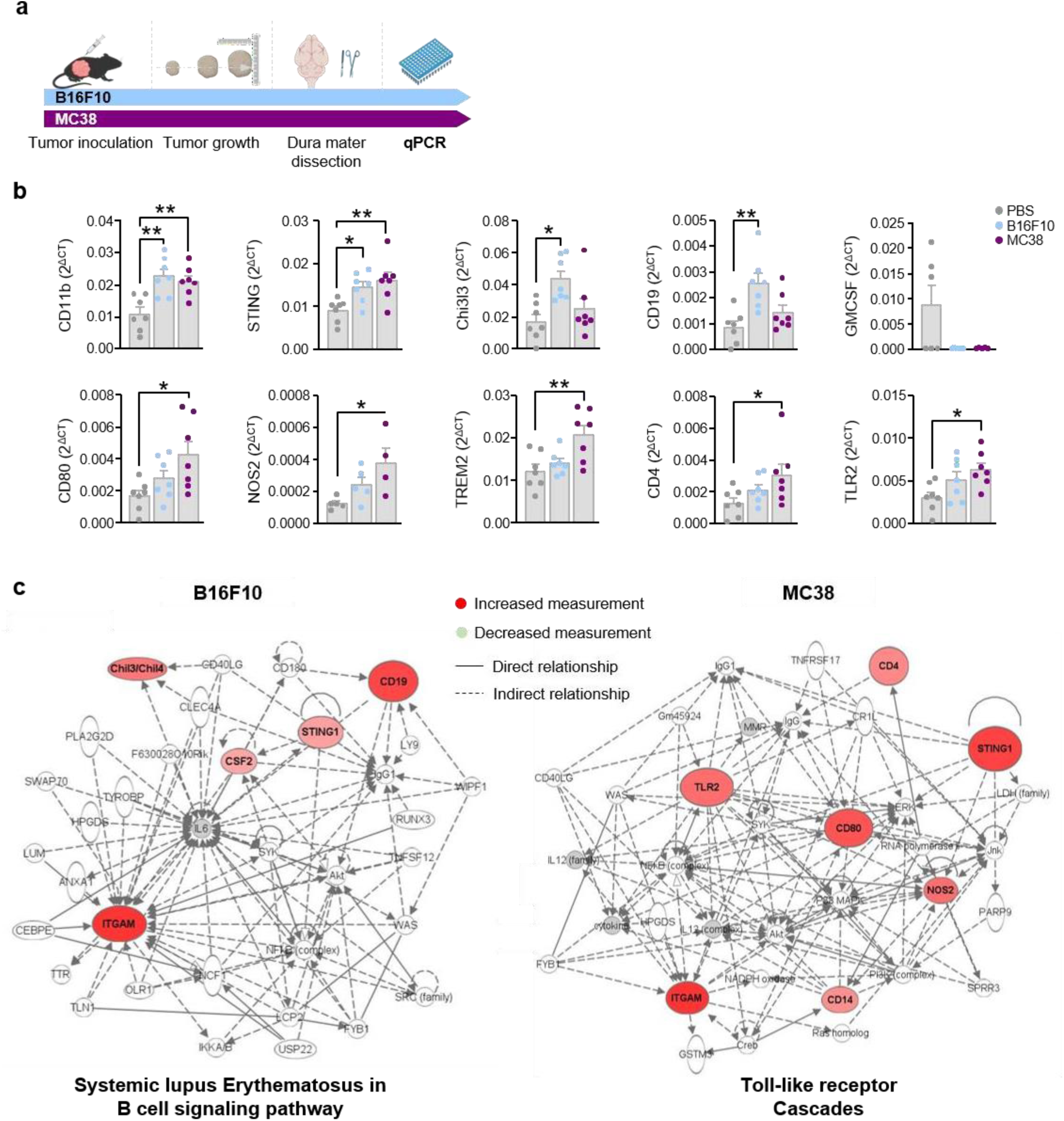
Peripheral cancers are associated with inflammation in the Dura Mater meningeal layer. **a.** Schematic diagram showing the timeline for dura mater isolation and qPCR analysis of inflammatory genes transcripts (n=48) in B16F10- (light blue) and MC38- (pink) bearing mice. **b.** Bars representing the statistical comparison of mRNA transcript levels expression of CD11b, STING, Chi3l3, CD19, GM-CSF, CD80, Nos2, CD4, TLR2 and TREM 2 in meningeal dura mater isolated from PBS, and B16- and MC38-bearing mice. Data are represented as bars with symbols for individual data points and they are expressed by mean ± SEM, n=7. Statistical analyses were performed by using one-way ANOVA or Kruskal-Wallis test with Bonferroni or Dunn’s correction for multiple comparisons, * p<0.05, ** p <0.01. **c**. Gene interaction network maps generated by the Ingenuity Pathway Analysis (IPA) software representing one of the five canonical significant pathways from B16 (left) and MC38 (right) mice. CD11b or ITGAM: beta 2 integrin adhesion molecule, Chi3l3: chitinase-like 3, CD: cluster of differentiation, GM-CSF or CSF2: Granulocyte-macrophage colony-stimulating factor, Nos2: nitric oxide synthase 2, STING: Stimulator of interferon genes, TLR: Toll-like receptor 2, TREM2: Triggering Receptor Expressed on Myeloid Cells 2.

### Immune-check point inhibitors differentially exacerbate cancer-induced emotional reactivity and cognitive functions

We next investigated the impact of ICIs administration on the behaviors of mice bearing B16- and MC38-and treated either by three injections (D2, D4, D7 for B16 and D6, D10, D13 for MC38) of anti-PD-1, anti-PD-L1 or IgG2a before behavioral assessment (**Fig. 8a**). Tumor growth was significantly delayed in MC38-PD1 and MC38-PD-L1 mice compared with MC38-IgG2a mice, while no significant tumoral growth kinetic between B16-PD1 and B16-PD-L1 mice compared with B16-IgG mice (**Fig. 8b**).

**Figure 8.**
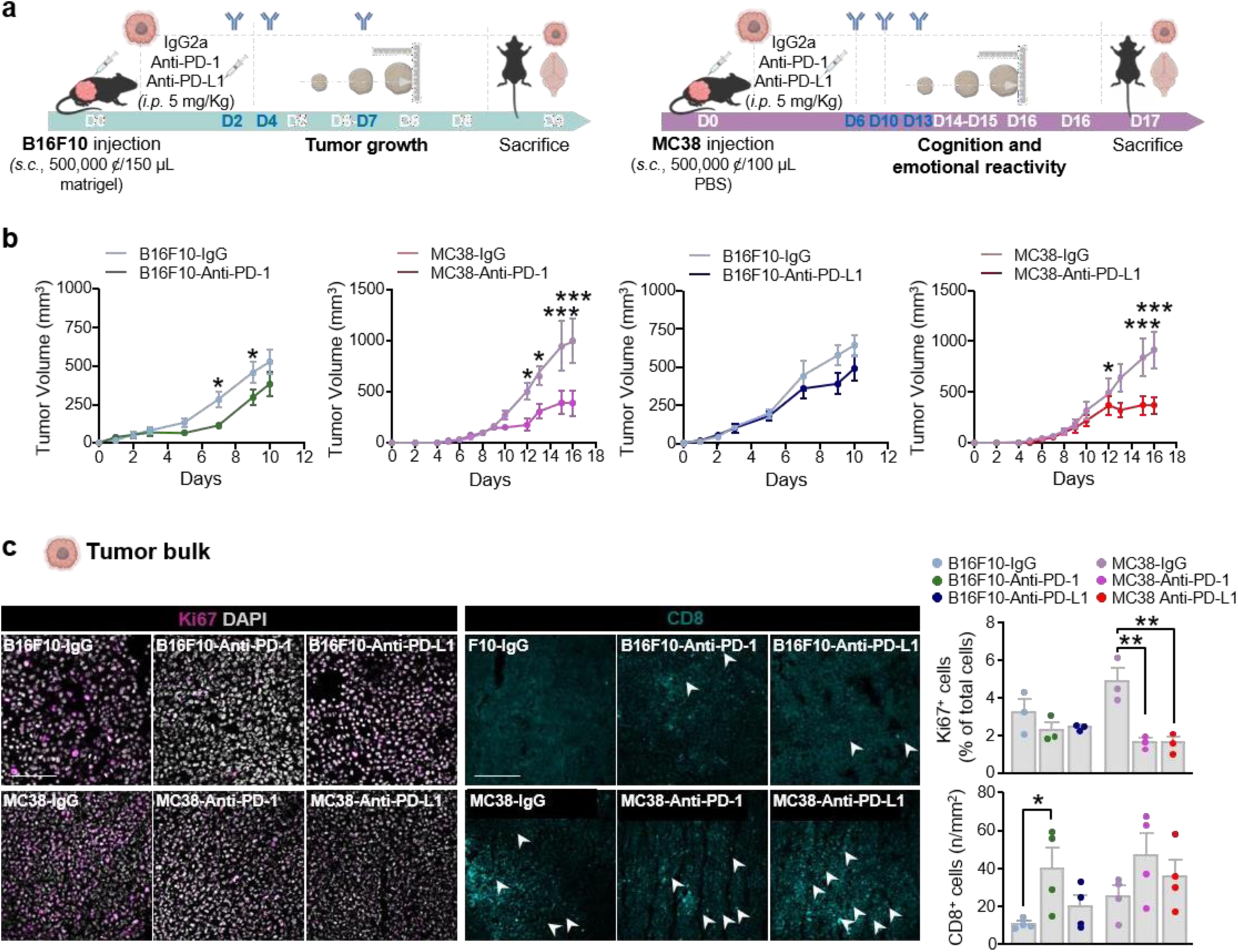
Impact of anti-PD-1 or anti-PD-L1 on tumor growth. **a.** Schematic timeline of anti-PD-1, anti-PD-L1 and IgG administrations in B16- (green) and MC38- (purple) bearing mice. B16F10 mice were injected (i.p.) with of anti-PD1, anti-PD-L1 or IgG at D2, D4 and D7. MC38 mice were injected (i.p.) with of anti-PD1, anti-PD-L1 or IgG at D6, D10 and D13 and tumor growth was measured by means of a caliper. Mice were sacrificed (B16F10: D9, MC38: D17) and brain, blood and tumors were collected for further analysis. **b.** Curves showing tumor volume as mean ± SEM (n = 9-10) of B16-IgG (light green), B16F10-PD1 (middle green) and of MC38-IgG (light pink), MC38-PD1 (middle pink) at indicated time points. On the right, curves of tumor volume progression represented as mean ± SEM (n = 10) of B16F10-IgG (light green) and B16F10-PD-L1 (dark green) and of MC38-IgG (light pink) and MC38-PD-L1 (dark pink) at indicated time points. Statistical comparison was performed by two-way ANOVA test with Bonferroni correction for multiple comparisons. * p<0.05, *** p <0.001. **c.** Representative immunofluorescence of proliferating (Ki67^+,^ magenta) cells (DAPI^+^, grey) and cytotoxic T lymphocytes (CD8^+^, cyan) in tumors slices of B16F10- and MC38-bearing mice treated with IgG, anti-PD1 and anti-PD-L1. On the right, bars representing statistical comparison of KI67^+^ proliferating cells, expressed as the percentage of total cells, and the density (n/mm^2^) of CD8^+^ cytotoxic T lymphocytes in tumors slices of B16F10- and MC38-bearing mice treated with anti-PD1 and anti-PD-L1compared to respective IgG-treated controls. Statistical analysis was performed by One-way ANOVA with Bonferroni correction for multiple comparisons (B16F10-IgG vs B16F10-PD1 or B16F10-PD-L1, MC38IgG vs MC38-PD-1 or MC38-PD-L1). Data are represented as bars with symbols for individual data points and they are expressed by mean ± SEM, n=3-4, * p<0.05, ** p <0.01. Scale bars: 100 μm. CD: cluster of differentiation, D: day, i.p.: intraperitoneal injection, DAPI: 4’,6-diamidino-2-phenylindol, Ki67: Antigen Kiel 67.

This indicates that the presence of immuno-inflamed MC38 cancer is associated with a better anti-tumoral response to ICIs than immuno-desert B16 cancer. The ICI-induced anti-tumor response was evidenced by the significant decreased number of Ki67^+^ proliferative cells exclusively in MC38 tumor bulk even if the number of CD8^+^ TLs was not modified within the B16 and MC38 tumors, in the absence or the presence of ICIs (**Fig. 8c**), suggesting that possible polarization of CD8 toward a proliferative/effector phenotype could explain the increased anti-tumoral efficacy of ICIs within MC38 tumors. To define if peripheral cancer-driven neuroinflammatory mechanisms can be exacerbated by ICIs treatment, a potential direct neurotoxicity of ICIs by treating cancer-naïve immunocompetent mice with anti-PD-1, anti-PD-L1, their combination (anti-PD-1+ anti-PD-L1) or IgG2a as control, once a week for three weeks before behavioral tests assessment (**Fig. 9a**). No alteration in spontaneous activity (OFT), anxiety evaluated both in EPM and LDB test, resignation (TST) and cognition (NORT) was detected in ICIs-treated mice compared with IgG-treated control mice suggesting that the appearance of ICI-induced neurotoxicity, as already shown for cardiac toxicity^24^, should be related to immune mechanisms already engaged before ICIs administration. Among anti-PD-1 treated mice, a significant aggravation of resignation-like behavior was found in MC38 mice compared with IgG-treated mice, while a tendency in short-term memory deficits was found in both B16F10 and MC38-mice compared with respective IgG controls. Then ICI were tested in cancer-bearing mice (**Fig. 9b**) and behavioral tests were followed between D4 and D8 post - tumor injections. Among anti-PD-L1 treated mice, a significant aggravation of anxiety-like behavior, as suggested by a decrease in time spent in open arms (EPM) and exacerbation of short-term memory impairment as indicated by a decreased preference index (NORT), was observed in both B16 and MC38-bearing mice (**Fig. 9c-d**).

**Figure 9.**
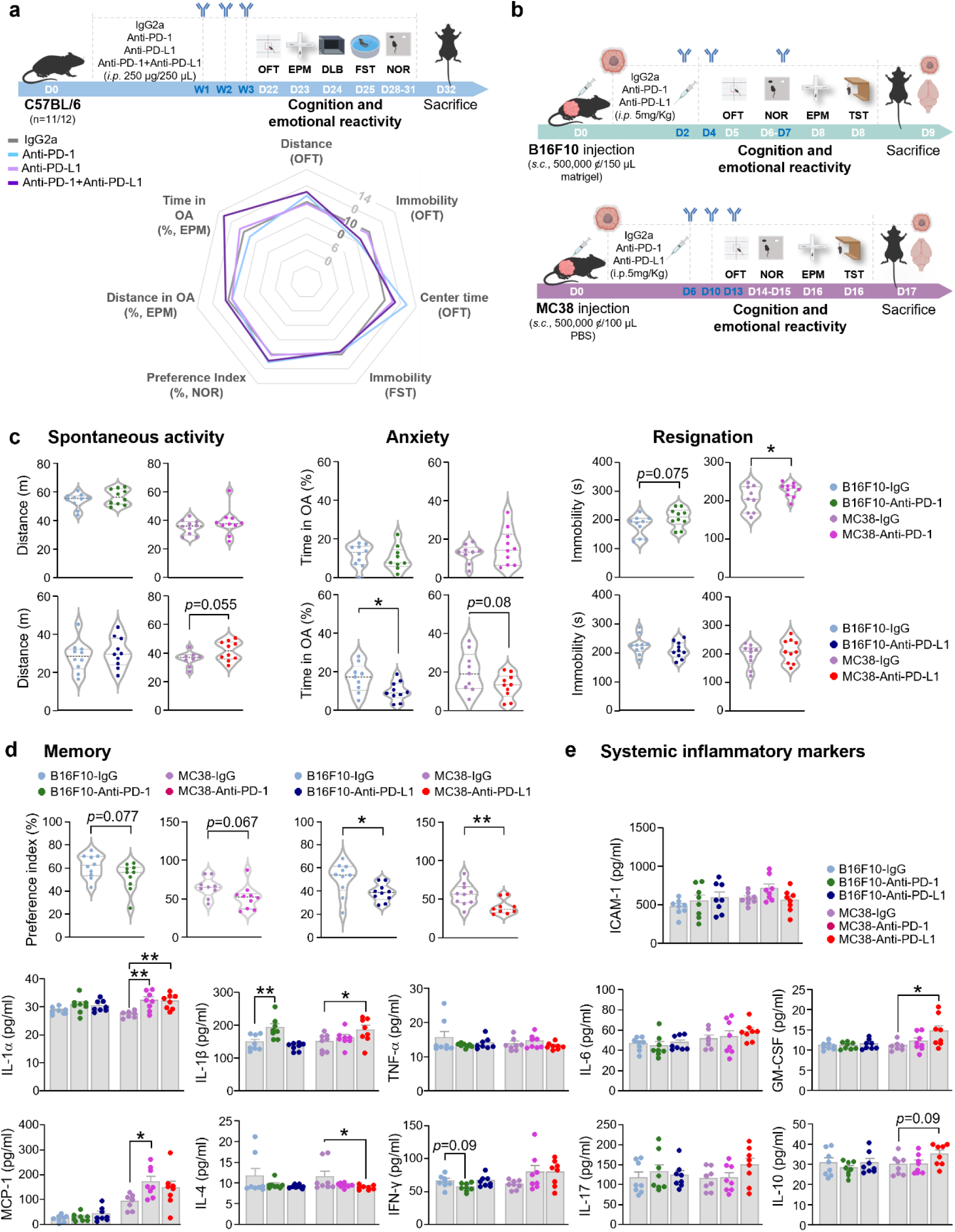
Impact of anti-PD1 and anti-PD-L1 on spontaneous activity, emotional reactivity and cognitive functions in c-ancer-bearing mice. **a.** Schematic timeline of anti-PD1, anti-PD-L1, anti-PD1 + anti-PD-L1 combination and IgG administration in cancer-naïve mice. Immunocompetent cancer-naïve mice were intraperitoneally injected (i.p.) with anti-PD-1, anti-PD-L1, anti-PD-1 + anti-PD-L1 and IgG once for week for three weeks (n=11/12). Starting from day 22, OFT, NORT, EPM and TST were used to evaluate activity, emotional and cognitive behaviors in mice. Animals were sacrificed the day after the end of the behavioral sessions. Below, comparative Radar Chart plot of performance index scores for emotional reactivity and cognitive functions after anti-PD-1 (blue), anti-PD-L1 (pink) and anti-PD-1 + anti-PD-L1 (purple) compared with IgG-treated mice (grey). On the right, bars showing the statistical comparisons of the distance crossed in the OFT and the time spent in open arms (%) in the EPM after treatment with anti-PD-1 (light blue), anti-PD-L1 (pink) and anti-PD-1 + anti-PD-L1 (purple) compared with IgG (grey). Statistical analysis was performed by using One way ANOVA or Kruskal-Wallis test with Bonferroni or Dunn’s correction for multiple comparisons. Data are represented as bars with symbols for individual data points and they are expressed by mean ± SEM, n=11-12. **b.** Schematic timeline of anti-PD-1, anti-PD-L1 and IgG administrations in B16F10- (green) and MC38- (purple) bearing mice. B16F10 mice were injected (i.p.) with anti-PD-1, anti-PD-L1 or IgG at D2, D4 and D7. MC38 mice were injected (i.p.) with anti-PD-1, anti-PD-L1 or IgG at D6, D10 and D13. Depending on the kinetic of tumor growth, behavioral tests were performed at D5-D8 (B16F10) or D13-D16 (MC38). Mice were sacrificed the day after the end of the behavioral session (B16F10: D9, MC38: D17) and brain, blood and tumors were collected for further analysis. **c.** Impact of anti-PD-1, anti-PD-L1 and IgG in B16F10- or MC38-bearing mice on spontaneous activity (left panel), on anxiety (middle panel) and resignation (right panel). Statistical analyses were performed (n=10) using unpaired t-test. Data are represented as violin plot with symbols for individual data points and they are expressed by mean ± SEM, * p<0.05. **d.** Impact of anti-PD-1, anti-PD-L1 and anti-IgG2a in B16-F10 or MC38-bearing mice on short-term memory. Statistical analyses were performed (n=10) using Mann-Whitney test; Data are represented as violin plot with symbols for individual data points and they are expressed by mean ± SEM, * p<0.05, ** p <0.01. **e.** Histograms quantification of 7 soluble inflammatory factors (IL-1α, IL-1β, MCP-1, ICAM-1, IL-6, IL-17, IL-10) levels analyzed in plasma of B16F10- and MC38-bearing mice treated with anti-PD-1 or anti-PD-L1 and compared with B16F10- and MC38-bearing mice treated with IgG. Statistical analyses (n=7-8) were performed by using one-way ANOVA or Kruskal-Wallis test with Bonferroni or Dunn’s correction for multiple comparisons (F10-IgG vs F10-PD1 or F10-PD-L1, MC38IgG vs MC38-PD-1 or MC38-PD-L1). Data are represented as bars with symbols for individual data points and they are expressed by mean ± SEM, * p<0.05, ** p <0.01. D: day, i.p.: intraperitoneal, DLB: dark light box, EPM: elevated plus maze, FST: forced swim test, ICAM-1: Intercellular Adhesion Molecule 1, IgG: immunoglobulin G, IL: interleukin, INF: interferon, MCP-1: monocyte chemoattractant protein 1, MHCII: major histocompatibility complex 2, NOR: novel object recognition test, OFT: open field, PD-1: programmed cell death 1, PD-L1: programmed cell death ligand 1, TNF: tumor necrosis factor.

To study whether the impact of anti-PD-1 or anti-PD-L1 administration on mouse behaviors was relayed by systemic inflammatory reactions, again, levels of the 11 inflammatory mediators (IL-17, IL-10, IL-4, INFγ, TNFα, IL-6, MCP-1, GM-CSF, IL-1α, IL-1β, ICAM-1) were measured in plasma of B16- and MC38-bearing mice, treated with either anti-PD-1, anti-PD-L1 or IgG2a (**Fig. 9e**).

We found an increase in IL-1α concentration in plasma of MC38-PD-1 and MC38-PD-L1 mice compared with MC38-IgG mice, an increase of MCP-1 in MC38-PD-1 mice compared with MC38-IgG mice, and an increase in IL-1β in B16F10-PD-1 and MC38-PD-L1 mice. All together, these results suggests that anti-PD1 and anti-PD-L1 neurotoxicity depends on cancer-associated immune mechanisms and that, despite their anti-tumoral activity, ICIs differentially aggravate cancer-induced behavioral alterations as well as cancer-induced systemic inflammation.

### ICIs aggravate systemic and vascular inflammation, brain vascular permeability and impair neurogenesis

As changes in peripheral cytokines are known to impact BBB integrity^25^ and vascular inflammation^26^, we tested how the presence of cancer would contribute to the effect of ICIs on BBB permeability and endothelial inflammation. The BBB was assessed *in vivo* by systemic injection of the fluorescent tracer Dextran_488_ in B16F10- and MC38-bearing mice treated with anti-PD-1, anti-PD-L1 or IgG2a (**Fig. 10a**). Surprisingly, an exacerbation of BBB permeability was detected for both cancer but exclusively when mice are treated with anti-PD-L1 compared with IgG. Moreover, treatment by anti-PD-L1 led to a marked augmentation of ICAM-1 immunofluorescence coverage of Lectin^+^ vessels within the DG of the hippocampus of B16 and MC38 mice, also observed in MC38 treated by anti-PD-1 compared with MC38-IgG mice (**Fig.10b**). To better understand if vascular permeability and inflammation could lead to neuroinflammation, we analyzed the levels of mRNAs expression of 48 gene of inflammation by qPCR in hippocampus of B16- or MC38-bearing mice treated with ICIs. Among the genes analyzed, CCL19 expression level was found stimulated in MC38-PD-1 and MC38-PD-L1 mice compared with MC38-IgG controls (**Fig. 10b**). We further observed by immunohistochemistry a marked increase in CCL19^+^ labeling intensity in Lectin^+^ vessels in the DG of MC38-PD-1 and MC38-PD-L1 compared with MC38-IgG mice (**Fig. 10b**). This feature was associated with a significant enhancement of extravascular CD3^+^ lymphocytes in the DG of MC38-PD1 and MC38-PD-L1 compared to MC38-IgG mice (**Fig.10c**). In the hippocampus, by counting the number of BrdU^+^ NPCs, we found a significant attenuation in MC38-PD-1 and MC38-PD-L1 compared with MC38-IgG mice suggesting that ICIs can worsen neurogenesis only in immune-inflamed cancer-bearing mice (**Fig 10d**). We then verified whether vascular inflammation, vascular permeability and T lymphocytes infiltration due to ICIs also exacerbate neuroinflammation and microglia reactivity in the DG of B16- and MC38-bearing mice.

**Figure 10.**
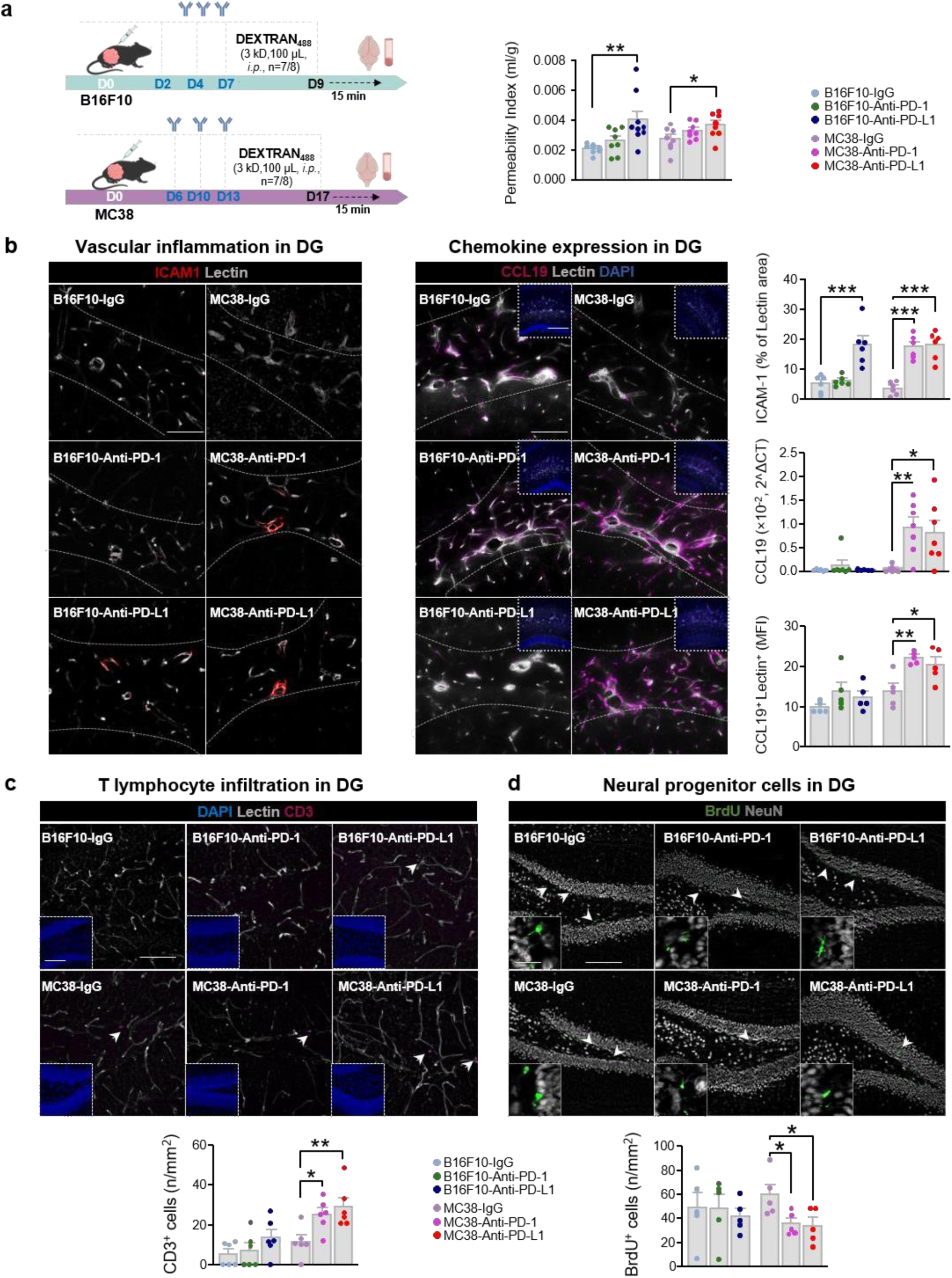
Anti-PD-1 and anti-PD-L1 impact on cerebral vascular permeability, inflammation and neurogenesis. **a**. Schematic diagram showing the timeline used for the *in vivo* blood-brain barrier permeability assay in B16F10- (blue) and MC38- (pink) bearing mice treated with anti-PD-1, anti-PD-L1 and IgG. After tumors inoculation (D0), mice received 100 µl of Dextran_488_ (3kD, i.p) at D9 for B16-bearing mice and at D17 for MC38-bearing mice. After 15 min post-injection, mice were sacrificed, while brains and plasma were collected for fluorescence quantification. On the left, bars represent the permeability index of B16- and MC38-mice treated with IgG compared with B16- and MC38-mice treated with anti-PD1 and anti-PD-L1. Statistical analyses were performed by using one-way ANOVA with Bonferroni correction for multiple comparisons (F10-IgG vs F10-PD1 or F10-PD-L1, MC38IgG vs MC38-PD-1 or MC38-PD-L1). Data are represented as bars with symbols for individual data points and they are expressed by mean ± SEM, n=7-8, * p<0.05, ** p <0.01. **b**. Representative micrographs of ICAM-1 (red) immunolabeling in Lectin^+^ (grey) vessels in the DG of the hippocampus of B16- and MC38-bearing mice treated with IgG, anti-PD1 and anti-PD-L1. Grey dotted lines delineate the granular layer of DG. Scale bar: 50 μm. On the right panel, representative immunoreactivity of CCL19 (magenta) expression in Lectin^+^ (grey) vessels of the DG of B16- and MC38-bearing mice treated with IgG, anti-PD-1 and anti-PD-L1. Boxed area illustrates the structure of the dentate gyrus by means of cell nuclei DAPI staining. Scale bars: 50 μm, boxed areas: 100 μm. On the right, histograms representing the statistical comparison of ICAM1 staining coverage (n=6), expressed as the percentage of the Lectin+ area, in vessels of DG of B16- and MC38-bearing mice treated with anti-PD-1 and anti-PD-L1 compared with IgG. Below, bars represent quantification of CCL19 mRNA levels (n=7) in hippocampal tissues isolated from brains of B16- and MC38-bearing mice treated with anti-PD-1 and anti-PD-L1 compared with IgG. Below, histograms representing the intensity of the immunofluorescence detection (n=5) of CCL19 on Lectin^+^ vessels in the DG of B16- and MC38-bearing mice treated with anti-PD-1 and anti-PD-L1 compared with IgG. Statistical analyses were performed by using one-way ANOVA or Kruskal-Wallis test with Bonferroni or Dunn’s correction for multiple comparisons (B16F10-IgG vs B16F10-PD1 or B16F10-PD-L1, MC38IgG vs MC38-PD-1 or MC38-PD-L1). Data are represented as bars with symbols for individual data points and they are expressed by mean ± SEM, * p<0.05, ** p <0.01, *** p <0.001. **c.** Representative microphotographs of T lymphocytes (CD3^+^, magenta) in proximity of Lectin^+^ vessels (grey) in the DG of B16F10- and MC38-bearing mice treated with anti-PD1, anti-PD-L1 and IgG. White arrows highlight CD3^+^ T lymphocytes in hippocampal parenchyma. Boxed area represents the granular layer of DG (DAPI, blue). Scale bars: 50 μm, boxed areas 100 μm. **d**. Representative microphotographs of proliferating NPCs (BrdU^+^, green) and mature neurons (NeuN^+^, grey) in the DG of B16F10-PD-1, B16F10-PD-L1, MC38-PD-1, MC38-PD-L1 and their respective IgG control. The white squares show magnification of the BrdU^+^ and NeuN^+^ fluorescent cells. Scale bars: 50 μm, zoom 10 μm. **c, d. *Bottom***, Histograms representing statistical comparisons of the density of CD3^+^ T lymphocytes in the dentate gyrus and of NPC BrdU^+^ (n/mm^2^) of B16F10 or MC38-bearing mice treated with anti-PD-1 or anti-PD-L1 compared to IgG-treated controls. Statistical analyses were performed by one-way ANOVA or Kruskal-Wallis test with Bonferroni or Dunn’s correction for multiple comparisons (B16F10-IgG vs B16F10-PD-1 or B16F10-PD-L1 and MC38-IgG vs MC38-PD-1 or MC38-PD-L1). Data are represented as bars with symbols for individual data points and they are expressed by mean ± SEM, n=5-6, * p<0.05, ** p <0.01. BrdU: Bromodeoxyuridine, CCL19: Chemokine (C-C motif) ligand 19, CD: cluster of differentiation, D: day, DAPI: 4’,6-diamidino-2-phenylindol, DG: dentate gyrus, IgG: immunoglobulin G, NeuN: neuronal nuclei antigen, PD-1: programmed cell death 1, PD-L1: programmed cell death ligand 1.

No significant alteration in the number of Iba1^+^CD68^+^ reactive microglia could be detected (**Fig. S6d**), suggesting that ICIs are not associated with a global exacerbation of hippocampal neuroinflammation in addition to the effect of the cancer itself.

Also, the density the immunolabeling of immunosuppressive CD206^+^ and inflammatory MHCII^+^ myeloid cells did not change in the VI area among B16 F10 and MC38 mice after treatment with anti-PD-1 or anti-PD-L1, suggesting that changes in immunophenotype of meningeal myeloid cells does not contribute to neurogenesis impairment or neuroinflammation in our mice (**Fig. S7c**). In order to verify a potential direct neurotoxic impact of anti-PD-1 or anti-PD-L1 by parenchymal penetration and resident brain cells targeting, the expression of PD-1 and PD-L1, an antibody directed against the IgG2a was specifically used to verify potential parenchymal extravasation of the peripheral administered anti-PD-1, anti-PD-L1 and IgG2a in brain slices (**Fig. S8**). IgG2a^+^ staining was exclusively observed in CP of B16-PD-L1 mice, but remained absent from the DG and frontal cortex of B16- and MC38-bearing mice receiving anti-PD1, anti-PD-L1 and IgG2a (**Fig. S8**), suggesting that ICIs associated neuroinflammatory alterations in cancer-bearing mice are not driven by ICIs crossing of the BBB and direct targeting of neural cells.

### Anti-PD-L1-induced TCRγδ infiltrates the blood-CSF barriers and relays cognitive deficits and anxiety-like behaviors in cancer-bearing mice

As peripheral immune cells, beyond cytokines, have been demonstrated to play key role in cerebral dysfunctions associated to cancer^21^, we then decided to characterize systemic blood immune cells composition in the presence of B16F10- and MC38-peripheral cancer and after treatment with ICIs. Through flow cytometry analysis, nine myeloid and nine lymphocytic subpopulations were identified from whole blood-derived immune cells sampled from B16F10 and M38 mice treated with IgG2a, anti-PD-1 or anti-PD-L1 and from PBS cancer-naïve mice (**Fig. 11a, Fig. S9a-b**). Increased number of leukocytes (CD45^+^), monocytes (CD11b^+^), M2-like macrophages (CD206^+^), TLs (CD3^+^), cytotoxic (CD8^+^) TLs and B lymphocytes (CD19^+^) was exclusively associated with MC38-IgG mice compared with PBS-mice, while, a specific increase in number of circulating natural killer cells (NK1.1^+^) was associated with B16-IgG mice compared with PBS-mice. Furthermore, the circulating γδ TLs number was exclusively found elevated in blood of B16-PD-L1 and MC38-PD-L1 mice compared with B16-IgG and MC38-IgG, respectively (**Fig. 11a**).

**Figure 11.**
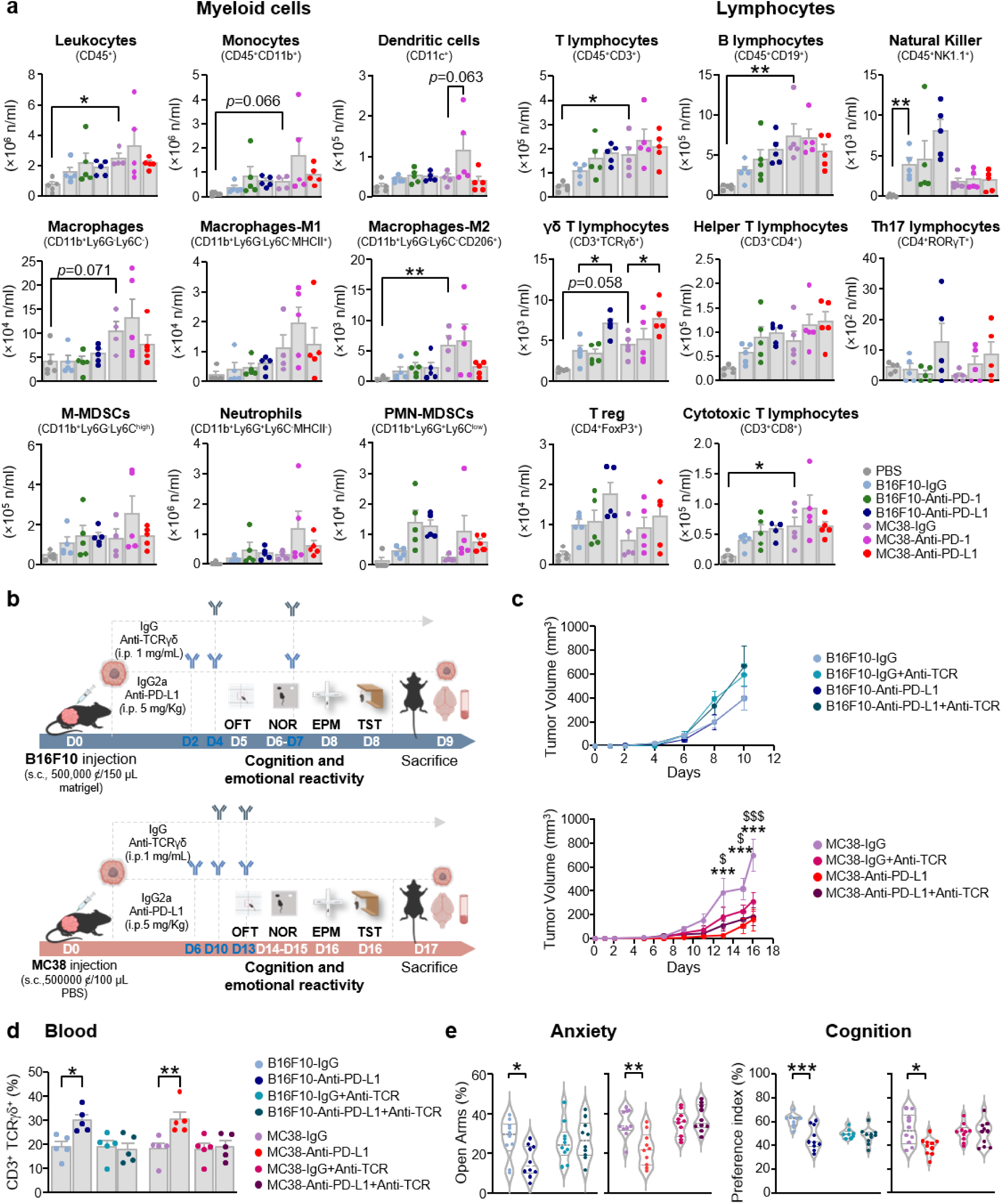
Specific impact of anti-PD-L1 on circulating and brain infiltration of γδ T lymphocyte relaying short-term memory and anxiety-like behaviors. **a.** Histograms representing statistical comparison of the number (n/ml) of myeloid (left) and lymphocyte (right) immune cells populations from whole blood collected from B16 or MC38-bearing mice treated with anti-PD1, anti-PD-L1or IgG. Data are represented as bars with symbols for individual data points and they are expressed by mean ± SEM, (n=4-5), * p<0.05, ** p <0.01. Statistical analyses were performed by one-way ANOVA or Kruskal-Wallis test with Bonferroni or Dunn’s correction for multiple comparisons (B16F10-IgG vs B16F10-PD-1 or B16F10-PD-L1 and MC38-IgG vs MC38-PD-1 or MC38-PD-L1 and PBS vs B16F10-IgG or MC38-IgG). **b.** Schematic timeline of immuno-neutralization of γδ T lymphocytes in B16- and MC38-bearing mice treated with anti-PD-L1 and IgG. B16F10 mice were injected (i.p.) with anti-PD-L1 or IgG on day 2-4-7 and with an anti-TCRγδ on days 4-7. MC38 mice were injected (i.p.) with anti-PD-L1 or IgG on days 6-10-13 and with anti-TCRγδ on days 6-10-13. Depending on the kinetics of tumor growth (B16: D5-D8, MC38: D13-D16), mice behavior was evaluated by means of OFT, NOR, EPM and TST tests, then sacrificed the day after the end of the behavioral session (B16: D9, MC38: D17) and brains, blood and tumors were collected for further analysis. **c**. Curves of tumoral volumes represented as mean ± SEM (n = 10) of B16-IgG/B16-PD-L1 and MC38-IgG/MC38-PD-L1 mice receiving or not anti-TCRγδ at indicated time points. Statistical comparison was performed by two-way ANOVA with Bonferroni test for multiple comparisons (MC38 IgG vs. MC38 IgG-TCRγδ, MC38 IgG vs. MC38 PD-L1, MC38 IgG vs. MC38 PD-L1-TCRγδ, MC38 IgG-TCRγδ vs. MC38 PD-L1, MC38 IgG-TCRγδ vs. MC38 PD-L1-TCRγδ, MC38 PD-L1 vs. MC38 PD-L1-TCRγδ, *** p<0.001 (IgG vs PD-L1), $ p<0.05, $$$ p<0.001 (IgG vs IgG-TCRγδ). **d.** Impact of anti-TCRγδ treatment on the proportion of circulating CD3^+^TCRγδ^+^ lymphocytes in blood of B16- and MC38-bearing mice treated with anti-PD-L1 and IgG. Statistical comparison was performed by one-way ANOVA test with Bonferroni correction for multiple comparisons (B16F10 IgG vs. B16F10 PD-L1, B16F10 IgG-TCRγδ vs. B16F10 PD-L1-TCRγδ, B16F10 PD-L1 vs B16F10 PD-L1-TCRγδ, B16F10 IgG vs B16F10 IgG-TCRγδ, MC38 IgG vs. MC38 PD-L1, MC38 IgG-TCRγδ vs. MC38 PDL1-TCRγδ, MC38 PD-L1 vs. MC38 PD-L1-TCRγδ, MC38 IgG vs. MC38 IgG-TCRγδ). Data are represented as bars with symbols for individual data points and they are expressed by mean ± SEM, * p<0.05, ** p<0.01. **e.** Impact of anti-TCRγδ treatment on anxiety-like behaviors in EPM and on short-term memory in NORT in B16- and MC38-bearing mice treated with anti-PD-L1 and IgG. Statistical analyses were performed (n=10) using one-way ANOVA or Kruskal-Wallis with Bonferroni or Dunn’s correction for multiple comparisons (B16F10 IgG vs. B16F10 PD-L1, B16F10 IgG-TCRγδ vs. B16F10 PD-L1-TCRγδ, B16F10 PD-L1 vs B16F10 PD-L1-TCRγδ, B16F10 IgG vs B16F10 IgG-TCRγδ, MC38 IgG vs. MC38 PD-L1, MC38 IgG-TCRγδ vs. MC38 PDL1-TCRγδ, MC38 PD-L1 vs. MC38 PD-L1-TCRγδ, MC38 IgG vs. MC38 IgG-TCRγδ). Data are represented as violin plot with symbols for individual data points and they are expressed by mean ± SEM, * p<0.05, ** p <0.01. CD: cluster of differentiation, CD11b: integrin alpha M subunit, CD11c: complement component 3 receptor 4 subunit, CD206: cluster of differentiation 206, CD45: leukocyte common antigen, D: day, DAPI: 4’,6-diamidino-2-phenylindol, EPM: elevated plus maze, FoxP3: forkhead box P3, IgG: immunoglobulin G, i.p.: intraperitoneal injection, Ly6C: lymphocyte antigen 6 family member C, Ly6G: lymphocyte antigen 6 family member G, MHCII: major histocompatibility complex 2, M-MDSC: monocytic-derived myeloid derived suppressor cells, NeuN: neuronal nuclei antigen, NK: natural killer, NOR: novel object recognition test, OFT: open field test, PD-1: programmed cell death 1, PD-L1: programmed cell death ligand 1, PMN-MDSC: polymorphonuclear myeloid derived suppressor cells, ROR: Retinoic acid-related Orphan Receptors, SSC: side scattering gating, TCR: T cell receptor, TST: tail suspension test.

As meningeal γδ T lymphocytes have been recently demonstrated to be involved in the control of anxiety-like behaviors^27^ and short-term memory^28^, we wanted to study whether increased number of circulating γδ TLs could be the origin of anti-PD-L1 aggravation of cognitive functions and anxiety-like behaviors observed in cancer mice. We specifically aimed to neutralize peripheral γδ TLs by means of two consecutive intraperitoneal injections of an anti-TCRγδ antibody before behavioral assessment (**Fig. 11b**).

**Figure 12.**
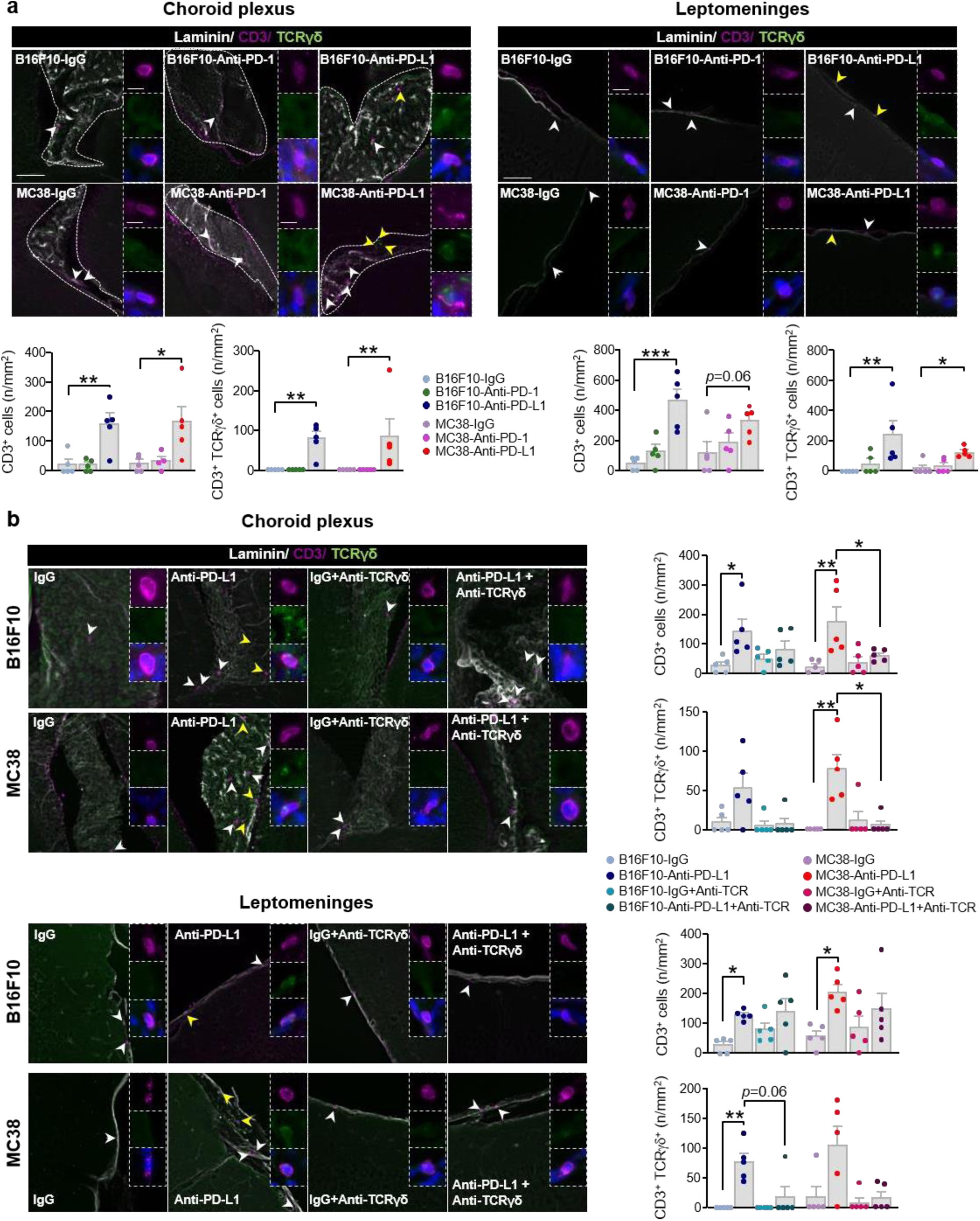
Impact of anti-PD-1 and anti-PD-L1 on the infiltration of γδT lymphocytes at the blood-CSF barrier. **a.** Representative images of immunohistochemical detection of CD3 (magenta), TCRγδ (green) and Lectin (grey) immunoreactivity in CP (left panel) and leptomeninges (right panel) of B16F10 or MC38-bearing mice treated with anti-PD-1, anti-PD-L1 or IgG. Boxed areas represent magnification of CD3 (magenta) T lymphocyte and colocalization with TCR γδ (green) and DAPI (blue) immunolabeling. White and yellow arrows indicate CD3^+^ and CD3^+^ TCRγδ^+^ staining respectively. Below, histograms represent quantification of the number of CD3^+^ T lymphocytes and of CD3^+^TCRγδ^+^ lymphocytes (density, n/mm^2^) in CP (left) and leptomeninges (right) of B16 or MC38-bearing mice treated with anti-PD1 or anti-PD-L1 compared to IgG-treated controls. Statistical analysis was performed by using One way ANOVA or Kruskal-Wallis test with Bonferroni or Dunn’s correction for multiple comparisons (F10-IgG vs F10-PD-1 or F10-PD-L1, MC38IgG vs MC38-PD1 or MC38-PD-L1). Data are represented as bars with symbols for individual data points and expressed by mean ± SEM, n=5, * p<0.05, ** p <0.01. Scale bars: 50 μm, zoom 10 μm. **b.** Representative microphotographs of CD3^+^ (magenta), TCRγδ^+^ (green) and Lectin^+^ (grey) immunoreactivities in CP (upper panel) and leptomeninges (lower panel) of B16 or MC38-bearing mice treated with anti-PD-L1 or IgG control and receiving or not anti-TCRγδ treatment. Boxed areas illustrate magnification of the CD3^+^ (magenta) T lymphocyte fluorescent staining and colocalization with TCRγδ (green) immunolabeling with nuclei stained by DAPI (blue). On the right, histograms quantification of the density (n/mm^2^) of CD3^+^ and CD3^+^TCRγδ^+^ T lymphocytes in CP (up) and leptomeninges (below) of B16F10- and MC38-bearing mice treated with anti-PD-L1 or IgG and receiving of not anti-TCRγδ treatment. Statistical analyses were performed (n=5) using one-way ANOVA or Kruskal-Wallis test with Bonferroni or Dunn’s correction for multiple comparisons (B16F10 IgG vs. B16F10 PD-L1, B16F10 IgG-TCRγδ vs. B16F10 PD-L1-TCRγδ, B16F10 IgG vs. B16F10 IgG-TCRγδ, B16F10 PD-L1 vs. B16F10 PD-L1-TCRγδ, MC38 IgG vs. MC38 PD-L1, MC38 IgG-TCRγδ vs. MC38 PD-L1-TCRγδ, MC38 IgG vs. MC38 IgG-TCRγδ, MC38 IgG vs. MC38 IgG-TCRγδ, MC38 PD-L1 vs. MC38 PD-L1-TCRγδ). Data are represented as bars with symbols for individual data points and they are expressed by mean ± SEM, * p<0.05, ** p <0.01, *** p <0.001. CD: cluster of differentiation, CP: choroid plexus, DAPI: 4’,6-diamidino-2-phenylindol, IgG: immunoglobulin G, PD-1: programmed cell death 1, PD-L1: programmed cell death ligand 1, TCR: T cell receptor.

Concerning tumoral growth, anti-TCRγδ injections did neither affect the therapeutic efficacy of anti-PD-L1 nor the proportion of other peripheral immune cells in B16 and in MC38-mice (**Fig. 11c, Fig. S10-12**). However, a significant reduction in proportion of circulating γδ TLs was observed in B16-PD-L1 and MC38-PD-L1 mice receiving anti-TCRγδ compared with B16-PD-L1 and MC38-PD-L1 receiving IgG-control, confirming optimal depletion of circulating γδ TLs (**Fig. 11d).** Interestingly, the anti-TCRγδ prevented the short-term memory impairments in the NORT and the anxiety-like behavior in the EPM of B16-PD-L1 and MC38-PD-L1 mice compared with B16-IgG and MC38-IgG mice respectively, suggesting that γδ TLs are involved in anti-PD-L1 induced cognitive deficits and anxiety-like behaviors (**Fig. 11e**). Beyond blood, increased number of CD3^+^ TLs and among them, of γδ TLs, were detected in leptomeninges and CP, but not in the hippocampus, of B16F10-PD-L1 vs B16-IgG mice and MC38-PD-L1 vs MC38-IgG mice (**Fig.12a, Fig S13**). Interestingly, after treatment with anti-TCRγδ, the increase of γδ LTs associated to anti-PD-L1 treatment was totally prevented in CP and leptomeninges of MC38-PD-L1 and B16-PD-L1 mice, suggesting that systemic γδ LTs are likely to impact cognitive and emotional behaviors by infiltrating CSF-filled spaces via CP (**Fig.12b**).

## Discussion

Up to now, an inflammatory challenge of the cancer itself by inducing systemic inflammation is assumed to be involved in behavioral alterations and cognitive impairment in CRCI while even though neuroinflammation seems to be an undeniable consequence, underlying mechanisms remain poorly understood^88^. Here it was sought to generate a comparative assessment of the effects of cancer according to its immunogenic and inflammatory status, and the functional interaction between these different cancer types and ICI-induced potential neurotoxicity. Among ICIs, anti-PD-1 antibodies nivolumab and pembrolizumab have demonstrated important efficacy compared with standard therapy in metastatic melanoma, advanced lung cancer, metastatic renal cancer and bladder cancer^89–92^, leading to prolonged survival and even durable remission^93^. But the tremendous hope raised by these therapies is currently flowed by various inflammatory toxicities refereed as irAEs that can be exacerbated when ICIs are used in association with radiotherapy or chemotherapy^94^. These treatments can more specifically be associated with neurological toxicities (e.g. encephalopathy) and autoimmune nervous side effects such as hypophysitis, the most frequent reaction in patients treated with anti-CTLA-4, and migraines or fatigue affecting cognitive functions^95,96^. Thus, the impact of ICIs likely modifying the immune balance, on neurobiological mechanisms remains to be established and there is only a preclinical model to address this^60^. A recent study showed that anti-CTLA-4 immunotherapy combined with peripheral targeted radiotherapy led to anxiety behavior and cognitive impairment associated with microglia activation in mice^65^ but the mechanisms linking tumor and immunotherapy are not understood.

Here we wanted to develop a mirror preclinical behavioral study of our Cog-Immuno multicenter longitudinal trial^56^ investigating the incidence and the severity of cognitive impairment and its impact on QoL in cancer patients treated by immunotherapy. Our objective was to evidence the impact of anti-PD-1 and anti-PD-L1 on cognitive functions and emotional reactivity and to characterize a signature of biomarkers predictive of the occurrence of neurological toxicities, depending on cancer immune and inflammatory profiles. Here, we have established that murine cancers differently alter activity, emotional reactivity and cognitive functions of mice depending on their corresponding systemic inflammation and immune responses, thus leading to common neuroinflammatory mechanisms and distinct leucocytic infiltration status. In addition, we have shown that anti-PD1 and anti-PD-L1 are associated with vascular inflammation and T lymphocytes infiltration in the hippocampal parenchyma exclusively in the presence of the most the inflamed cancer MC38, and that anti-PD-L1 whatever the cancer promotes γδT lymphocytes in the blood and in CP and leptomeninges, relaying cognitive deficits and anxiety-like behavior.

In our study, B16F10, B16F10-Ova and MC38 cancer cell lines xenografted in immunocompetent C57B/l6 mice were first selected for their potentially poorly immunogenic, intermediate, and highly immunogenic status. Immunogenicity is widely defined as the ability of a tumor to induce an immune response^97^ which depends on high tumoral mutational burden (TMB) supplier of tumor neoantigens^77^ and their capacity of presentation by the MHC class I pathway expressed by tumor cells for specific recognition by CD8^+^TL^98,99^ and subsequent intratumoral TL infiltration (TILs)^100^. The BF16 melanoma tumor model, in spite of its moderate TMB level^101^, is frequently classified as poorly immunogenic because of the loss of MHC-I expression and/or functional pathway and consequently poor TILs infiltration^75,78,102^, building a non-inflamed TME enriched with Treg and MDSCs^58,103^. In agreement, we provide evidence of high density of CD11b^+^Gr1^+^ MDSCs, pro-tumoral CD206^+^ TAMs and corresponding relative low levels of CD8^+^ TILs the tumoral bulk of B16F10 xenografted tumors. In contrast, the murine colon carcinoma MC38 was previously recognized as a high TMB exhibiting high surface expression of MHC-I as well as with TILs within the TME^78,104^. In highly immunogenic tumors comprising these TILs, co-occurrence of inflamed TME has been described comprising a wide variety of immune B lymphocytes and anti-tumoral TAMs^79^. Beside highest density of CD8^+^ TILs, we show a marked expression of MHC-II^+^ pro-tumoral TAMs, CD11b^+^ monocytes and Meca79^+^ tumor-associated high endothelial venules (TA-HEV), typical blood vessels specialized in recruiting lymphocytes to lymphoid organs^80–82^ in the MC38 colon cancer model, thus pinpointing its immunogenic and inflammatory status. To these two types of cancers with extreme phenotype, we must add “Intermediate” immunogenic tumors, characterized by intact MHC-I system presentation, combined with an immunosuppressive TME^57^. We established the engineered non-immunogenic B16F10 cell line expressing the chicken ovalbumin antigen (B16F10-Ova)^105^ resembles B16F10 tumors by displaying higher density of immunosuppressive CD11b^+^Gr1^+^ MDSCs and pro-tumoral CD206^+^ TAMs as well as CD8^+^TIL exclusion, but also MC38 by highest density of CD11b^+^ monocytes and anti-tumoral MHC-II^+^ TAMs. This suggests that B16F10-Ova tumors could be representative of an intermediate immunogenic profile characterized by an immunosuppressive and immuno-excluded microenvironment. One of the contributors of this immunosuppressive TME is the expression of PD-L1, a major immune-escape cloaking signal expressed in tumor cells and hiding a tumor from immune surveillance by inactivating TILs expressing PD-1 and/or promoting Treg development^106–108^. In tumor bulk immunohistochemical analysis, PD-L1 expression appears strongly lower in MC38 tumors than in B16F10 or B16F10-Ova. Although in a clinical context PD-L1 can both confer an unfavorable prognosis in a variety of solid tumors^109–112^ being inversely correlated with TILs in NSCLC and glioma^113,114^ and be associated with TILs being a marker of good prognosis cancers^115–118^, in BF1016 and MC38, the presence of immune-infiltrating cells appear of critical importance in suppressing the anti-tumoral immunity^119,120^. Considering that PD-L1 can be expressed by CD11b^+^Gr1^+^ MDSCs and anti-tumoral TAMs in both B16F10- and MC38-bearing mice models, the decreases density of PD-L1^+^ in MC38 might relate to the reduced level of MDCSs^121–123^. From a clinical scoring system called “Immunoscore®” based on a tumor-containing TILs status, to discriminate responders from non-responders among all cancer patients^124,125^, we also build an “immunoscore” and an “immunosuppressive score”, introducing additional items to make this score more versatile and representative. For this purpose, we here integrated immunolabeling of CD8^+^TILs, CD11b^+^Gr1^-^ monocytes, MHCII^+^ TAMs and Meca-79^+^ HEVs as anti-tumoral marker, and of CD206^+^ TAMs, CD11b^+^Gr1^+^ MDSCs and PD-L1 as pro-tumoral markers, leading to the accepted “immune desert” term for BF16 which were characterized by the highest immunosuppressive score, “immune-excluded” for B16F10-Ova characterized by intermediate values of both immunosuppressive and immunoscore and “immune-inflamed” for MC38 we classify with the highest immunoscore and the lowest immunosuppressive score. Then, we assessed whether the different immune cells populating immune-desert, immune-excluded and immune-inflamed cancers drive changes 10 cytokines tested. Interestingly, only IL-1β appears stimulated in the plasma after more than ten days of B16F10 tumoral growth. Within TME, IL-1β is mainly expressed by innate cells, both by pro-tumoral TAMs and MDSCs but also in a subset of TAMs expressing inflammatory but non-cytotoxic programs (and correlated with circulating IL-1β), whose abundance correlates with poor prognosis in PDAC^126,127^. IL-1β would here result from the high density of CD206^+^ TAMs and MDCS in BF16, but also the high density of MHC-II anti-tumoral TAMs in MC38, thus suggesting that this cytokine should not be considered as signature of immune desert cancer. MC38 differs from B16F10 since IL-6, IL-1α, IL-17 and IL-10 were found elevated in the plasma of mice bearing cancer. In particular, anti-tumoral TAMs are known as the main source of IL-6 in melanoma or carcinoma TME^128,129^ and level of plasma IL-6 is associated with the density of anti-tumoral TAMs in these tumors. In addition, the MCP-1 increased circulating level has been previously reported in the context of inflamed cancer^130,131^. Interestingly, the secretion of MCP-1 (CCL2) by MC38 tumoral cells was demonstrated to drive the recruitment of monocytic and pro-tumoral TAMs *via* the CCL2-CCR2 axis in the TME and is the unique chemokine also found in the serum after 1 week of post-MC38 inoculation^119^. We thus hypothesize that high density of CD11b^+^ monocytes and MHC-II^+^ pro-tumoral TAMs within the MC38 TME, involve MCP-1 secretion by tumor cells MCP-1 being a key factor at play. In addition to MCP-1, IL-10 and Il-17 when secreted in the TME have been shown to mainly result from TME-hosting Treg^132^ and MDSCs and a decreased cytotoxic activity of TILs^133^, suggesting the presence of Treg in our MC38 model. Interestingly, the pro-tumoral role IL-17 has been previously demonstrated by using IL-17^-/-^ mice, or by injecting adenovirus deleting locally IL-17A expression^133^, since tumor-infiltrating CD8^+^ T cells were increased and produced more IFN-γ compared with WT mice^134,135^. In addition, IL-17-exposed monocytes were found to suppress cytotoxic T-cell immunity *in vitro*^136^. Together, we may propose that IL-10, MCP-1 and IL-17 in plasma should reflect the immunosuppressive environment created by TILs and Treg, while IL-6 at least in part might constitute the unique cytokine signature of anti-tumoral activity in MC38.

As inflammatory cytokines and chemokines specifically expressed in the CNS have already been shown to impact different aspects of neuroinflammation and cognitive functions such as hippocampal long-term potentiation and memory and white matter damages^137^. In cancer patients, increased in peripheral cytokines including IL-6 were reported and mostly associated with cognitive complaints, and objective cognitive impairment prior to treatment^39,138^. We here originally establish a correlation between the different cancer-induced immune and inflammatory challenges and the level of exploratory/activity, emotional and cognitive impact in mice. Short-term memory/executive functions are systematically altered whatever the presence of peripheral immuno-inflamed or immuno-desert cancer, but a decrease in spontaneous activity/exploration, and altered emotional reactivity including anxiety-like behavior and resignation (depressive-like symptom) are exclusively detected in the immune-excluded B16F10-Ova and immune-inflamed MC38-bearing mice. Even if the contribution of tumoral burden was previously associated with mouse depressive-behavior^139^, we note the absence of correlation between each behavioral item cognitive or emotional ones and the tumor volume, excluding physical interference like reinforcing systemic immunity and/or nerve contribution on behavioral deficits. To discriminate the effects of immune challenge associated with cancer immunogenicity, and the inflammatory situation generated by the presence of cancer/TME, B16F10, B16-Ova and MC38 tumoral necrotic lysates were injected into C57B/l6 cancer-naïve mice to optimize the exposure of tumoral associated antigens (TAA)^140^ and to increase the secretion of heat shock proteins (HSPs), such as HSP60, HSP70, and HSP90 by dying cancer cells^141,142^ without the multicellular TME. We find that B16F10 lysate is not inductor of any alteration in cognition and emotional reactivity as well as cytokine plasma level modification, in agreement with experimental models using B16F10-derived cell lysates as vaccines, succeeding in a therapeutic response only when melanoma cells were genetically modified to express the antigen Ova for instance^143–146^. This confirms the weak immunogenicity of B16F10 cells potentially due to suboptimal activation of DCs and/or an inefficient delivery of relevant TAA through deficient MHC-I pathway to resident DC populations responsible for cross-priming of CD8^+^ T cells^142,147,148^. Accordingly B16F10-Ova lysate, a better model of tumoral antigen-induced immunogenicity^77^ when injected in mice, led to cognitive performance deficit and resignation-like behavior, associated with plasma IL-6 elevated level. Moreover, besides cognitive deficits and resignation alterations, and additional increase in anxiety-like behavior was measured in mice injected with MC38 lysate in association with increased IL-1β plasma level. As IL-6 and IL-1β are the main cytokines of DC maturation^142,149^, these cytokines may result from B16F10-Ova and MC38 antigen-presentation processes.

To connect the potential of circulating cytokines in driving behavioral alterations, we looked forward correlations between plasma cytokines levels and behavioral results in cancer-bearing mice. A positive correlation between increase in plasma levels of IL-6 and immobility time in the TST suggesting a link between IL-6 and depressive-like behavior. Systemic IL-6 level has already been described in association with a resigned profile and with IL-6 expression in the hippocampus^150^ in cancer mice, while IL-6 neutralization led to improve depressive-like symptoms in murine model of depression^151^, all consistent with the increased level of IL-6 in plasma of cancer patients with depressive symptoms in melanoma, peritoneal carcinoma and breast cancer patients before treatment^88,152,153^. Interestingly, we also show a positive correlation between plasma IL-17 and the depressive like behavior, suggesting as the increased level of blood IL-17 would accompagny brain Th-17 lymphocyte infiltration to drive resignation behavior in mice^154^. In addition to IL-6 and IL-17, the correlated systemic MCP-1 plasma concentration and the resigned behavior of MC38 mice also suggests controlled depressive-like behavior by MCP-1 (CCL2) acting on CCR2-expressing dopaminergic neurons of *nucleus accumbens*, and thus leading to dopaminergic transmission defaults^155^, or on serotoninergic neurons of raphe nucleus in mice^156^. In addition to these cancer-evoked depressive symptoms, we importantly stress a correlation between increased circulating levels of IL-6 and decreased shot-term memory performance. The causal role of circulating cytokines in general, and IL-6 in particular in modulating cognitive functioning has been studied more specifically in neurodegenerative disorders in which serum IL-6 concentration has been related to an increased risk of dementia and cognitive decline over time in humans^157^, and aging-related cognitive decline and microglial cell reactivity^158^ or Alzheimer (AD)-related cognitive decline by tau phosphorylation in hippocampal neurons in mice^159^. The direct impact of IL-6 was better demonstrated by using the anti-IL-6 receptor antibody tocilizumab in an experimental murine model of AD or of autoimmune arthritis showing improvement in learning and spatial memory functions^160,161^, thus supporting a key role of IL-6 in inflamed cancer-mediated cognitive dysfunctions in our models and in cancer patients.

Peripheral inflammatory stimuli may obviously damage CNS by altering brain barriers, such as the BCSFB, constituted by CP epithelial cell layer and fenestrated vessels, which serves at steady state as active and selective immune-skewing gate. In a context of peripheral inflammation, IL-1β or IL-6 have been featured in BCSFB permeability and upregulation of adhesion molecules such as L-selectin, ICAM-1, and VCAM-1 on CP epithelial cells^162–169^, thus promoting access of immune cells into the brain. In our study, in B16F10-related plasma IL-1β and MC38-associated plasma IL-6, the expression of adhesion molecules ICAM-1 and MadCam-1 was markedly increase in CP compartment in Lectin+ cells indicative of an immuno-desert or immuno-inflamed cancer-driving CP neuroinflammation. This mechanism could lead to a neuroinflammatory loop since CP stroma can consequently synthesize and release cytokines such as IL-1β and chemokines such as MCP-1, crucial for the recruitment of immune cells, including macrophages and T lymphocytes during systemic inflammation^170–173^. This is supported by the described infiltration of monocytes and leucocytes in these CSF-filled spaces in several models of peripheral inflammation such as hepatic-inflammation^174^, LPS- and complete Freund adjuvant-induced inflammation^175,176^. Here the increased level of F4/80^+^ macrophages within CP and leptomeningeal compartment lining the subarachnoid space in MC38 corroborates the BCSFB inflammation responsible for myeloid recruitment whatever the cancer type. In agreement, accumulation of peripheral macrophages and border-associated macrophages at the CP were recently involved in the extranormal CSF production by CP epithelial cells expressing TLR-4, and activating TNF-receptor-associated kinase SPAK, regulatory scaffold of a multi-ion transporter protein complex^177^. The resulting excess of fluid should lead to ventricles enlargement and harmful pressure on the brain’s tissues sufficient to brain injury or cognitive dysfunctions^178,179^. We have previously demonstrated that depletion strategies of homing macrophages in CP in a context of sub-arachnoid hemorrhage were beneficial to normalize ventriculomegaly^180^, a systemic injection of clodronate-liposomes was used to deplete peripheral macrophages. This procedure normalizes ventricular volume in cancer-bearing mice suggesting that systemic inflammation in both B16F10 and MC38 can lead to cognitive impairment by interfering with CSF fluid dynamics *via* recruitment of macrophages to inflammatory CP. Peripheral origin of invading F4/80^+^ myeloid cells/macrophages was supported by the unchanged level of macrophages in cancer-naïve mice after clodronate liposome administration.

In addition to CP and subarachnoid meningeal space, peripheral inflammation has been shown to promote cluster formation of myeloid cells in the VI, a CSF-filled *velae* in continuity of the choroidal tissue of the third ventricle, formed by networks of pia and arachnoid membranes and anatomically close to the hippocampus^181,182^. In line with the connection between systemic inflammation and hippocampus neuroinflammation and cognitive deficits^183–185^, the observed F4/80^+^ macrophage accumulation in the VI structure of B16F10- and MC38-bearing mice questions how cancer impacts VI and the hippocampal area and contributes to altered behavior. Among all factors involved in cognitive functions, IL-4 secretion by resident patrolling T lymphocytes in leptomeninges helps the normal functioning of the neurogenic niche, in the presence of pro-inflammatory (rather M1) macrophages^186–188^. The decrease in number of BrdU^+^ NPCs in the hippocampal DG measured in both B16F10- and MC38-bearing mice first indicates a strong correlate with the concept of “all-cancer-induced cognitive impairment”. Second, the similar inflammatory (M1) CD206^-^MHC-II^+^ myeloid cells status in the VI between B16F10 or MC38 and PBS mice but the high density of immunosuppressive (M2) CD206^+^MHC-II^-^ myeloid cells specifically in MC38-bearing mice, indicates that NPCs decreased proliferation cannot be explained by the disequilibrium between CD4 TL and M2 macrophages in VI, but maybe to a decreased hippocampal IL-4, according to a study on ageing-related learning and memory deficits relayed by reduced stimulation of IL-4 receptors expressed by hippocampal neurons^189^. However, the M2-like macrophage population in the VI of MC38 would add deleterious mechanisms in the hippocampus relaying for example anxiety- and depressive-like behaviors, by competing with CD4^+^TL-promoting neurogenesis and BDNF production. The absence of F4/80^+^ macrophages in the hippocampal parenchyma of cancer-bearing and cancer-naïve mice, therefore does not support a “parenchymal-contact” based hypothesis in which myeloid/macrophage cells invade the neural parenchyma and interact directly with neurons and/or glial cells^190^. Thus, the major indirect contributor of the B16F10 and MC38-altered neurogenesis is the anti-inflammatory Arg1^+^Iba1^+^ transition towards Iba1^+^CD68^+^ reactive microglia relaying a local neuroinflammation in the hippocampus.

Cancer-associated pro-inflammatory cytokines drive neuroinflammation by compromising the BBB integrity, then allowing some cytokines to interact with the brain’s resident immune cells, and potentially interfere with behavioral functions^191–193^. BBB leakage has already been described in association with plasma IL-1β, IL-6 and MCP-1^194,195^, we thus investigated the BBB permeability in cancer-bearing mice and whether it may be associated with upregulation of inflammatory gene expression in key brain areas involved in activity, emotional reactivity, and cognitive functions. Interestingly, it has already been demonstrated that circulating IL-6 subsequent binding to its IL-6R receptor on hippocampal endothelial cells induces BBB disruption and hippocampal reactive astrogliosis in B16F10 mice^196^, thus this could drive cognitive dysfunctions and resignation observed in cancer-bearing mice. Here we also show a BBB permeability of Dextran_488_ in both B16F10- and MC38-bearing mice, but according to immunofluorescence imaging from brain sections, no vascular leakage was observed within the hippocampus, suggesting that hippocampal microglial reactivity is not driven by an altered BBB, corroborated by the absence of increased systemic level of IL-6 in B16F10 mice, and that cognitive and emotional impairment would be associated with altered BBB in other brain areas. Cortical lesions were shown to affect rodent cognitive long-term memory measured in the NORT ^197–199^. In addition, the infralimbic cortex is involved in the modulation of visceral functions related to emotional inputs, and the medial prefrontal cortex stimulation exabit antidepressant-like effect in the FST^200^. In this study, we imaged a vascular leakage in pre-frontal and somato-motor cortex, as well as in the in cerebellum. According to this observation, IL-1β has been described in pre-frontal cortex of cancer bearing mice in association with fatigue and depressive-like behaviors^150,201^ and alteration of cerebellar architecture can contribute to cognitive and emotional alterations in rodents in association to increase in blood inflammatory TNF-α and MCP-1 mediators^202^ while reduced numbers and size of Purkinje neurons have been described in cancer-bearing mice^203^. Together, our data suggests that main cancer effects are manifested at the level of the brain barriers including BBB to compromise cognitive functions and emotional reactivity in different brain areas.

Peripheral inflammatory mediators can gain brain tissues through tight-junctions-free brain areas such as circumventricular organs and meningeal dura mater^204^. Recent studies have demonstrated a close relationship between the meningeal immune response and CNS functions, and manipulating dural sinuses immunity has been shown to shape neuronal function and associated rodent behaviors^205–208^. In a context of peripheral cancer, the dura mater immune cell composition potentially regulating CNS functions was not previously described. By means of a panel expression of inflammatory gene mRNAs, we here highlight that STING is upregulated in dural brain tissue from both B16F10- and MC38-bearing mice. The cGAS/STING pathway is known to recognize and respond to cytosolic DNA, indicative of cell damage or infection. Activation of cGAS/STING DNA sensing pathway in tumoral and non-tumoral cells has been recently reported to play key role in anti-tumor responses in B16F10 and MC38-bearing mice models, suggesting that STING upregulation in dural meninges of cancer-bearing mice can originate from the TME^209^. But STING activation classically results in the phosphorylation and production of INFγ and downstream interferon stimulatory genes, as pro-inflammatory signature^210^. In the brain, microglia is the main cell type expressing STING, an inflammatory phenotype being responsible of cognitive and motor performances decline associated with ageing^211^. Microglia being absent of the dura mater meningeal compartment, we thus questioned the contribution of peripheral immune cells in re-filling dural immune environment as previously studied in homeostasis^212^ and LPS-induced sepsis-associated encephalopathy conditions^213^. In this latter, monocytes and macrophages, but not T and B lymphocytes, colonized dura mater meninges, suggesting that first changes in dural immune composition concern peripheral myeloid cells stimuli^213^. In our study, concomitant with STING upregulation, mRNA encoding CD80 expressed by inflammatory macrophages and CD4 expressed by T helper lymphocytes, or CD19 expressed by B lymphocytes and chitinase-3-like-1 gene (Chil3) secreted by macrophages and neutrophils are detected in dura of MC38- and B16F10-bearing mice, respectively^214,215^, suggesting infiltration of different immune cells depending on the immuno-inflamed or immune-desert cancer. By using FACS analysis of CD45^+^ immune cell populations, we confirmed that more CD11b^+^LY6G^-^Ly6C^-^MHCII^+^ pro-inflammatory macrophages and CD3^+^CD4^+^ T helper lymphocytes were found in blood of MC38-bearing mice and increased number of CD11b^+^Ly6g^+^Ly6C^-^MHCII^-^ neutrophils are detected in blood of B16F10- and MC38-mice compared with PBS mice. Interestingly, in cancer patients, transcriptional changes of cGAS-STING have been recently reported to occur in circulating PBMCs^216,217^, suggesting that STING expression may be a general cancer signature of circulating reprogramming immune cells, able to infiltrate the dural tissue. However, the source of immune supply of dura meninges may also results from B lymphocytes^218^ and myeloid cells^219^ migration from the skull bone barrow. Indeed, CSF would instruct cranial hematopoiesis *via* direct contact of dural vessels and *calvaria* cavity-bone-marrow^220^, thus the differential blood immune cell composition in B16F10- and MC38-bearing mice by modifying the CSF composition, may regulate cranial hematopoiesis which in turn can supply the dural layer with skull-derived immune cells in cancer mice.

Anti-tumoral response to ICIs largely depends on pre-existing TME immune infiltration^57,221,222^. Despite therapeutic success for only a portion of patient populations with immune-inflamed cancer, the occurrence irAEs complicates patient management and provoke treatment discontinuation^223^, even if irAEs are intrinsically associated with a better anti-tumoral response. This is also revealed through peripheral biomarkers shown as good markers of anti-tumoral response, such as Neutrophil-to-Lymphocyte ratio and Platelet-to-Lymphocytes ratio^224–227^, but also associated with increased risk of irAEs development in cancer patients^228^. Given that the immune response plays an important role in the development of cognitive disorders^229,230^, and that good immune stimulation should be a prerequisite for ICI-antitumor response and development of ICI-related toxicities, in this study we aimed at understanding the impact of ICI on cognitive functions considering the immune-inflamed and immune-desert status of cancers.

Here in cancer-naive mice peripheral administration of anti-PD-1, anti-PD-L1 or their combination did not impact cognitive functions or emotional reactivity, showing that in absence of cancer, ICIs should not alter behaviors. However, cancer-induced cognitive impairment was exacerbated in immune-inflamed MC38-PD-1 compared with IgG mice while no behavioral deficit occurred with anti-PD-1 in B16F10-bearing mice. Similar results in a murine model, such as anxiety-like behavior and decreased cognitive performances, have been previously described when anti-CTLA-4^65^ was tested only in combination with cancer likely suggesting that immune stimulation prior to ICI treatment is an obligatory condition of ICI-induced on cognitive dysfunctions. This effect of anti-CTLA-4 was predictable, since expression of CTLA-4 outside the immune system has been described, in addition to potential “off-target” action of anti-CTLA-4 due to auto-immune reactivation against neuronal antigens, driving endocrine toxicity in patients^231^. Little is known about PD-1 or PD-L1 expression in the brain at the physiological state^232,233^, however, their upregulation in several cell types during inflammation has been well characterized^234,235^. It was recently demonstrated that neuronal PD-1, at physiological state, has a major role in cognitive performances *via* the regulation of neuronal excitability and synaptic ability^233^. Given the central expression of ICIs targets and cancer-associated BBB permeability, we thus hypothesized that systemic administered ICIs could drive cognitive and depressive-like behavior through direct action on brain targets. To test this hypothesis, an anti-IgG2a-fluorescently coupled antibody directed against anti-PD-1 or anti-PD-L1 was used to map a potential distribution of ICIs in the brain parenchyma. We do not detect these monoclonal antibodies in the hippocampus or the prefrontal cortex, excluding BBB penetration and distribution toward neuronal brain targets, even if efficacy of ICIs has been proven in context of brain tumors or central metastasis^236–238^. This central efficacy should be indirect as recently suggested with anti-CTLA-4 providing therapeutic efficacy on central malignances *via* brain infiltration of peripheral T lymphocytes^239^. This supports anti-PD-1 neurocognitive impact likely involving peripheral activation of the immune system in MC38-bearing mice.

Only a moderate exacerbation of peripheral inflammation after ICIs treatment was observed, with increased plasma levels of MCP-1 and IL-1α in MC38-mice after PD-1 and/or PD-L1 treatment, and of IL-1β in B16F10 mice after PD-1 administration. These changes were associated with a slight higher level of circulating ICAM-1 in MC38-PD-1 mice compared with MC38-IgG mice, thus in the whole stressing only subtle exacerbation of systemic and vascular inflammation after PD-1/PD-L1 whatever the cancer profile. Interestingly, compared with the cancer condition, BBB permeability was only exacerbated in mice treated with PD-L1 and bearing either B16F10 or MC38. Such observation is strongly related to the ICAM-1+ endothelial activation of DG hippocampal vessels. We suppose that expression of IL-1α and IL-1β IL-RI receptor in mouse cerebral endothelial cells relay endothelial activation in the presence of PD-L1^240–242^, making the vessel endothelium a direct target of peripheral immune stimulation associated with ICIs. This agrees with recent clinical observations showing specific alteration of plasma levels of IL-1β and IL-1α in cancer patients treated by ICIs with poorer cognitive performances. Considering that endothelial activation can be also associated with cognitive decline and mood disorders in patients with several psychiatric disorders^243^, we propose that once stimulated after PD-1 and more markedly after PD-L1 or in the presence of MC38, systemic cytokines induce brain vascular inflammation, mistranslated by ICAM-1 or soluble sICAM-1systemic level. Among pro-inflammatory mRNA cytokines, CCL19 has been found overexpressed in intimal myeloid cells undergoing reverse transendothelial migration into the arterial circulation after systemic stimulation of pattern-recognition receptors resulting from infection^244^. Here, we originally showed that CCL19 appears intensely expressed in proximity with DG hippocampal vessels after treatment of MC38-mice with ICIs. CCL19^+^Lectin^+^ vessels are normally located in lymphoid structures in which they regulate homing of T and B cells^245^. However, in a mouse model of EAE, increased expression of CCL19 by brain endothelial cells of a specific subset of inflamed vessels was demonstrated to be necessary for T lymphocytes recruitment and adherence^246^, thus suggesting that ICIs may promote CCL19 expression in myeloid cells and/or inflamed vessels essential for immune cells transmigration. We indeed highlight infiltration of CD3^+^ T lymphocytes in the hippocampal parenchyma of PD-1/PD-L1 MC38-bearing mice. As in a mouse model of AD in which hippocampal T cells infiltration is associated with neuroinflammation and diminished spatial memory performances^247^, we suggest the exacerbation of cognitive impairment and the decreased hippocampal NPC proliferation in MC38-bearing mice treated with anti-PD-1/ant-PD-L1 can be driven by T lymphocytes infiltration in the hippocampus. The absence of modification of VI meningeal density of pro- or anti-tumoral macrophages let us exclude a myeloid contribution in favor of a leucocyte-mediating mechanism. In agreement, T lymphocytes infiltration has already been demonstrated to impact adult hippocampal neurogenesis in a context of aging^248^ and of post-operative cognitive decline^249^. These data indicate that for immune-inflamed cancer, ICIs induce vascular inflammation, T lymphocyte infiltration, altered neurogenesis likely responsible for cognitive dysfunctions and depressive-like behaviors.

Interestingly, whatever the immune-inflamed or immune-desert cancer, mice treated with the anti-PD-L1 specifically exhibit diminished memory performances as well as anxiety-like behaviors, in association with a higher proportion of circulating TCRγδ lymphocytes (γδT) as well as distribution within the BBB in line with hyperpermeability. Interestingly, this specific γδT subpopulation have been pointed out as a critical player in progression of several central diseases by promoting BBB permeability *via* IL-17 secretion^250–252^. In our study, despite the increased circulating number of γδT lymphocytes, we do not detect increased levels of IL-17 in plasma in cancer-bearing mice treated with anti-PD-L1. This can be explained the a rapid activation with elevated expression of PD-1 for instance can be followed by exhaustion of γδT, characterized by decreased cytokine production and functional impairment^253^. We further demonstrated that anti-PD-L1 exacerbates cognitive and anxiety impairment *via* γδT infiltration in CSF-filled spaces of B16F10 and MC38-mice. It has been already shown that meningeal resident γδT populate the meninges at perinatal stages, they are maintained by local self-renewal during adulthood and are essential for regulation of CNS functions^205^. In fact, during homeostasis, meningeal γδ LTs regulate short-term memory *via* secretion of IL-17a necessary for glutamatergic synaptic plasticity and BDNF secretion by glial cells^254^. Inversely, physiological secretion of IL-17 by meningeal γδ LT has been associated to promote anxiety-like behaviors^205^ *via* stimulation of IL-17R-expressing glutamatergic cortical neurons. During neuroinflammation, accumulation of γδ LT in the meninges and subsequent IL-17 hypersecretion has been demonstrated to trigger the onset of cognitive decline in a murine model of AD^250^. In our study, we hypothesize that during cancer-driving neuroinflammation, accumulation of IL-17-γδ T-secreting cells in the leptomeninges could imbalance homeostatic IL-17-dependent cerebral mechanisms and potentially deliver INFγ, leading to cognitive dysfunctions and anxiety-like-behaviors. The origin of these γδT lymphocytes after PD-L1 treatment remains unclear. In peripheral tissues, γδ T cells represent a minor part (1–5%) of the circulating T cell compartment^255^, as a matter of fact γδT resident cells are found in much higher proportion in epithelial tissues, such as the reproductive tract, skin epidermis and gastrointestinal tract^256,257^. Interestingly, gut resident γδ T cells have been demonstrated to be enhanced after intestinal dysbiosis thus infiltrating the leptomeninges *via* a direct gut-brain trafficking in a context of ischemic stroke, without modified γδ T blood proportion^206^. As dysbiosis is frequently described in cancer patients treated with ICIs and developing irAEs^258,259^, we must ask whether PD-L1-induced intestinal dysbiosis drives γδT leptomeningeal infiltration in cancer. Onother hypothesis is based on the rapid proliferation of blood γδT cells previously observed in response to phospho-antigens expressed at the cancer cells membrane^260,261^, more in line with the observed increased blood γδT in our cancer mice.

To demonstrate the causal link between circulating γδT and behavioral changes induced by ant-PD-L1, we tested an anti-γδTCR in cancer MC38 mice treated with anti-PD-L1. The peripheral γδT immunoneutralization led to concomitant decrease of γδT in the blood and within the CP and subarachnoid space. This suggests that circulating γδT resulting from the treatment with PD-L1 may cross the BCSFB independently of the cancer immune status. Here, γδT showed no contribution on PD-L1 therapeutic efficacy, but clinical studies provided evidence that high γδT frequency in tumor infiltrates^262–265^ and in blood^266^ were shown to correlate with better clinical outcome in different malignancies after ICIs. Together, the present data establish that γδT blood-brain barrier infiltrations represent an underlying mechanism of cognitive and emotional alterations associated with anti-PD-L1 immunotherapy in immune-inflamed or immune-desert cancers, γδT consequently represents a key, to be verified, actionable and relative safe target to prevent ICI-induced CRCI and to maintain cancer patients’ quality of life during cancer immunotherapy.

There are some limiting factors that must be considered when examining our data. First syngeneic models used in this study significantly differ from tumors generated spontaneously in term of tissue microenvironment, which is not the organ specific environment, particularly in term of kinetic of growth which is much more rapid than spontaneously developing tumors. Our priority was to characterize the impact of tumor-associated immune and inflammatory responses on cerebral functions, so the choice of a subcutaneous tumor model allowed us for precise and reliable comparison of all groups, with behavioral test done when tumoral volume mass was comparable among all mice bearing different cancers. Second, we cannot totally exclude a potential metastatic process leading to cancer cells infiltration into the brain, driving immune responses. Here, we never detect anaplastic mass in the number of analyzed brain slices, and despite the potential metastatic profile of B16F10 cell line or the less frequent MC38-derived metastasis, metastasis does not typically occur after subcutaneous implantation^267–270^. Third, circulating immune cells in the context of cancer, and after ICI treatment, should have infiltrate and influence functions of other organs, thus influencing performances in behavioral tests. For example, huge infiltration of T lymphocytes has been demonstrated to drive cardiac toxicity in ICI-treated cancer patients^271^. Moreover, pharmacokinetic distribution studies in mice have shown that excessive accumulation of ICIs after systemic administration can occur in several peripheral organs contributing to the development of peripheral toxicities^272–274^. Fourth, in our study, only male mice were used. Since female hormones have been demonstrated to influence immunity^275^, immunotherapy response^276^ and behavior^277^, sex would be a potential confounding factor. In agreement, sex-dependent differences in the context of CRCI have already been shown in colon cancer patients, with women more likely to develop cognitive dysfunctions than men^278^, thus reinforcing the need for testing both female and male mice in future studies.

In conclusion, with the growing use of immunotherapy strategies in many types of cancers, the severity of cognitive impairment in patients treated by immunotherapy is a major issue. Our results demonstrate an increased neurotoxicity of immune-inflamed cancer and specific impact of PD-L1-promoting γδT blood and brain infiltration, on cognitive functions and anxiety-like behavior whatever the cancer type. This work should provide new biomarkers of CRCI induced by cancer and cancer combined with ICIs, and help the future prevention of the long-term consequences of these treatments on brain functions enabling progress in survival, QoL, return at work and/or autonomy of cancer patients.

## Acknowledgements

Images were obtained on PRIMACEN, the Cell Imaging Platform of Normandy, IRIB, Normandy Rouen University, France. We thank Arnaud Arabo and Julie Maucotel for animal housing and care and for access to the behavioral equipment of the Biological Resources Department (Normandie Rouen University, UMRS Heracles, France). This work was supported by the Canceropole Northwest, the Institut National de la Santé et de la Recherche Médicale (INSERM, grant number U1245 to HC); Normandy Rouen University (Grant CBG to HC); the Regional Council of Normandy and the European Community FEDER program (Europe gets involved in regional development through the ERDF program) (Grants CANCER-COG and RIN/FEDER Tremplin 3 R); and grants from Ligue contre le Cancer Normandie. C.N. was supported by the Regional Council of Normandy and the European Community FEDER program.

## Contributions

C.N. performed all surgeries, designed and performed image acquisition studies, performed the main data analysis and wrote the drafted manuscript. H.C., M.P., M.D., C.N., P.N., L.D., P-K.D. performed cytokines Elisa Assays, imaging experiments and data acquisition, and all data analyses. M.D, I.J., P.H. helped with behavioral studies and their interpretation. G.R., S.A. developed and customized and helped in analysis of flow cytometry. H.C., C.N., D.V., P.L., designed, performed the transcriptomic study and mRNA analyses. H.C. conceived the study design and data analysis. C.N., H.C and O.W. worked on the Ingenuity Pathway analysis. O.W., F.J., P.H. provided editorial review and advice. H.C. obtained funding, conceived the idea, experimental design, analysis and interpretation, project supervision, and wrote the drafted and final manuscript.

## Data availability

The data are available within the Article, Supplementary Information or Source Data file. Source data will be provided with this paper. The experimental data that support the findings of this study are available in the Supplementary information file.

