## Supplementary material for "Anti-PD-1/PD-L1 Therapy Triggers Cognitive Deficits and Anxiety-Like Behaviors Through Tumor-Initiated Neuroinflammatory Niches in Male Mice": supp figures

**a** Correlations between vascular and immune markers

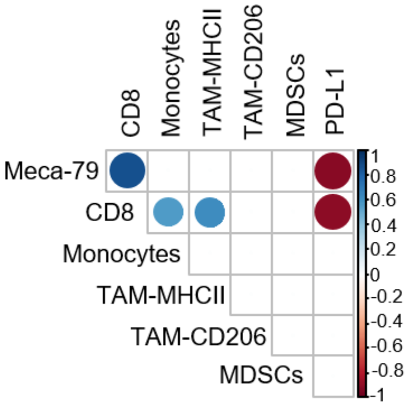


**b** Immunoscore

| Treatment | CD8⁺ cells (n/μm²) | Score | HEVs (MECA79⁺ cells, % of total area) | Score | Monocytes (CD11b⁺GR1⁻ cells · n/mm²) | Score | MHCII⁺ cells (n/mm²) | Score | Immunoscore |
|---|---|---|---|---|---|---|---|---|---|
| B16F10 | 9.013 | 0 | 0.960 | 0 | 72.100 | 1 | 72.101 | 0 | **1** |
| | 6.008 | 0 | 3.005 | 1 | 36.050 | 0 | 130.683 | 1 | **2** |
| | 12.017 | 1 | 3.270 | 2 | 18.025 | 0 | 78.860 | 0 | **3** |
| | 12.017 | 1 | 1.660 | 0 | 45.063 | 0 | 114.911 | 0 | **1** |
| | 12.017 | 1 | 2.080 | 0 | 54.070 | 1 | 90.126 | 0 | **2** |
| B16F10-Ova | 24.034 | 2 | 3.210 | 1 | 207.290 | 2 | 207.290 | 1 | **6** |
| | 12.017 | 1 | 4.856 | 3 | 351.400 | 3 | 259.113 | 1 | **8** |
| | 15.021 | 2 | 2.947 | 1 | 216.302 | 2 | 196.024 | 1 | **6** |
| | 0.000 | 0 | 3.113 | 1 | 36.050 | 0 | 459.643 | 3 | **4** |
| | 3.004 | 0 | 1.250 | 0 | 261.360 | 3 | 416.833 | 2 | **5** |
| MC38 | 75.105 | 3 | 5.470 | 3 | 234.320 | 2 | 414.580 | 2 | **10** |
| | 45.063 | 3 | 5.720 | 3 | 243.340 | 2 | 351.492 | 2 | **10** |
| | 51.072 | 3 | 3.450 | 2 | 171.230 | 1 | 259.113 | 1 | **7** |
| | 27.038 | 2 | 3.770 | 2 | 360.500 | 3 | 585.820 | 3 | **10** |
| | 30.042 | 2 | 4.136 | 2 | 126.176 | 1 | 675.946 | 3 | **8** |
| **Score** | | | | | | | | | |
| Maximum | 75.105 | 3 | 5.720 | 3 | 360.500 | 3 | 675.946 | 3 | |
| 75% percentile | 30.042 | | 4.136 | | 243.340 | | 416.833 | | |
| 75% percentile | 30.042 | 2 | 4.136 | 2 | 243.340 | 2 | 416.833 | 2 | |
| Median | 12.017 | | 3.210 | | 171.230 | | 259.113 | | |
| Median | 12.017 | 1 | 3.210 | 1 | 171.230 | 1 | 259.113 | 1 | |
| 25% percentile | 9.013 | | 2.080 | | 45.063 | | 114.911 | | |
| 25% percentile | 9.013 | 0 | 2.080 | 0 | 45.063 | 0 | 114.911 | 0 | |
| Minimum | 0.000 | | 0.960 | | 18.025 | | 72.101 | | |

**c** Immunosuppressive Score

| Treatment | PD-L1 (% of total cells) | Score | MDSCs (n/mm²) | Score | CD206 (n/mm²) | Score | Immunosuppressive score |
|---|---|---|---|---|---|---|---|
| B16F10 | 94.620 | 3 | 126.177 | 2 | 423.593 | 2 | 7 |
| | 88.050 | 1 | 180.252 | 3 | 419.087 | 2 | 6 |
| | 88.660 | 1 | 81.114 | 1 | 675.946 | 3 | 5 |
| | 91.190 | 2 | 117.164 | 2 | 648.908 | 3 | 7 |
| | 67.660 | 1 | 99.139 | 2 | 225.315 | 0 | 3 |
| B16F10-Ova | 96.610 | 3 | 135.189 | 2 | 430.352 | 3 | 8 |
| | 91.620 | 2 | 234.328 | 3 | 351.492 | 1 | 6 |
| | 93.690 | 2 | 153.214 | 3 | 247.847 | 0 | 5 |
| | 92.410 | 2 | 162.227 | 3 | 254.606 | 1 | 6 |
| | 94.790 | 3 | 63.088 | 1 | 414.580 | 2 | 6 |
| MC38 | 38.050 | 1 | 18.025 | 0 | 123.923 | 0 | 1 |
| | 13.800 | 0 | 45.063 | 1 | 344.732 | 1 | 2 |
| | 10.320 | 0 | 18.025 | 0 | 259.113 | 1 | 1 |
| | 15.890 | 0 | 54.076 | 1 | 180.252 | 0 | 1 |
| | 14.780 | 0 | 36.050 | 0 | 403.314 | 2 | 2 |
| **Score** | | | | | | | |
| **Maximum** | 96.610 | 3 | 234.320 | 3 | 675.946 | 3 | |
| **75% percentile** | 93.690 | | 153.210 | | 423.593 | | |
| **75% percentile** | 93.690 | 2 | 153.210 | 2 | 423.593 | 2 | |
| **Median** | 88.660 | | 99.138 | | 351.492 | | |
| **Median** | 88.660 | 1 | 99.138 | 1 | 351.492 | 1 | |
| **25% percentile** | 15.890 | | 45.063 | | 247.847 | | |
| **25% percentile** | 15.890 | 0 | 45.063 | 0 | 247.847 | 0 | |
| **Minimum** | 10.320 | | 18.025 | | 123.923 | | |

**Figure S1. Immune cells and markers contributing to immune- or immunosuppressive scores calculation. a.** Correlation between immunostaining items used to define immunoscore and immune-suppressive score. Heatmap of Kendall correlation coefficients; only significant correlations are displayed (adjusted p-value <0.05) and values obtained in B16F10, B16F10-Ova and MC38 have been pooled. **b.** Table recapitulating the immunoscore calculation methods in B16F10, B16F10-Ova and MC38 tumor slices. Densities of $CD8^+$ T lymphocytes, Meca-$79^+$ HEVs, $CD11b^+$ monocytes and $MHCII^+$ anti-tumoral TAMs in tumor slices were converted into percentile values. Percentiles values were then converted into score values with score 0 attributed to percentile values ranking from (0-25%), score 1 (25-50%), score 2 (50-70%), score 3 (75-100%). The mean of each score was then calculated to generate immunoscore. **c.** Table recapitulating the immunosuppressive score calculation methods in B16F10, B16F10-Ova and MC38 tumor slices. Densities of $CD206^+$ pro-tumoral TAMs, $CD11b^+GR1^+$ MDSCs and percentage of PD-$L1^+$ cells in tumor slices were converted into percentile values. Percentiles values were then converted into score values with score 0 attributed to percentile values ranking from (0-25%), score 1 (25-50%), score 2 (50-70%), score 3 (75-100%). The mean of each score was then calculated to generate immune-suppressive score. CD11b: beta 2 integrin adhesion molecule, CD206: cluster of differentiation 206, CD8: cluster of differentiation 8, GR1: protein gamma response 1, HEVs: high endothelial venules, MDSCs: myeloid-derived suppressor cells, Meca-79: peripheral node adressin, MHCII: major histocompatibility complex 2, PD-L1: Programmed death-ligand 1, TAM: tumor associated macrophages. PD-L1: programmed death-ligand 1, TAM: tumor associated macrophages.

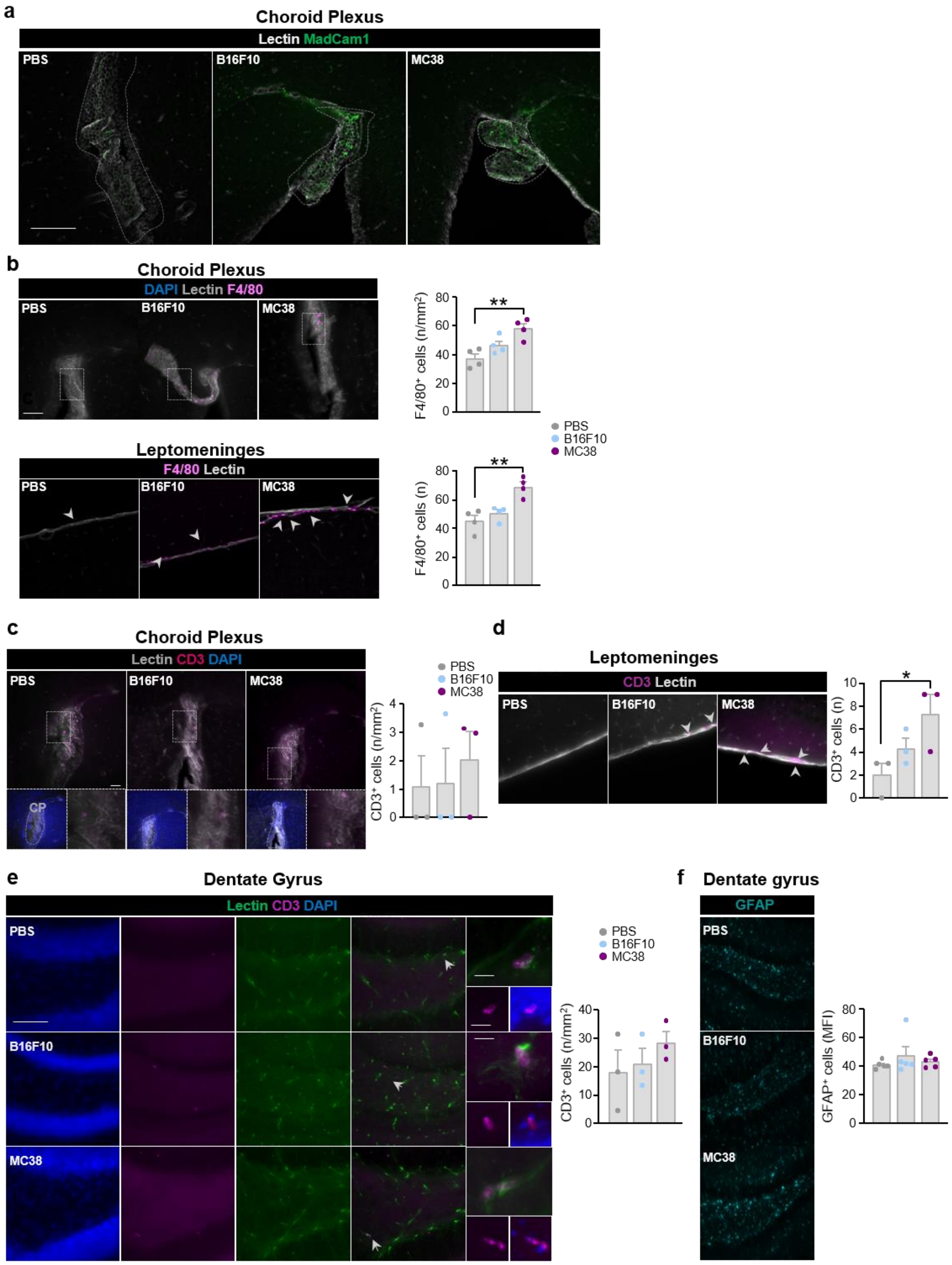


**Figure S2. Inflammatory markers and neuroinflammation in Choroid plexus, leptomeninges and hippocampus in immuno-inflamed and immuno-desert cancer-bearing mice. a.** Representative images of MadCam-1$^{+}$ (green) and Lectin$^{+}$ (grey) immunoreactivity in CP of PBS-control or B16F10 and MC38 mice. Scale bar: 100 µm. **b.** Representative images of Lectin (gray) and F4/80 (magenta) immunoreactivity in CP and leptomeninges of PBS-control or B16F10 and MC38 mice. Scale bar: 50 µm. Statistical analysis was performed by using one-way ANOVA with Bonferroni correction for multiple comparisons. Data are represented as bars with symbols for individual data points and they are expressed by mean ± SEM, n=4, * p<0.05, ** p<0.01. **c.** Representative images and statistical quantification of number of T lymphocytes ($CD3^+$, magenta) in choroid plexus ($Lectin^+$, grey) of B16F10- and MC38-bearing mice compared to PBS-injected controls. Boxed areas represent lateral ventricular localization (DAPI, blue) of $Lectin^+$ CP and magnification of $CD3^+$ T lymphocytes (magenta). On the right, bars indicate statistical comparison of number of T lymphocytes ($n/mm^2$, density) in B16F10- and MC38-mice when compared to PBS-mice. Scale bar: 50 µm, zoom 10 µm. **d.** Representative images and statistical quantification of number of T lymphocytes ($CD3^+$, magenta) in leptomeninges ($Lectin^+$, grey) of B16F10- and MC38-bearing mice compared to PBS-injected controls. White arrows represent $CD3^+$ T lymphocytes (magenta). On the right, bars indicate statistical comparison of number of T lymphocytes ($n/mm^2$, density) in B16F10- and MC38-mice when compared to PBS-mice. Scale bar: 50 µm. **e. and f.** Representative images of T lymphocytes ($CD3^+$, magenta) cells, vessels (Lectin+, green) and cells nuclei (DAPI, blue) (**e**) or GFAP (cyan, **f**) immunoreactivities in dentate gyrus of PBS and B16F10- and MC38-bearing mice. On the right, bars indicate statistical comparison of number of T lymphocytes (**e**, $n/mm^2$, density) or GFAP (**f**, $n/mm^2$, density) in B16F10- and MC38-mice when compared to PBS-mice. Scale bars: 100 µm, zooms 50 µm and 10 µm. Statistical analysis was performed by using one-way ANOVA or Kruskal-Wallis test with Bonferroni or Dunn's correction for multiple comparison. Data are expressed by mean ± SEM, n=4-5. F4/80: EGF-like module-containing mucin-like hormone receptor-like, GFAP, Glial fibrillary acidic protein, MadCam1: Mucosal vascular addressin cell adhesion molecule 1.

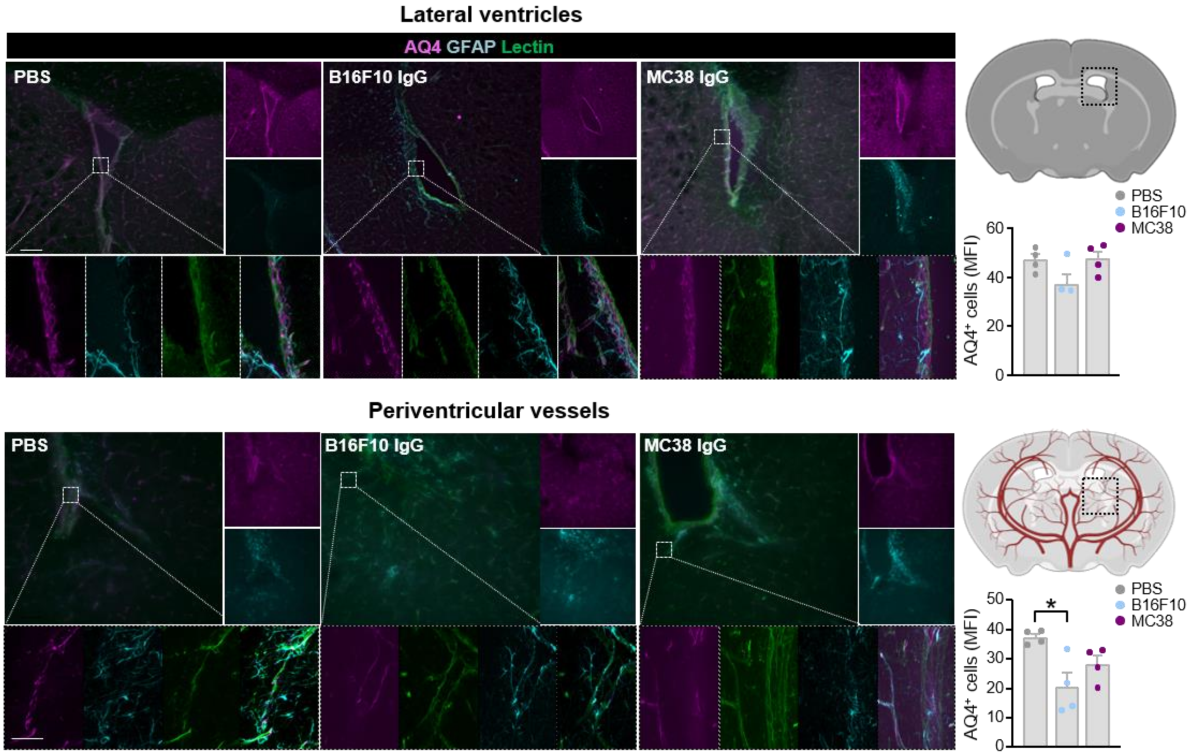


**Figure S3. Aquaporin 4 in lateral ventricles and perivascular vessels in immuno-inflamed and immuno-desert cancer-bearing mice.** On the upper panel, representative immunolabeling of Lectin$^{+}$ (green) vessels, AQ4$^{+}$ (magenta) and GFAP$^{+}$ (cyan) astrocytes in lateral ventricles of PBS, B16F10-IgG, MC38-IgG-mice brain slices. Boxed areas show magnification of AQ4$^{+}$GFAP+ astrocytes. On the lower panel, representative immunolabeling of Lectin$^{+}$ periventricular vessels (green), AQ4$^{+}$ (magenta) and GFAP$^{+}$ (cyan) astrocytes of PBS, B16F10-IgG, MC38-IgG-mice brain slices. Boxed areas show magnification of AQ4$^{+}$GFAP+ astrocytes of Lectin$^{+}$ vessels. On the left, bars of quantification of AQ4 staining intensity expressed as median fluorescence intensity (MFI) in PBS, B16F10-IgG, MC38-IgG-mice lateral ventricles and periventricular vessels. Scale bars: 200 µm, zoom 10µm. Statistical analysis was performed by using one-way ANOVA with Bonferroni correction for multiple comparisons. Data are represented as bars with symbols for individual data points and they are expressed by mean ± SEM, n=4, * $p<0.05$. AQ4: acquaporin 4, CP: choroid plexus, DAPI: 4',6-diamidino-2-phénylindole.

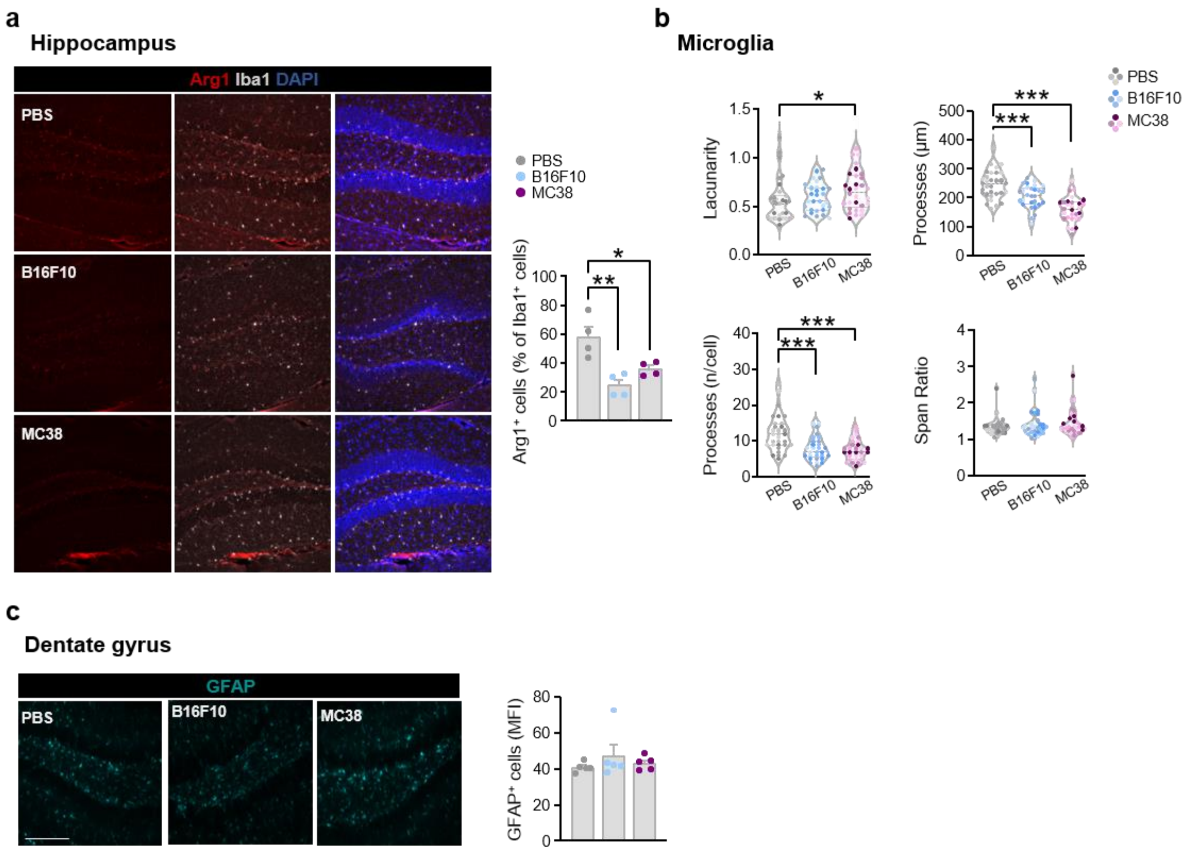


**Figure S4. Microglia reactivity in the hippocampus of immuno-inflamed and immuno-desert cancer-bearing mice. a.** Immunofluorescence and statistical quantification of anti-inflammatory (Arg-1$^{+}$, red) microglial cells (Iba1$^{+}$, grey) in dentate gyrus (DAPI, blue) of B16F10- and MC38- bearing mice compared to PBS-mice. Scale bar: 100 µm. Statistical analysis was performed by one-way ANOVA test with Bonferroni correction for multiple comparisons. Data are represented as bars with symbols for individual data points and they are expressed by mean ± SEM, n=4, * $p<0.05$, ** $p<0.01$. **b.** Quantification of lacunarity, span ratio, number and length of processes of microglial cells of PBS-, B16F10- and MC38-bearing mice. Data are represented as violin-plots with individual points representing individual microglial cell (n=32 for experimental group). **c.** Statistical quantifications (n=4) were performed using the Kruskal-Wallis test with Dunn's correction for multiple comparisons, * $p< 0.05$, *** $p<0.001$. Arg-1: arginase-1, CD3: cluster of differentiation 3, CP: choroid plexus, DAPI: 4',6-diamidino-2-phenylindol, GFAP: Glial fibrillary acidic protein, Iba1: Ionized calcium-binding adaptor molecule 1.

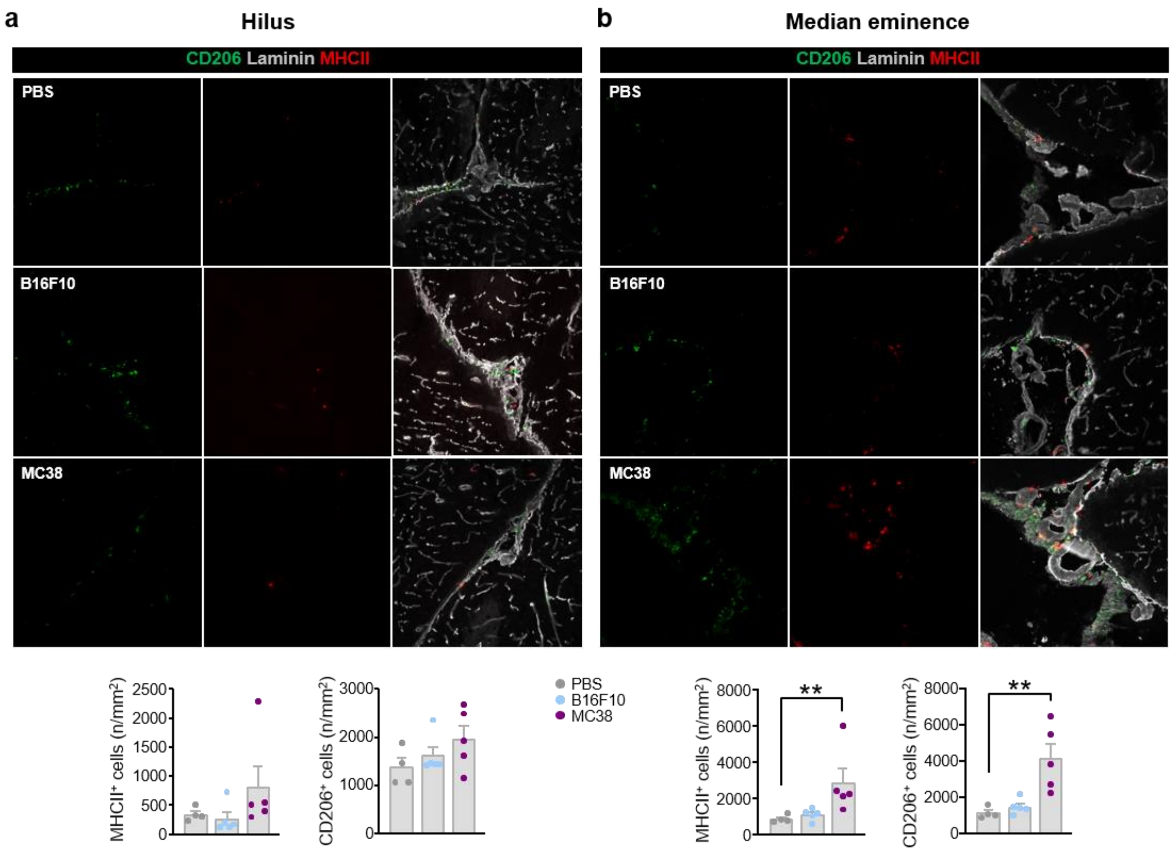


**Figure S5. Quantification of myeloid cells in circumventricular organs of cancer-bearing mice. a.** Representative images and statistical quantification of density (n/mm$^2$) of pro-tumoral (CD206$^+$) and anti-tumoral (MHCII$^+$) myeloid cells staining in perivascular and vascular Laminin$^+$ spaces (grey) of the hilus (**a**) and median eminence (**b**) of B16F10- and MC38-bearing mice compared to PBS-controls. Scale bar: 100 µm. **a. and b.** Statistical quantifications were performed using the One-way ANOVA test or Kruskal-Wallis test with Bonferroni or Dunn's correction for multiple comparisons. Data are represented as bars with symbols for individual data points and they are expressed by mean ± SEM, n=4-5, ** p <0.01. CD206: cluster of differentiation 206, D: day, i.p.: intraperitoneal injection, MHCII: major histocompatibility complex 2.

**a** **Differential analysis of selected genes in meninges of B16F10 *vs* PBS mice by Ingenuity pathway**

| Top Canonical pathways (PBS *vs* B16F10) | | |
|---|---|---|
| Name | p-value | Overlap |
| Systemic lupus Erythematosus in B Cell Signaling Pathway | 1.63E-04 | 0.6% (3/508) |
| Cgas-Sting Signaling Pathway | 3.87E-04 | 1.6% (2/124) |
| IL-17 Signaling | 6.19E-04 | 1.3% (2/157) |
| IL-12 Signaling and Production in Macrophages | 1.06E-03 | 1.0% (2/206) |
| Pathogen Induced Cytokine Storm Signaling Pathway | 2.46E-03 | 0.6% (2/315) |
| **Top Disease and Bio Functions** | | |
| **Diseases and disorders** | | |
| Name | p-value range | # molecules |
| Inflammatory Response | 4.43E-02 – 6.06E-07 | 5 |
| Infectious Disease | 2.48E-02 – 6.12E-07 | 4 |
| Organismal Injury and Abnormalities | 4.74E-02 – 6.12E-07 | 5 |
| Dermatological Disease | 1.64E-02 – 1.55E-06 | 4 |
| Connective tissue disorders | 2.86E-02 – 2.46E-06 | 3 |
| **Molecular and Cellular Functions** | | |
| Name | p-value range | # molecules |
| Cellular Mouvement | 3.33E-02 – 2.58E-08 | 5 |
| Cell-To-Cell Signaling and Interaction | 3.33E-02 – 1.02E-06 | 4 |
| Cellular Development | 4.82E-02 – 1.96E-06 | 4 |
| Cellular Death and Survival | 3.69E-02 – 1.05E-05 | 4 |
| Cellular Compromise | 2.38E-02 – 1.41E-05 | 4 |
| **Physiological System Development and Function** | | |
| Name | p-value range | # molecules |
| Hematological System Development and Function | 4.84E-02 – 2.58E-08 | 5 |
| Immune Cell Trafflicking | 3.33E-02 – 2.58E-08 | 5 |
| Tissue Morphology | 4.04E-02 – 6.06E-07 | 5 |
| Lymphoid Tissue Structure and Development | 4.82E-02 – 1.96E-06 | 5 |
| Connective Tissue Development and Function | 4.80E-02 – 1.67E-05 | 4 |

**b** **Differential analysis of selected genes in meninges of MC38 *vs* PBS mice by Ingenuity pathway**

| Top Canonical pathways (PBS *vs* MC38) | | |
|---|---|---|
| Name | p-value | Overlap |
| Toll-Like Receptor Cascades | 9.95E-08 | 10.3% (3/29) |
| MSP-RON Signaling Pathway | 7.11E-07 | 5.5% (3/55) |
| Neutrophil Extracellular Trap Signaling Pathway | 3.30E-06 | 1.1% (4/351) |
| Neutrophil Degranulation | 7.95E-06 | 0.9% (4/438) |
| Defensins | 9.80E-06 | 14.3% (2/14) |
| **Top Disease and Bio Functions** | | |
| **Diseases and disorders** | | |
| Name | p-value range | # molecules |
| Inflammatory Response | 4.90E-02 – 1.20E-10 | 7 |
| Infectious Disease | 2.00E-02 – 2.97E-10 | 6 |
| Organismal Injury and Abnormalities | 4.96E-02 – 2.97E-10 | 7 |
| Endocrine System Disorders | 2.56E-02 – 3.65E-08 | 5 |
| Gastrointestinal Disease | 4.96E-02 – 3.65E-08 | 7 |
| **Molecular and Cellular Functions** | | |
| Name | p-value range | # molecules |
| Cell-To-Cell Signaling Interaction | 4.32E-02 – 1.20E-10 | 7 |
| Cellular Development | 3.08E-02 – 1.50E-09 | 7 |
| Cellular Growth and Proliferation | 2.84E-02 – 1.50E-09 | 7 |
| Cellular Function and Maintenance | 3.80E-02 – 5.64E-09 | 7 |
| Cell Death and Survival | 4.39E-02 – 3.65E-08 | 7 |
| **Physiological System Development and Function** | | |
| Name | p-value range | # molecules |
| Hematological System Development and Function | 4.56E-02 – 1.20E-10 | 7 |
| Immune Cell Trafflicking | 4.56E-02 – 1.20E-10 | 7 |
| Organismal Survival | 1.06E-03 – 2.59E-09 | 7 |
| Tissue Morphology | 4.39E-02 – 1.67E-08 | 7 |
| Lymphoid Tissue Structure and Development | 4.39E-02 – 6.37E-08 | 7 |

**Figure S6. Signaling pathway enhanced in dura mater of cancer-bearing mice.** Canonical pathway analysis, disease and function were studied by using the IPA system. The P-value of overlap <0.05 was set as the threshold and the significance level is obtained by Fisher's exact test at the right tail. IPA: ingenuity pathway analysis.

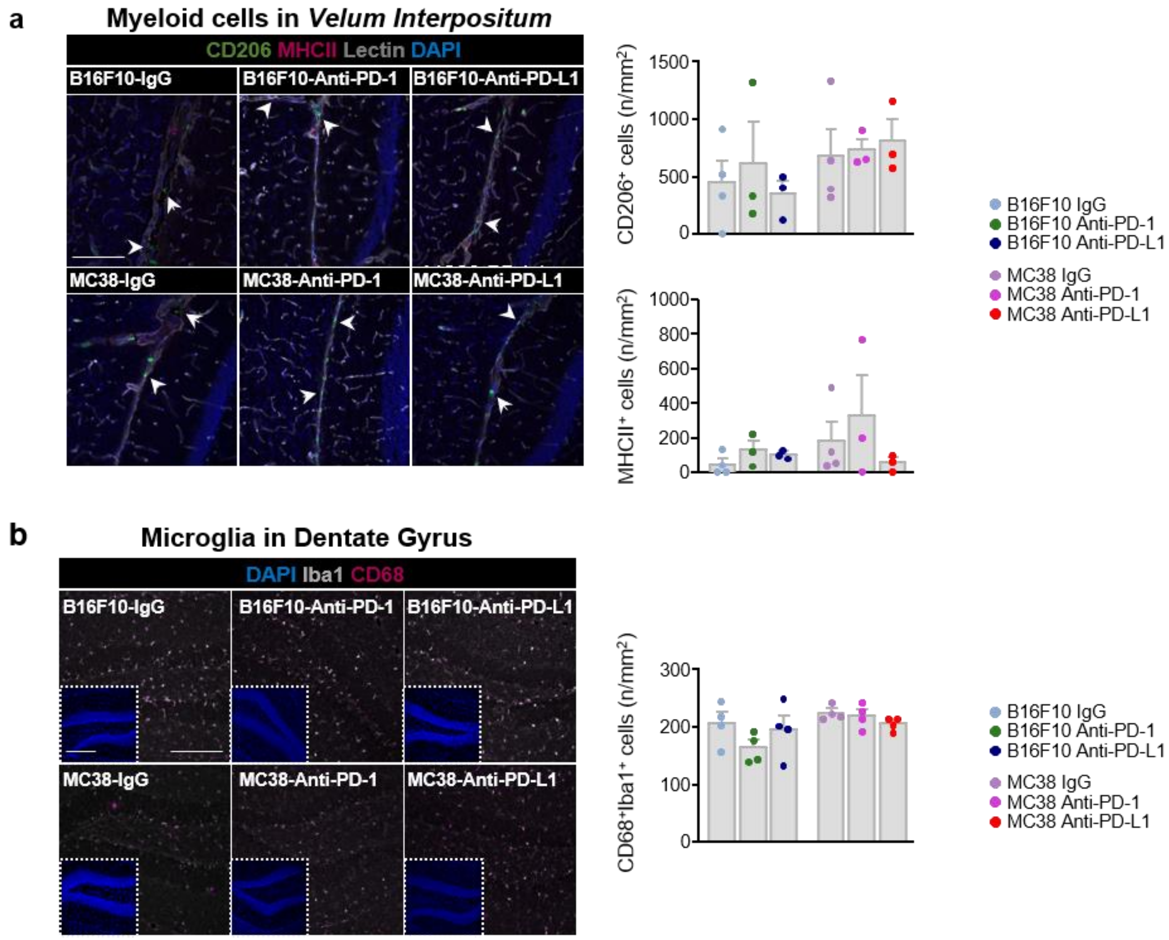


**Figure S7. Quantification of myeloid cells in circumventricular organs of cancer-bearing mice. a.** Representative images and statistical quantification of density (n/mm$^2$) of pro-tumoral (CD206$^+$) and anti-tumoral (MHCII$^+$) myeloid cells staining in perivascular and vascular Laminin$^+$ spaces (grey) of the hilus (left panel) and median eminence (right panel) of B16F10- and MC38-bearing mice compared to PBS-controls. Scale bar: 100 µm. Statistical quantifications were performed using the One-way ANOVA test or Kruskal-Wallis test with Bonferroni or Dunn's correction for multiple comparisons. Data are represented as bars with symbols for individual data points and they are expressed by mean ± SEM, n=4-5, * $p<0.05$, ** $p<0.01$. CD206: cluster of differentiation 206, D: day, i.p.: intraperitoneal injection, MHCII: major histocompatibility complex 2.

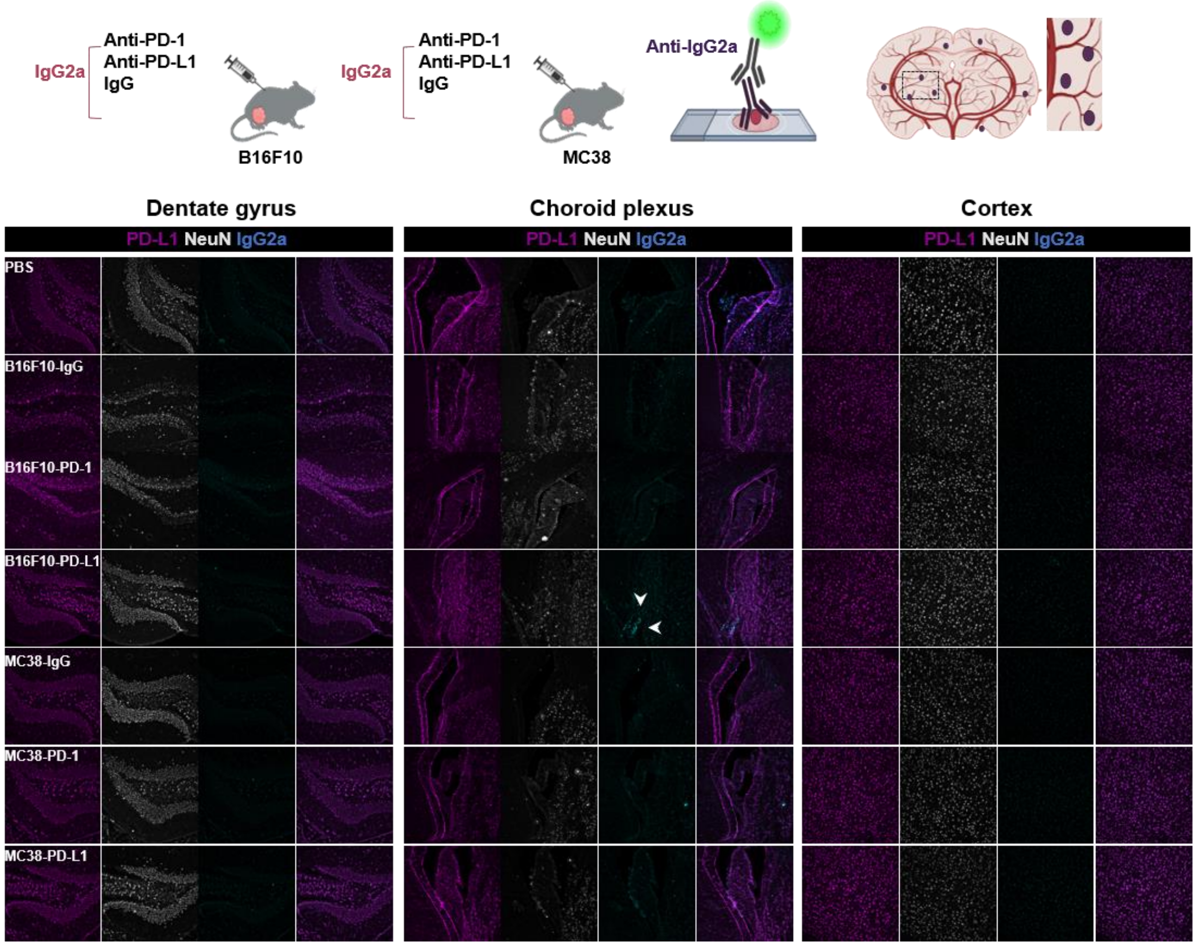


**Figure S8. Detection of IgG2a into brain areas from B16F10- and MC38-bearing mice treated with Anti-PD1, Anti-PD-L1 or IgG.** Timeline representation of IgG2a detection by immunofluorescence after injection of Anti-PD1, Anti-PD-L1or IgG in B16F10- and MC38-bearing mice. Representative images of PD-L1 (magenta), NeuN (grey) neurons and IgG2a (cyan) immunoreactivities in dentate gyrus (left panel), choroid plexus (middle panel) and cortex (right panel) of PBS-treated mice and B16F10 or MC38-bearing mice treated with Anti-PD1, Anti-PD-L1 or IgG. White arrows highlight IgG2a detection in PD-L1+ cells in choroid plexus. Scale bar: 100 µm. IgG: immunoglobulin G, NeuN: neuronal nuclei antigen, PD-1: programmed cell death 1, PD-L1: programmed cell death ligand 1.

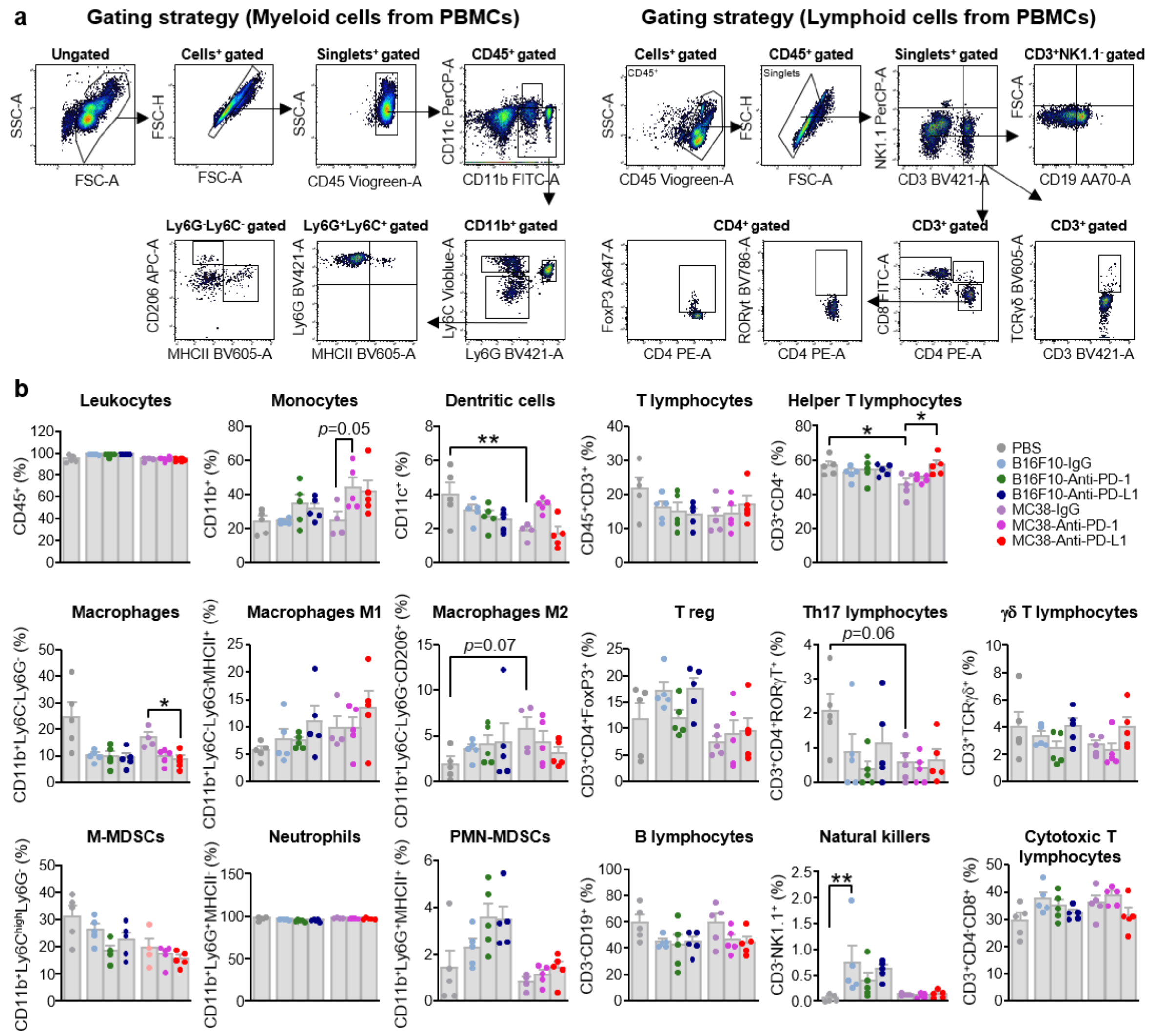


**Figure S9. Impact of anti-PD1 and anti-PD-L1 on peripheral immune cells and brain infiltrating TCRγδ cells. a.** Gating strategy for the analysis of peripheral blood immune cells by flow cytometry from B16F10- and MC38-bearing mice treated with Anti-PD1, Anti-PD-L1or IgG. For myeloid panel, live cells were gated from FSC-A and SSC-A, singlets were gated from FSC-A and FSC-H, CD45 was used to identify leucocytes (CD45$^{+}$). From these cells, CD11c and CD11b were used to identify dendritic cells (CD11c$^{+}$CD11b$^{+}$) and monocytes (CD11c$^{-}$CD11b$^{+}$). From CD11b$^{+}$ cells, Ly6C, Ly6G and MHCII were used to identify macrophages (CD11b$^{+}$Ly6C$^{-}$Ly6G$^{-}$), neutrophils (CD11b$^{+}$Ly6G$^{+}$Ly6c$^{-}$MHCII$^{-}$), pro-tumoral macrophages (CD11b$^{+}$Ly6C$^{-}$Ly6G-CD206+), anti-tumoral macrophages (CD11b$^{+}$Ly6C$^{-}$Ly6G$^{-}$MHCII$^{+}$), M-MDSCs (CD11b$^{+}$Ly6G$^{-}$Ly6c$^{high}$), PMN-MDSCs (CD11b$^{+}$Ly6G$^{+}$Ly6C$^{low}$MHCII$^{+}$). For lymphocytic panel, Live cells were gated from FSC-A and SSC-A, singlets were gated from FSC-A and FSC-H, CD45 was used to identify leucocytes (CD45$^{+}$). From these cells, CD3 and NK1.1 were used to identify lymphocytes (CD3$^{+}$NK1.1$^{-}$) and NK cells (CD3$^{-}$NK1.1$^{+}$). From CD3$^{+}$NK1.1$^{-}$ cells, CD4 and CD8 were used to identify CD4 T cells (CD4$^{+}$CD8$^{-}$) and CD8 T cells (CD8$^{+}$CD4$^{-}$) and CD19 was used to identify B cells (CD19$^{+}$). From CD3$^{+}$ cells, TCRγδ cells were identify by using TCRγδ (CD3$^{+}$TCRγδ$^{+}$). From CD4$^{+}$ cells, FoxP3 and RORγT were used to identify Treg cells (FoxP3$^{+}$CD3$^{+}$CD4$^{+}$) and Th17 cells (CD3$^{+}$CD4$^{+}$RORγT$^{+}$). **b.** Histograms representing variation in percentage of myeloid (left) and lymphocytic (right) immune cells population from whole blood collected from B16F10 or

MC38-bearing mice treated with Anti-PD1, Anti-PD-L1or IgG. Statistical analysis was performed by using one way ANOVA or Kruskal-Wallis test with Bonferroni or Dunn's correction for multiple comparisons (F10-IgG vs F10-PD1 or F10-PD-L1, MC38IgG vs MC38-PD1 or MC38-PD-L1). Data are represented as bars with symbols for individual data points and they are expressed by mean ± SEM, n=5, * $p<0.05$, ** $p<0.01$. **c.** Representative images of CD3 (magenta), TCR γδ (green), Lectin (grey) and DAPI (blue) immunoreactivities in hippocampus of B16F10 or MC38-bearing mice treated with Anti-PD1, Anti-PD-L1or IgG. Scale bars: 50 µm, zoom 10 µm. CD: cluster of differentiation, CD11b: integrin alpha M subunit, CD11c: complement component 3 receptor 4 subunit, CD206: cluster of differentiation 206, CD45: leukocyte common antigen, DAPI: 4',6-diamidino-2-phenylindol, FoxP3: forkhead box P3, FSC: forward scattering gating, IgG: immunoglobulin G, Ly6C: lymphocyte antigen 6 family member C, Ly6G: lymphocyte antigen 6 family member G, MHCII: major histocompatibility complex 2, M-MDSC: monocytic-derived myeloid derived suppressor cells, NeuN: neuronal nuclei antigen, NK: natural killer, PD-1: programmed cell death 1, PD-L1: programmed cell death ligand 1, PMN-MDSC: polymorphonuclear myeloid derived suppressor cells, ROR: Retinoic acid-related Orphan Receptors, SSC: side scattering gating, TCR: T cell receptor.

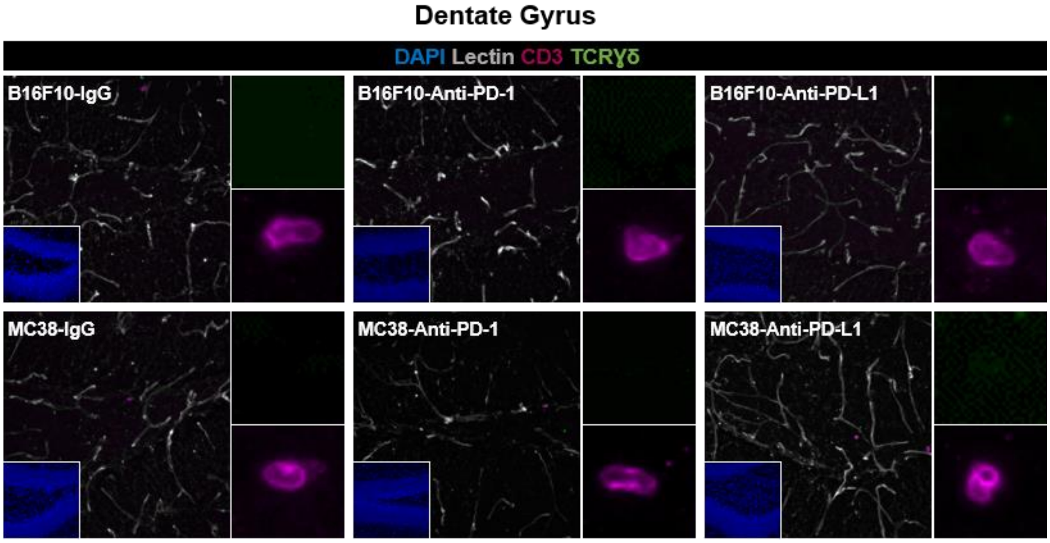


**Figure S10. T lymphocytes in infiltration in the hippocampus of cancer-bearing mice.** Representative images of CD3 (magenta), TCRγδ (green), Lectin (grey) and DAPI (blue) immunoreactivities in hippocampus of B16F10 or MC38-bearing mice treated with Anti-PD1, Anti-PD-L1or IgG. Scale bars: 50 µm, zoom 10 µm. CD: cluster of differentiation, DAPI: 4',6-diamidino-2-phenylindol, NeuN: neuronal nuclei antigen, PD-L1: programmed cell death ligand 1, TCR: T cell receptor.

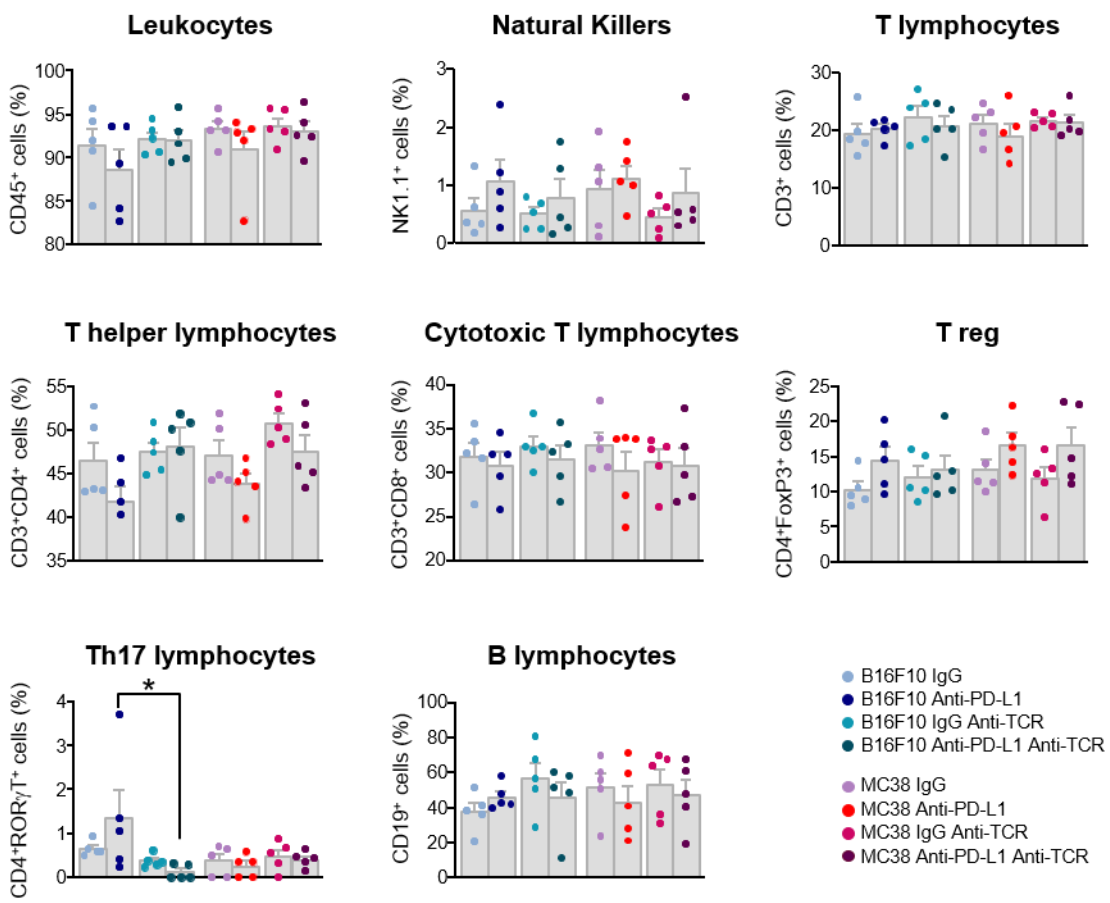


**Figure S11. Impact of anti-TCRγδ on circulating immune cells of cancer-bearing mice treated or not with anti-PD-L1.** Impact of Anti-TCRγδ administration on proportion of circulating $CD45^+$ leucocytes, CD3-$NK1.1^+$ NK cells, $CD3^+$ lymphocytes, $CD3^+CD4^+$ CD4 T cells, $CD3^+CD8^+$ CD8 T cells, $CD3^+CD4^+FoxP3^+$ T helper cells, double positive $CD4^+CD8^+$ T cells, $CD19^+$ B cells and $CD3^+CD4^+ROR\gamma T^+$ Th17 cells in blood of B16F10- and MC38-bearing mice treated with Anti-PD-L1 and IgG. Statistical analyses were performed using one-way ANOVA or Kruskal-Wallis test with Bonferroni or Dunn's correction for multiple comparisons (B16F10 IgG vs. B16F10-PD-L1, B16F10 IgG-TCRγδ vs. B16F10-PD-L1-TCRγδ, B16F10 IgG vs. B16F10 IgG- TCRγδ, B16F10-PD-L1 vs. B16F10-PD-L1-TCRγδ). Data are represented as bars with symbols for individual data points and they are expressed by mean ± SEM, n=5. CD: cluster of differentiation, CD45: leukocyte common antigen, FoxP3: forkhead box P3, IgG: immunoglobulin G, NK: natural killer, PD-L1: programmed cell death ligand 1, ROR: Retinoic acid-related Orphan Receptors, TCR: T cell receptor.

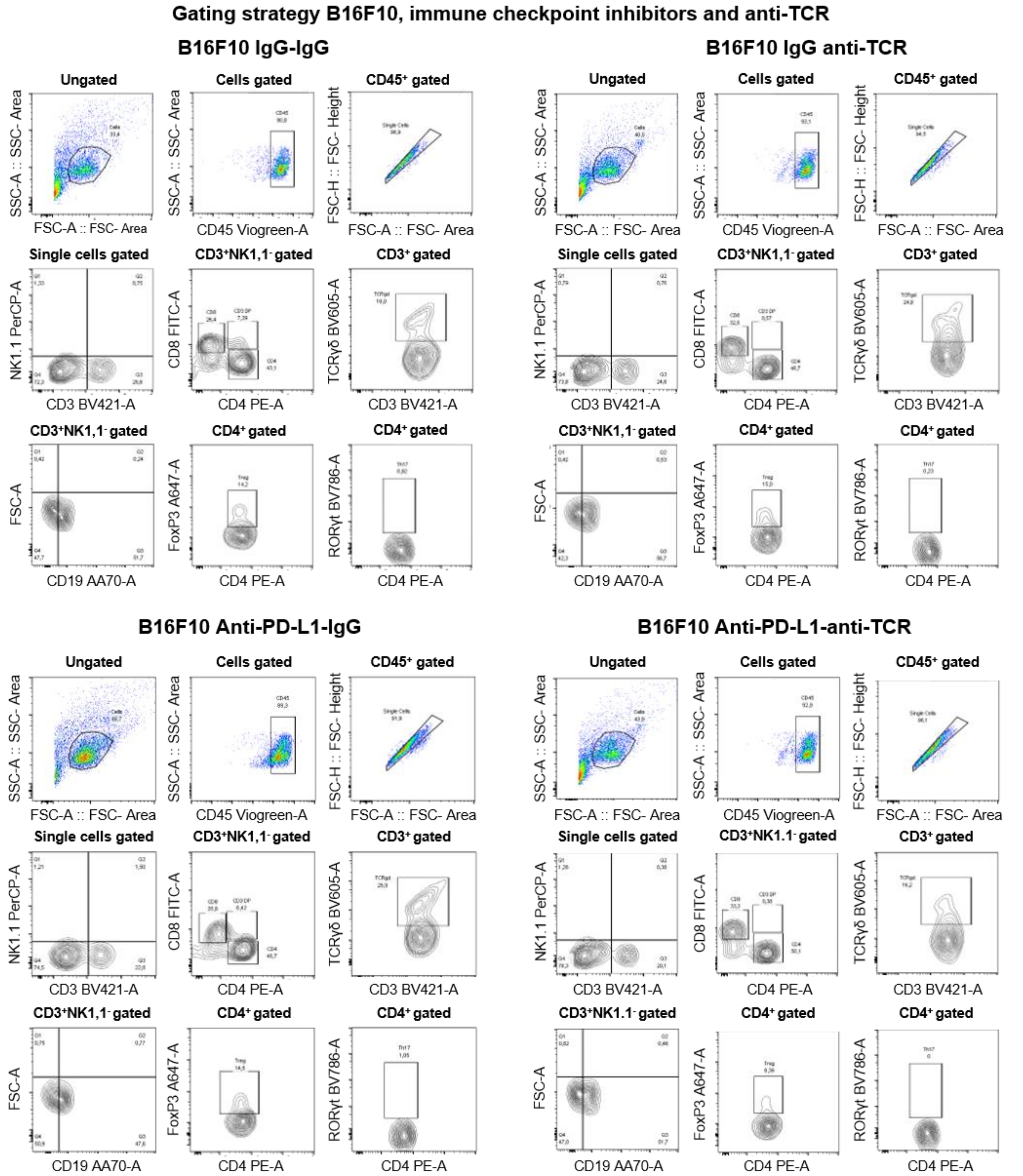


**Figure S12. Gating strategy for the analysis of peripheral blood cells by flow cytometry from B16F10-bearing mice treated or not with anti-PD-L1 and anti-TCRγδ.** Live cells were gated from FSC-A and SSC-A, singlets were gated from FSC-A and FSC-H, CD45 was used to identify leucocytes ($CD45^+$). From these cells, CD3 and NK1.1 were used to identify lymphocytes ($CD3^+NK1.1^-$) and NK cells ($CD3^-$, $NK1.1^+$). From $CD3^+NK1.1^-$ cells, CD4 and CD8 were used to identify CD4 T cells ($CD4^+CD8^-$) and CD8 T cells ($CD8^+CD4^-$) and CD19 was used to identify B cells ($CD19^+$). From $CD3^+$ cells, TCRγδ cells were identify by using TCRγδ ($CD3^+TCR\gamma\delta^+$). From $CD4^+$ cells, FoxP3 and RORγT were used to identify Treg cells ($FoxP3^+CD3^+CD4^+$) and Th17 cells ($CD3^+CD4^+ROR\gamma T^+$). CD: cluster of differentiation, CD45: leukocyte common antigen, FoxP3: forkhead box P3, FSC: forward scattering gating,

IgG: immunoglobulin G NK: natural killer, PD-L1: programmed cell death ligand 1, ROR: Retinoic acid-related Orphan Receptors, SSC: side scattering gating, TCR: T cell receptor.

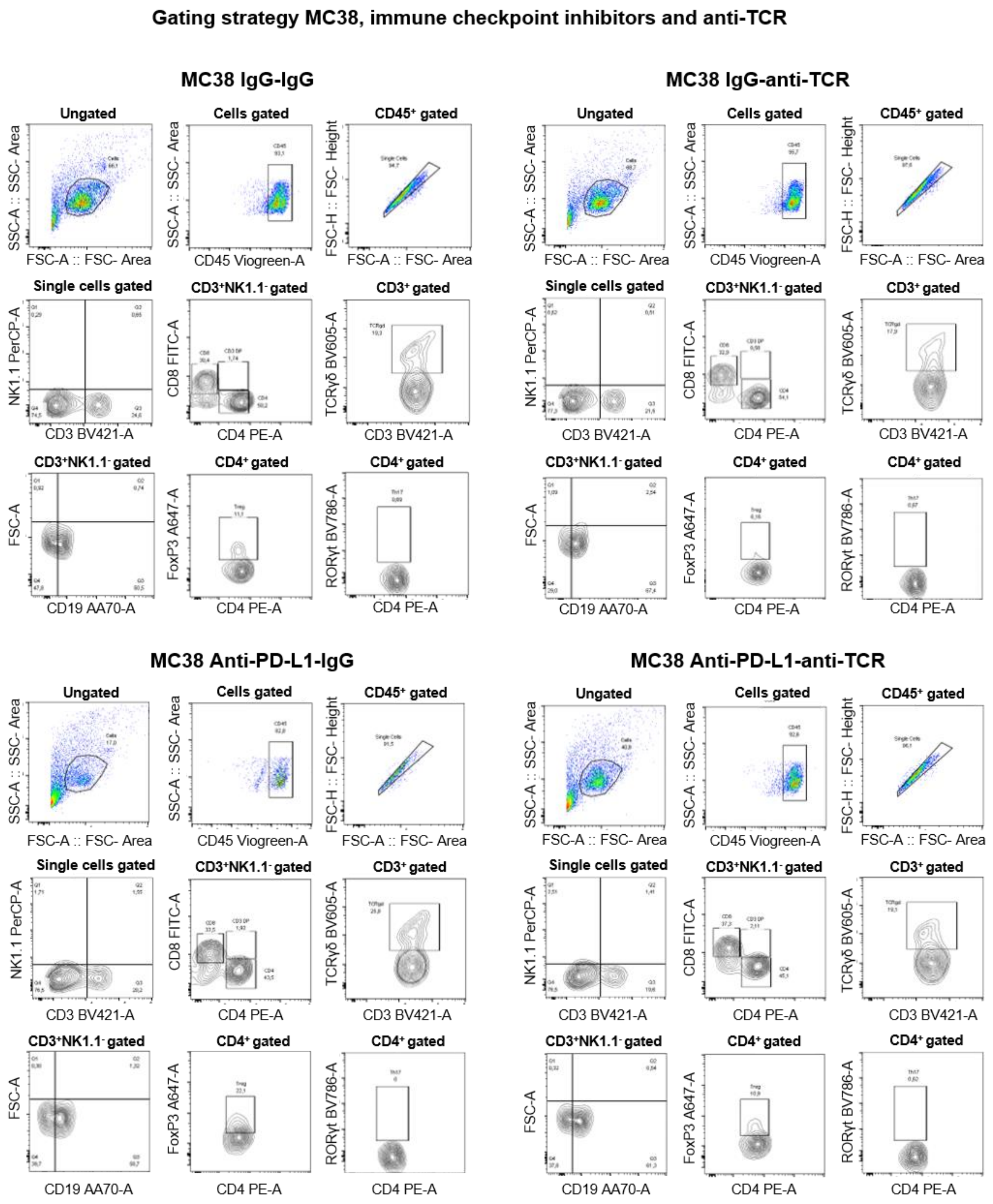


**Figure S13. Gating strategy for the analysis of peripheral blood cells by flow cytometry from MC38-bearing mice treated or not with anti-PD-L1 and anti-TCRγδ.** Live cells were gated from FSC-A and SSC-A, singlets were gated from FSC-A and FSC-H, CD45 was used to identify leucocytes ($CD45^+$). From these cells, CD3 and NK1.1 were used to identify lymphocytes ($CD3^+NK1.1^-$) and NK cells ($CD3^-$, $NK1.1^+$). From $CD3^+NK1.1^-$ cells, CD4 and CD8 were used to identify CD4 T cells ($CD4^+$, $CD8^-$) and CD8 T cells ($CD8^+$, $CD4^-$) and CD19 was used to identify B cells ($CD19^+$). From $CD3^+$ cells, TCRγδ cells were identify by using TCRγδ ($CD3^+TCR\gamma\delta^+$). From $CD4^+$ cells, FoxP3 and RORγT were used to identify Treg cells ($FoxP3^+CD3^+CD4^+$) and Th17 cells ($CD3^+CD4^+ROR\gamma T^+$). CD: cluster of differentiation,

CD45: leukocyte common antigen, FoxP3: forkhead box P3, FSC: forward scattering gating, IgG: immunoglobulin G NK: natural killer, PD-L1: programmed cell death ligand 1, ROR: Retinoic acid-related Orphan Receptors, SSC: side scattering gating, TCR: T cell receptor.
